# The cell cycle variant in multiciliated cells incorporates 2 centriole biogenesis cycles

**DOI:** 10.1101/2024.02.09.579615

**Authors:** Amélie-Rose Boudjema, Rémi Balagué, Cayla E Jewett, Jacques Serizay, Gina M LoMastro, Olivier Mercey, Adel Al Jord, Adrien Candat, Marion Faucourt, Alexandre Schaeffer, Camille Noûs, Nathalie Delgehyr, Andrew J Holland, Nathalie Spassky, Alice Meunier

**Author notes:** These authors contributed equally to this work (Amélie-Rose Boudjema, Rémi Balagué).

## Abstract

Multiciliated cell (MCC) differentiation is a calibrated version of the canonical cell cycle. The MCC cell cycle variant sustains amplification of centrioles for the nucleation of dozens of motile cilia, while avoiding cell division. In this study, we show that the MCC cell cycle variant is an accelerated version of the canonical cell cycle, which superposes two cycles of centriole biogenesis, in order to obtain multiple mature centrioles within a single -instead of double- cell cycle iteration. We further show that the precocious maturation of amplified procentrioles is even determinant for their spatial self-organization, disengagement and apical migration for cilia nucleation. Our findings collectively suggest that the decomposition of centriole biogenesis over two cycle iterations in dividing cells may have been adopted to ensure the growth of a solitary primary cilium, and exemplify how minimal deviations of the canonical cell cycle allow MCC progenitors to both amplify, and accelerate, centriole biogenesis for vital motile ciliogenesis.

## Introduction

Centriole amplification is a characteristic of multiciliated cell (MCC) differentiation, necessary for the nucleation of dozens of motile cilia that propel physiological fluids in the brain, the respiratory and the reproductive tracts. However, it is a pathological mechanism in cancer cells, where it can cause genomic instability that accelerates tumor formation (Nigg & Holland, 2018). It is interesting to note that centriole amplification dynamics in post-mitotic MCCs show similarities with the highly conserved process of centriole duplication that occurs in parallel with DNA during cell division (Al Jord et al., 2014; Herawati et al., 2016; Ma et al., 2014; Revinski et al., 2018; Stubbs et al., 2012; Tan et al., 2013; Vladar & Stearns, 2007; Zhao et al., 2013) . Consistently, MCC differentiation is driven by a true variant of the cell cycle, co-opting the mitotic oscillator and transcriptional topology of the cell cycle, but changing the cyclins, E2F transcription factor and some regulators of APC/C to hijack cell cycle activity and skip DNA duplication while (over-) replicating centrioles (Al Jord et al., 2017, 2019; Choksi et al., 2024; Khoury Damaa et al., 2025; Kim et al., 2022; Serizay et al., 2025; Vladar et al., 2018).

Three highly stereotyped sequential stages have been resolved by live imaging mouse brain MCCs: the amplification stage (called A-stage), during which dozens of latent procentrioles form sequentially around centrosomal centrioles and MCC specific deuterosome organelles; the growth stage (called G-stage), during which all procentrioles elongate synchronously; the disengagement stage (called D-stage), during which all centrioles detach from their growing platforms in a synchronized manner to finally migrate towards the apical membrane and nucleate motile cilia (Al Jord et al., 2014). Transitions between these stages, which share similarities with the S-, G2- and M-phase of the canonical cell cycle, are controlled by the mitotic oscillator. Specific inhibition or disinhibition of core components of this cell cycle clock-like regulatory circuit (CDK1, PLK1 and APC/C) results in altering centriole number, growth and disengagement, in a manner similar to centriole duplication (Al Jord et al., 2017; Kim et al., 2022; Revinski et al., 2018).

While the dynamics and molecular cascade are mostly comparable between centriole biogenesis in cycling and MCC differentiating cells, two major differences exist. The first one is the number of centrioles being produced : a dividing MCC progenitor produces one procentriole per parental centriole during the canonical cell cycle, but when it differentiates, the MCC progenitor cell enters a final modified iteration of the cell cycle that results in the production of several dozen centrioles. The second difference is the pace of centriole production : it takes almost two iterations of the canonical cell cycle to produce a mature « mother » centriole -or « basal body »- able to nucleate microtubules (MT) and grow a cilium, but a single iteration of the MCC cell cycle variant is needed to produce basal bodies, able to nucleate cytoplasmic MT and grow cilia.

Here, we used micro-patterned MCCs, new knock-in transgenic mice, ultrastructure expansion microscopy (U-ExM), correlative light and electron microscopy and MT/dynein pharmacological inhibitions, to explore these two specificities of the MCC cell cycle variant. In space, we found that centriole amplification emerges in a pericentrosomal « nest » concentrating core centriole/deuterosome elements, and that both the nest and the production of centrioles/deuterosomes are altered when MT or dynein are inhibited. In time, we found that the MCC cell cycle accelerates the centriole biogenesis program by superposing 2 canonical centriole cycles, leading to the concomitant elongation and maturation of procentrioles in a single cycle iteration. In this 2-in-1 cycle, the precocious maturation of procentrioles is moreover determinant for their subsequent spatial self-organization, disengagement and apical migration.

## Results

### MEF-MCCs grown on micropatterns localize the onset of centriole amplification at the pericentrosomal region

The origin of amplified centrioles in MCC remains controversial. Some live imaging experiments and electron microscopy suggest that the centrosome could constitute a nest for centriole and deuterosome biogenesis (Al Jord et al., 2014; Kalnins et al., 1972; Mori et al., 2017), but others have proposed that procentriole-loaded deuterosomes emerge independently from the centrosome location, all over the cytoplasm (Nanjundappa et al., 2019; Sorokin, 1968; Zhao et al., 2013, 2019). Interestingly, in the absence of deuterosome, procentrioles are massively amplified from and around centrosomal centrioles, yet, centrosomal centrioles are dispensable (Mercey et al., 2019a; Mercey et al., 2019b; Nanjundappa et al., 2019; Zhao et al., 2019). Finally, in the absence of both centrosomes and deuterosomes, procentrioles are seen emerging from the middle of a self-organized microtubule organizing center (MTOC) suggesting that such MTOC could organize procentriole biogenesis (Mercey et al., 2019).

To further assess whether procentrioles are emerging from a particular location in MCCs, we took advantage of a cellular model based on the induced differentiation of MEFs into MCCs by ectopic adenoviral expression of MCIDAS, E2F4 and E2F4 transactivator Dp1 (Kim et al., 2018). This model of MCCs is advantageous because it is not tissue-specific and does not require epithelial cell-to-cell contact for differentiation. This allowed us to trigger MCC differentiatiation in single cells platted on micropatterns to impose a reproducible cytoplasmic organization facilitating spatial quantifications. We induced MCC differentiation in MEFs grown on crossbow-shaped micropatterns to analyze the global spatial dynamics of procentriole amplification (Fig. 1A). We first confirmed that, in MEF-MCCs, the same dynamics of amplification was observed as in brain MCC: an “amplification stage (A-stage)”, marked by the progressive accumulation of immature non-polyglutamylated (GT335 “-”) procentrioles is followed by a “growth stage (G-stage)” where procentrioles mature and become polyglutamylated (GT335 “+”). This maturation is followed by ciliation (MCC-stage) which is often partial, probably because of the absence of the typical epithelial apico-basal polarity (Fig. 1B). Consistently, in this cellular model, centrosomes can localize below the nucleus. Here, only cells with apically positioned centrosome were studied. By density mapping the location of centrosomes in multiple crossbow-shaped MEFs before MCC differentiation (precursor stage), we confirmed that centrosomes exhibit a reproducible position and are located at the center of convergence of microtubules (Fig. 1C, top row left) colocalized with PERICENTRIN (PCNT, Fig 1D, top row left). Then, by density mapping the position of SAS6+ procentrioles in differentiating MEF-MCCs containing less than 10 SAS6 foci (early A-stage), we observed that early procentriole emergence is localized in the centrosomal region, where PCNT also accumulates (Fig. 1C, 1D top row middle). Later during amplification (>10 SAS6+ foci, late A-stage and G-stage), procentrioles and PCNT occupy a larger portion of the cells centered around the centrosome location and surrounding the nuclear region (Fig. 1C-D top row right). These results suggest that, in this cellular model, procentrioles emerge in the centrosomal region.

**Figure 1:**
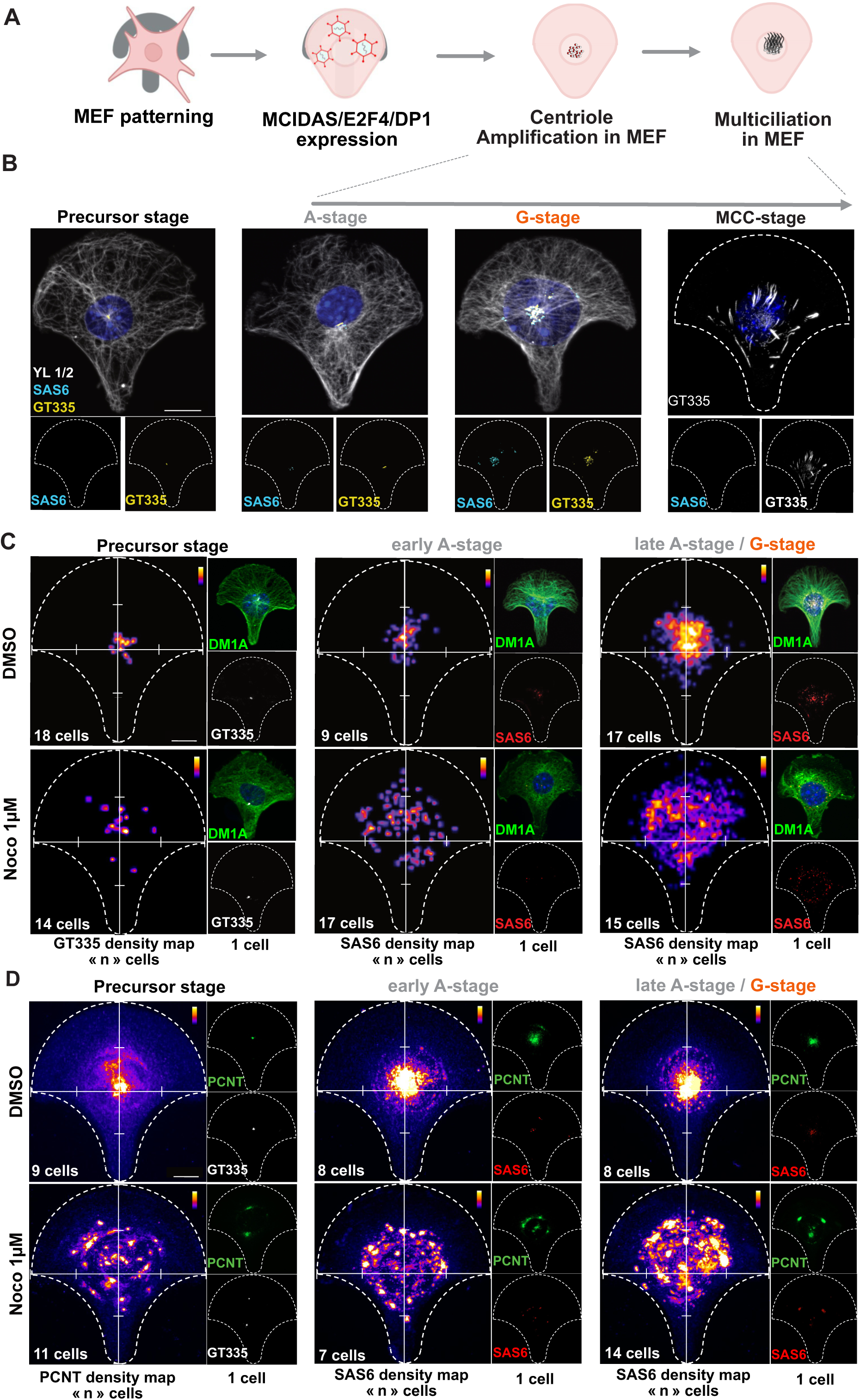
MEF-MCCs grown on micropatterns highlight the pericentrosomal focalization of procentriole production. **(A)** Scheme of the MEF-MCC differentiation process on crossbow micropatterns. **(B)** MCC-induced fibroblasts on micropatterns perform the typical step-wise dynamics of centriole amplification. Representative immunofluorescence images of MEF-MCC on crossbow micropatterns showing the 4 stages of differentiation. Scale bar, 10 µM. **(C)** Density mapping of centrosome in MEF-MCCs before amplification (precursor stage, left column), and of procentrioles during early amplification (cells with less than 10 SAS6 focis/cell, middle column) or late amplification (cells with more than 10 SAS6 focis/cell, right column) treated with DMSO (top row) or Nocodazole 1 µM (bottom row). Three independent experiments were quantified. Scale bar, 10 µM. **(D)** Density mapping of PCNT fluorescence signal during centrosome stage (left column) early A-stage (middle column) and late A-stage (right column) in MEF-MCCs under DMSO (top row) or Nocodazole 1 µM (bottom row). Two independent experiments were quantified. Scale bar, 10 µM. DM1A marks all MT, YL1/2 marks tyrosinated MT, SAS6 marks procentrioles, GT335 marks centrosome, mature procentrioles and cilia, PCNT marks PERICENTRIN. Dotted lines show the crossbow shape of the micropattern, white graduated lines show x and y axes. Insets on the right of each panel on (C) and (D) show one representative cell used for the density mapping.

We previously showed in brain MCC deprived of both centrosomes and deuterosomes, that centrioles emerged from the center of a self-organized microtubule network, within a PCNT cloud (Mercey, Levine, et al., 2019). To test whether microtubules drive the organization of a centrosomal nest from which procentrioles emerge, we treated MEF-MCCs with a dose of nocodazole affecting mildly the microtubule network (Fig. 1C, bottom row), before the onset of centriole amplification (see methods). In this condition, SAS6+ structure density mapping shows a clear dispersion of procentrioles compared to control cells (Fig. 1C bottom row). The centrosomal PCNT cloud also scatters in the cell cytoplasm (Fig. 1D bottom row). Since MT perturbation also induced centrosomal centriole splitting and decentralization of centrosomal centrioles (Fig. 1C, bottom row left), we also measured the distance between emerging SAS6+ foci and the closest centrosomal centriole. This confirmed that procentriole-to-centrosome distance increases with nocodazole (Fig. 1 Supplementary 1A-B).

Altogether, these results suggest that, in this non tissue specific proxy of MCC progenitors, MT organize the onset of centriole amplification in the pericentrosomal region.

### Correlative DEUP1 live-imaging and EM highlights the existence of a pericentrosomal “nest” in brain MCC

We then sought to study early onset of centriole biogenesis in physiological MCCs. Live monitoring centriole amplification in MCCs was only reported in brain cultured progenitors since respiratory and reproductive MCCs are integrated in stratified epithelia, hindering their live observation by inverted high-resolution live microscopy. It was done using GFP-tagged CENTRIN2 (CEN2-GFP) in control cells and in cells deprived of centrosome and/or deuterosomes through Centrinone treatment and/or DEUP1 gene invalidation (DEUP1 KO). These studies revealed that procentrioles, in presence (WT) or absence of deuterosomes (DEUP1 KO), where emerging from a pericentrosomal cloud of CEN2-GFP, suggesting that they were formed around the centrosome (Al Jord et al., 2014; Mercey, Al Jord, et al., 2019; Mercey, Levine, et al., 2019) . Further studies overexpressing a tagged version of DEUP1, contested these results by showing that DEUP1+ structures were appearing scattered in the cytoplasm (Zhao et al., 2019). However, these observations were done through DEUP1 overexpression, which induces the formation of non physiological DEUP1 aggregates, and in the presence of nocodazole, which can disturb the dynamics (Fig. 1C-D).

In order to catch the very fist events of centriole amplification, we developed a transgenic mouse line expressing CEN2-GFP, together with the core deuterosome component DEUP1, endogenously tagged with a mRuby fluorescent reporter (mRuby-DEUP1; see methods). Imaging brain MCC progenitors from these mice confirmed the absence of non physiological DEUP1 aggregates and the accurate monitoring of mRuby-DEUP1+ deuterosomes in addition to the CEN2-GFP+ procentrioles during the A-, G- and D-stages (Fig. 2A, Fig. 2 Supplementary 1A, (Al Jord et al., 2014, 2017)). To highlight the very early onset of amplification we live imaged early cultures and revealed that centriole amplification A-stage begins with the formation of a cloud of DEUP1, which we called “primordial” cloud (Fig. 2B, red arrowhead, 05:50 to 12:30; Video 1). This cloud forms and grows around the centrosome (Fig. 2B, white arrowheads, n>30 cells). Over time, DEUP1+ foci, which are also CEN2-GFP+, emerge from this cloud (Fig 2B, red and white arrowheads, 15:50-18:20). They represent procentriole-loaded deuterosomes characterizing the A-stage previously described using CEN2-GFP only (Al Jord et al., 2014).

**Figure 2:**
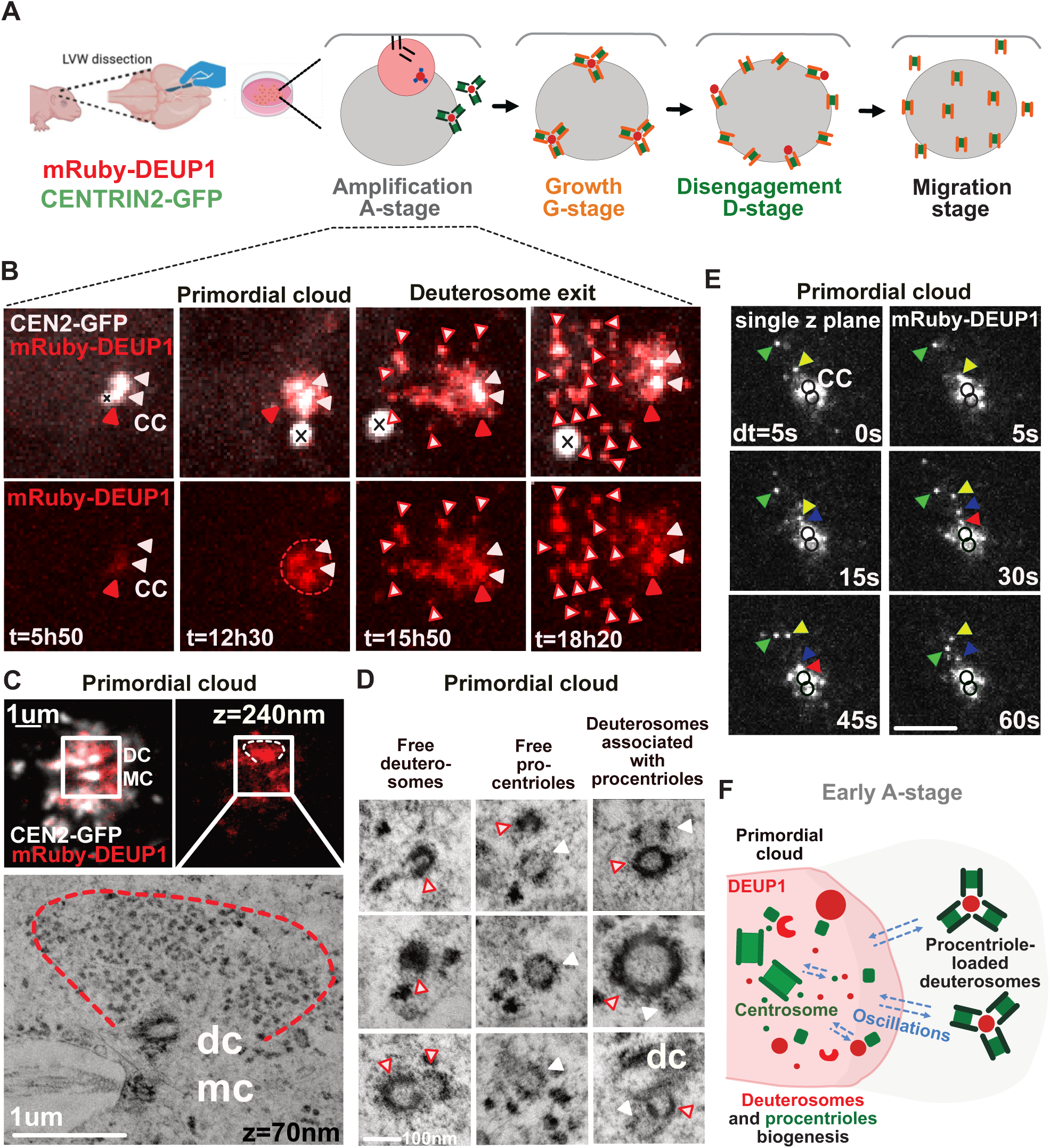
Live imaging endogenous DEUP1 highlights the first events of centriole amplification in brain MCC. **(A)** Scheme of the ependymal cell culture protocol. LVW: lateral ventricular wall. **(B)** Early A-stage dynamics in CEN2-GFP; mRuby-DEUP1+ cells. White arrowheads point the 2 centrosomal centrioles. The red dotted line corresponds to the cloud of DEUP1+ forming around the parental centrioles during early A-stage. The red arrowhead points to DEUP1 accumulation at one centrosomal centriole as compared to the other one. The red and white arrowheads point to CEN2-GFP+ structures corresponding to centriole loaded deuterosomes. The crosses rule out the CEN2-GFP+ aggregate not to be mistaken with a centrosome; CEN2-GFP in white, DEUP1 in red; scale bar, 5 µM. See associated Video 1. **(C)** Correlative light and electron microscopy resolves fibrogranular aggregate concentration in a mRuby-DEUP1+ primordial cloud. Light microscopy integrates the fluorescence signal of a z-stack of 240 nm. Electron microscopy z-sections are 70 nm. dc= centrosomal daughter centriole, mc= centrosomal mother centriole. **(D)** Representative ultrastructural elements present in the mRuby-DEUP1 primordial cloud resolved by correlative light and electron microscopy. Red and white arrowheads indicate deuterosome structures. White arrowheads indicate procentrioles. dc= centrosomal daughter centriole. See also Video 3. **(E)** Representative images of live imaged mRuby-DEUP1+ foci fluctuations in the primordial cloud (dt= 5s). A single z-slice of 0,7 µM is filmed. Black circles indicate the position of centrosomal centrioles located thanks to the CEN2-GFP signal, not shown here. mRuby-DEUP1+ foci are color-coded with arrowheads to appreciate their oscillatory behaviors towards the centrosomal mRuby-DEUP1+cloud. See associated video 4. Scale bar, 5 µM. **(F)** Scheme depicting the observations resulting from live-imaging CEN2-GFP; mRuby-DEUP1+ cells and correlative light and electron microscopy.

To further assess the composition of the primordial cloud from which procentriole-loaded deuterosome emerge, we increased spatial resolution and identified the cloud as diffuse and/or composed of intermingled mRuby-DEUP1+ and CEN2-GFP+ foci of different sizes (Fig. 2 Supplementary 1B), which do not necessarily colocalize and can move independently from each others (Fig. 2 Supplementary 1B, video 2). We then used correlative light and electron microscopy (CLEM) to reveal the ultrastructure of the cloud and found that the mRuby-DEUP1+ diffuse signals outline regions rich in fibrogranular material, previously characterized as centriolar satellites (Zhao et al., 2021). This region was found to be either deprived of deuterosomes (Fig. 2C, Fig. 2 Supplementary 2) or embedding small nascent deuterosomes taking the form of amorphous electron dense structures, horseshoes or complete spheres (Fig. 2D, Fig. 2 Supplementary 3-6, video 3). As previously described, small deuterosomes can be seen connected to the centrosomal daughter centriole (“dc”, Fig. 2D, bottom right; Fig. 2 Supplementary 3, 4, video 3, (Al Jord et al., 2014; Khoury Damaa et al., 2025)). Nascent deuterosomes are either free (Fig. 2D, Fig. 2 Supplementary 3-4, blue stars), or loaded with small procentrioles (Fig. 2D, Fig. 2 Supplementary 3-5, black stars). Procentrioles seemingly not loaded on deuterosome can sometimes be detected (Fig. 2D, Fig. 2 Supplementary 4, red arrows; video 3). At this early stage, correlation between deuterosome size and the centriole number they are associated with is not evident: CLEM shows that even big deuterosomes are associated with only one or two procentrioles (Fig. 2D; Fig. 2 Supplementary 6). Live monitoring mRuby-DEUP1 further shows that the size of these young deuterosomes homogenizes with time. This is achieved through the splitting of big deuterosomes into smaller ones, when big deuterosomes are present (Fig. 2 Supplementary 1C), and through either fusion (never observed live) or growth, when amplification begins with small deuterosomes.

To better resolve how deuterosome and centriolar material concentrate around the centrosome, we then increased the temporal resolution (dt=5-15s for 1-4min) and revealed stochastic back and forth fluctuations to the centrosome for both early CEN2-GFP-/mRuby-DEUP1+ or CEN2-GFP+/mRuby DEUP1- structures (Fig. 2E, Video 4-5, 5/10 cells). These stochastic fluctuations are also visible later on CEN2-GFP+/mRuby-DEUP1+ deuterosomes (Video 6, Fig. 2 Supplementary 1D, 13/15 cells, white arrowheads). This movement to the centrosome is reminiscent to the dynamics of the centriolar satellite protein PCM1 in *Xenopus* and mice, and the PCM protein Centrosomin in *Drosophila* (Hall et al., 2023; Kubo et al., 1999; Megraw et al., 2002), suggesting that centriole and deuterosome components are connected to the centrosome, as other pericentrosomal components during A-stage. Interestingly, back and forth movements of mRuby-DEUP1+ structures to one centrosomal centriole are frequently observed in live (Video 6, 7, Fig. 2 Supplementary 1E) and may explain the observed connection of deuterosomes to the centrosomal daughter centriole on fixed samples during A-stage (this paper, (Boudjema et al., 2024; Khoury Damaa et al., 2025)),

Altogether live imaging mRuby-DEUP1/CEN2-GFP during early A-stage suggests that core deuterosome and centriole components are concentrated in a primordial cloud around the centrosome, which constitutes a nest where centrioles and deuterosomes concomitantly form before they move away from the centrosomal region (Fig. 2F). Back and forth fluctuations of CEN2-GFP+ and mRuby-DEUP1+ structures during A-stage suggest the existence of centrosome-related forces avoiding dispersion of primordial structures away from the centrosomal region.

### S-phase like steps are recapitulated in the pericentrosomal nest

To gain molecular resolution on the early events occurring within the pericentrosomal nest, we performed ultrastructure expansion microscopy (U-ExM). U-ExM is an extension of expansion microscopy that allows the visualization of structures by optical microscopy with 4-fold increased resolution (Gambarotto et al., 2019).

U-ExM analysis confirmed the existence of a pericentrosomal cloud characterized by live imaging. In these fixed detergent-treated cells though, the cloud is around 10-fold dimmer than the centrosomal centrioles, and thus difficult to visualize without over-saturating the signal, explaining why it was not described before (Fig. 3 Supplementary 1A-E). In the primordial cloud, U-ExM identifies DEUP1 foci at and around the parent centrioles, with a light bias to the daughter centriole (Fig. 3A, H; Fig. 3 Supplementary 1C-E). This cloud of DEUP1 and CENTRIN also includes PCNT (Fig. 3 Supplementary 1F). Notably, CENTRIN and DEUP1 foci are often in close proximity but not necessarily colocalized, confirming live imaging, while PCNT puncta partially overlap with CENTRIN (Fig. 3 Supplementary 1F). At this stage, PLK4, the master regulatory kinase, and SAS6, one of the first centriolar components are either absent, or present as small foci within the cloud, often on the wall of the parent centrioles, also with a bias to the daughter centriole (Fig. 3A-B).

**Figure 3:**
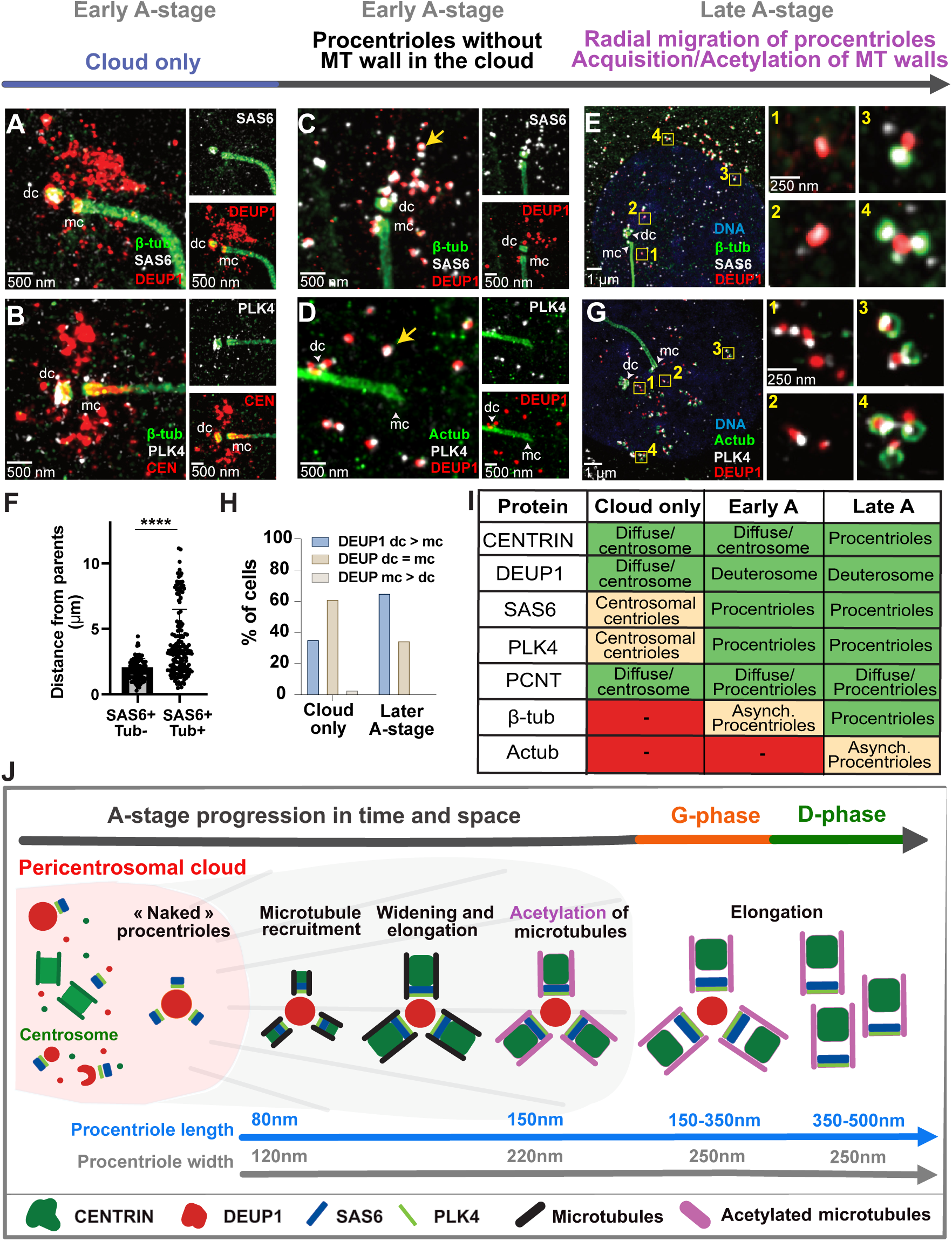
Procentriole assembly begins in a pericentrosomal nest and progresses during radial migration. **(A-B)** Representative U-ExM images of brain MCCs during early A-stage, when only a primordial cloud is visible but no procentriole-loaded deuterosomes can be identified. Cells were immunostained with antibodies to β-TUBULIN, SAS6, DEUP1 (A), β-TUBULIN, CENTRIN, PLK4 (B). mc: mother centriole, dc: daughter centriole. **(C-D)** Representative U-ExM images of brain MCCs during early A-stage, when procentriole-loaded deuterosomes are visible. Cells were immunostained with antibodies to β-TUBULIN, SAS6, DEUP1 (C) or Acetylated-TUBULIN, PLK4, DEUP1 (D). Arrows mark DEUP1 foci with either SAS6 (C) or PLK4 (D). mc: mother centriole, dc: daughter centriole. **(E)** Representative U-ExM images of brain MCCs during late A-stage. Boxes denote zoomed in regions on right, with 1-4 labels marking increasing distance from the parent centrioles. Cells were immunostained with antibodies to β-TUBULIN, SAS6, DEUP1 (E) **(F)** Quantitation showing the procentriole distance from parents is shorter in procentrioles without αβ-TUBULIN compared to procentrioles with αβ-TUBULIN. SAS6 is used as a marker for procentrioles associated with deuterosomes. Graphs show mean ± SD from n=7 cells across 3 different experiments. ****p<0,0001 using unpaired t-test. **(G)** Representative U-ExM images of brain MCCs during late A-stage. Boxes denote zoomed in regions on right, with 1-4 labels marking increasing distance from the parent centrioles. Cells were immunostained with antibodies to Acetylated-TUBULIN, PLK4, DEUP1. **(H)** Quantitation comparing percent of cells showing DEUP1 enrichment at daughter centriole over mother centriole (dc>mc), equal enrichment at both parent centrioles (dc = mc), or enrichment at mother centriole over daughter centriole (mc>dc) when cloud-stage and A-stage. A total of 54 cells were analyzed across 3 experiments. **(I)** Table summarizing protein localization as visualized by U-ExM along A-stage progression. “Cloud only” refers to the beginning of A-stage, when a cloud is visible but no procentriole-loaded deuterosomes can be identified. **(J)** Scheme representing A-stage progression in time and space.

As DEUP1 accumulates, some DEUP1+ foci within the cloud become bigger, more regular in shape and colocalize with PLK4 and SAS6, which marks the association of nascent procentrioles with deuterosomes (Fig. 3C-D, yellow arrow). These procentrioles are however deprived of β-tubulin and therefore of a microtubule wall (Fig. 3C). Such “naked” cartwheel assembly was also recently revealed as the earliest step of centriole assembly in cycling cells (Laporte et al., 2024). As the number of nascent procentrioles increases, they migrate away from the centrosome and β-tubulin is recruited asynchronously on procentrioles, with procentrioles further away from the parents acquiring β-tubulin, and procentrioles closer to the parents often lacking β-tubulin (Fig. 3E-F, Fig. 3 Supplementary 1G). Consistent with a progression of procentriole assembly during radial migration away from the centrosome, the procentriole walls increase in length from 80 nm to 150 nm and in width from 120 nm to 220 nm as a function of distance from the parent centrioles (Fig. 3 Supplementary 1H-I). The range and correlation of procentriole length and width increases are consistent with the recent description of S-stage procentriole growth (Laporte et al., 2024). Around the same time as β-tubulin, CENTRIN becomes also visible on procentrioles (Fig. 3 Supplementary 1J). This stage is probably the first stage we previously detected by CEN2-GFP live imaging and correlative CEN2-GFP light and EM microscopy (Al Jord 2014). The DEUP1 asymmetry to the daughter centriole previously described (Al Jord et al., 2014) becomes more visible than during earlier stage (Fig. 3 D-E, G-H). Centriolar microtubules are then acetylated on procentrioles migrating away from the parents (Fig. 3G, Fig. 3 Supplementary 1J). Procentrioles achieve their final width (Fig. 3 Supplementary 2 A-E) and will continue to elongate during G-stage – in the same range as described recently for G2 procentrioles – to finally reach the length of M-stage procentrioles during D-stage (Fig. 3 Supplementary 2 A-E, (Laporte et al., 2024)).

Altogether, these data reveal that (i) a pericentrosomal nest composed of CENTRIN, PCNT and DEUP1 is preassembled before the onset of centriole amplification, (ii) deuterosome and centriole biogenesis begin in this nest before what had been previously resolved using CEN2-GFP and EM, (iii) procentrioles recapitulate the S-stage step-wise dynamics, recently revealed by U-ExM time-series reconstruction in cycling cells, during their radial migration from the pericentrosomal nest, and (iv) their subsequent G- and D-stage elongation follows the G2- and M-like elongation process of the canonical cell cycle (Laporte et al., 2024). In MCC as in cycling cells, centriole elongation therefore follows the same step-wise dynamics, correlated with cell cycle stages (Fig. 3I-J).

### Dynein-dependent MT transport organizes the pericentrosomal nest and is required for deuterosome and centriole production

MEF-MCCs experiments suggest that the pericentrosomal cloud from which procentrioles emerge is organized by microtubules (Fig. 1 C-D). Consistently, in physiological brain MCC, dynamic tyrozinated microtubules are typically focalized to the centrosome before and during A-stage (Fig. 4A). This polarity is progressively lost during the following G- and D-stages to reappear at the basal body patch at the end of amplification, consistent with the known role of mature centriole appendages as MTOC in terminal MCC (Herawati et al., 2016; Kunimoto et al., 2012; Liu et al., 2020; Tateishi et al., 2017) (Fig. 4A).

**Figure 4:**
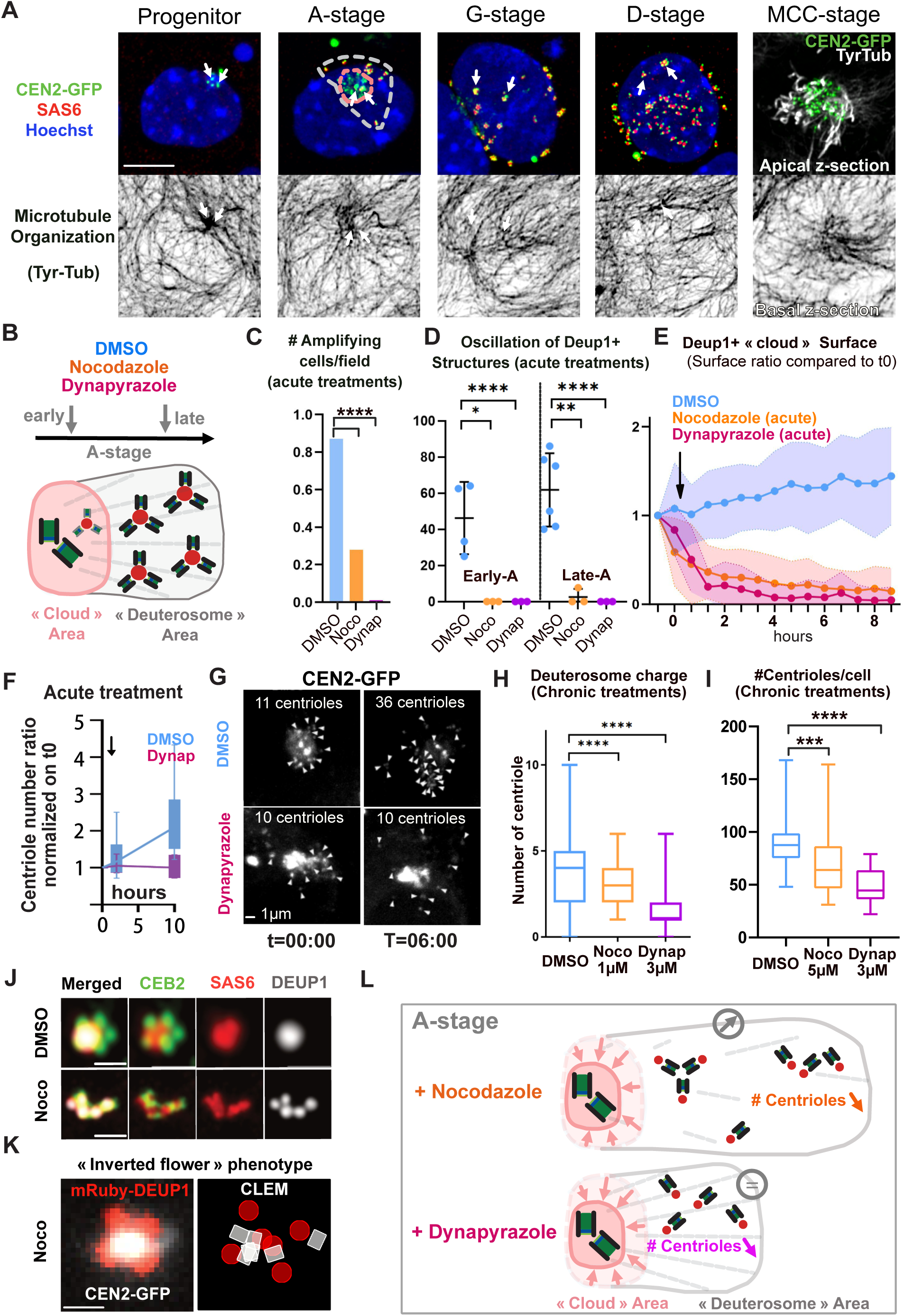
Dynein dependent MT transport organizes the pericentrosomal nest and is required for procentriole production. **(A)** Immunostaining profiles of procentrioles and microtubules during brain CEN2-GFP MCC differentiation. Procentrioles were counter-stained with anti-SAS6 antibodies (red), dynamic tyrosinated MTs were counter-stained with YL1-2 antibodies and the nucleus was counter-stained with Hoechst (blue). All micrographs are max projections of whole z-stacks except for the MCC-stage where max were done from apical or basal z-stacks selected from the same cell. Arrows show centrosomal centrioles, pink dotted line delineates the CEN2-GFP+ pericentrosomal cloud area, grey dotted line delineates the area occupied by deuterosomes. Scale bar, 5 µm. **(B)** Scheme depicting the experimental protocol. **(C)** Number of cells beginning centriole amplification (emergence of mRuby-DEUP1+/CEN2-GFP+ structures in the cell) in a microscope field during a 20h movie under acute DMSO (60 fields), Nocodazole (10 µM, 43 fields) or Dynapyrazole (3 µM, 30 fields) treatments. n>3 independent experiments. **(D)** Quantification of the percentage of early and late A-stage cells displaying at least one mRuby-DEUP1+ structure oscillation to and from the centrosome during 1-4min movies at dt5-15s, in DMSO, acute Nocodazole (10 µM) and acute Dynapyrazole (7.5 µM) treated cells. n>3 independent experiments were scored, n=81 DMSO cells, n=20 Nocodazole cells, n=56 Dynapyrazole cells analyzed. *p = 0.0108, **p=0.005, ****p<0.0001; Chi-square test with yates correction. See Videos 8-11. **(E)** Quantification of the area occupied by the mRuby-DEUP1+ cloud over time in control (DMSO), Nocodazole (10 µM) and Dynapyrazole (3 µM) treated cells. Black arrow indicates that DMSO and drugs were added right after the first acquisition. Three independent experiments were analyzed, n=12 DMSO cells, n=14 Noco 10 µM cells, n=14 Dynapyrazole 3 µM cells; ****p<0,0001; non-parametric Mann Whitney test; error bars represent mean ± SD. See Videos 12-14. **(F)** Centriole number ratio produced during A-stage under DMSO and Dynapyrazole (3 µM) treatment normalized on t0. DMSO and Dynapyrazole are added after the first time point. dt=40 min. **(G)** Representative images of CEN2-GFP movies used in (F). Arrowheads represent A-stage procentrioles. See also video 15. Scale bar, 1 µm. **(H)** Quantification of centriole number per deuterosome (using CEN2-GFP signal) in G-stage cells under DMSO, Nocodazole (1 µM) and Dynapyrazole (3 µM) chronic treatments (48h). Three independent experiments were scored, n_deut_=459 in DMSO, n_deut_= 215 in Nocodazole, n_deut_=383 in Dynapyrazole. Error bars represent min to max ± median. **** P<0.0001; non-parametric Mann Whitney test. **(I)** Quantification of SAS6+ centriole number in G- and D-stage (pooled) cells under DMSO, Nocodazole (1 µM) and Dynapyrazole (3 µM) chronic treatments (48h). Three independent experiments were scored; error bars represent min to max ± median. n=42 DMSO cells, n=34 Noco 1 µM; n= 29 Noco 5 µM cells analyzed. Error bars represent min to max ± median. ***<0.001, **** P<0.0001; non-parametric Mann Whitney test. **(J)** Super resolution immunostaining profile of deuterosomes (DEUP1) and procentrioles (SAS6) under chronic Nocodazole (10 µM, 24h). Deuterosomes arrange in grapes and are reduced to small subunits scarcely loaded with procentrioles. Scale bar, 500 nm. **(K)** Super resolution fluorescence of the mRuby-DEUP1;CEN2-GFP “inverted flower” profile frequently observed under chronic Nocodazole (10 µM, 24h) along with the underlying deuterosome and procentriole arrangement revealed by correlative light and electron microscopy (CLEM). See Fig. 4 Supplementary 5 for serial CLEM sections. **(L)** Scheme representing the effect of Nocodazole and Dynapyrazole on A-stage. To be compared with the scheme of Fig. 4B.

Because the pericentrosomal nest builds at the convergence of MT minus ends, we tested the role of MT and the MT retrograde molecular motors dyneins in the formation of the nest, the deuterosomes and the centrioles. To proceed, we live imaged MCC progenitors under acute nocodazole (10 µM) or dynapyrazole (3 µM) (Steinman et al., 2017) treatments, added after the first time point (see methods and Fig. 4B, Fig. 4 Supplementary 1A-B). We found a reduction or a complete abolition of the propensity of MCC progenitor cells to enter A-stage during a 24h period (Fig. 4C). When added on cells that had already entered A-stage, drug treatments block DEUP1+ structure oscillations within minutes in both early or late A-stage (Fig. 4D, video 8-11) and dissolves the mRuby-DEUP1+/CEN2-GFP+ cloud in few hours (Fig. 4E, Video 12-14). Under dynapyrazole, no new mRuby-DEUP1 foci appeared and no new centrioles were produced after drug addition (Fig. 4F-G, video 15). By contrast, in nocodazole, the existing deuterosomes moved away from the centrosome and some new mRuby-DEUP1+/CEN2-GFP+ foci appeared scattered in the cytoplasm (Fig. 4 Supplementary 2A, Video 12-14). Although they can increase in size, they are less regular and the associated CEN2-GFP+ signal is fainter (Video 12-13) suggesting the A-to-G transition is perturbed, probably because of a block in MT wall polymerization. This precluded the counting of final centriole number.

To confirm and further document the role of MT and dyneins, we also performed milder pharmacological treatments, but over longer timescale (“chronic” nococazole 48h, 5 µM or dynapyrazole 16h, 3 µM), and counted the number of amplifying cells, deuterosomes and centrioles. Consistent with live imaging, nocodazole and dynapyrazole treatments lead to a sharp decrease in the number of amplifying cells (Fig. 4 Supplementary 2B). In those undergoing centriole amplification, deuterosomes are more numerous but smaller and loaded with fewer procentrioles than in the control (Fig. 4H, Fig. 4 Supplementary 2C-D). Eventually, the final number of procentrioles is decreased (Fig. 4I). Altogether, these results show that MT and dyneins are required for the formation and maintenance of the pericentrosomal nest, the production of deuterosomes and the biogenesis of centrioles.

To further highlight how deuterosome formation is affected by MT alteration, we performed super-resolution imaging and CLEM on early A-stage MCC treated with nocodazole (10 µM, 24h). Super resolution imaging revealed the presence of grapes of small deuterosomes seemingly loaded with only one centriole, instead of the usual flower-like arrangement of procentrioles around a spherical deuterosomes (Fig. 4J). An “inverted flower” phenotype, where mRuby-DEUP1+ subunits are organized around a CEN2-GFP+ core was also frequently observed (Fig. 4K). CLEM on these structures confirmed the presence of single centriole-loaded deuterosomes (Fig. 4 Supplementary 3) and show that the “inverted flower” phenotype corresponds to an inversion of deuterosome/centriole organization where small deuterosomes, growing a single procentriole each, face each-others, regrouping procentrioles at the center (Fig. 4K, Fig. 4 Supplementary 4). CLEM furher showed that mRuby-Deup1+ structures can also correspond to groups of fibrogranular aggregates, probably resulting from pericentrosomal cloud scattering (Fig. 4 Supplementary 3). These aberrant organizations of procentrioles and small deuterosomes in a 1:1 stoechiometry under nocodazole suggest that concentration of DEUP1 at the centrosome is required for DEUP1 subunits to merge into large spherical deuterosomes. Non exclusively, other MT-dependent forces may be involved in the gathering of centriole/deuterosome subunits into larger deuterosomes.

Altogether, these data show that MT and dyneins are required to concentrate centriolar and deuterosome core components within a pericentrosomal nest. MT depolymerisation and dynein inhibition dissolve this nest and hinder or even block, the formation of deuterosomes and centrioles (Fig. 4B, L).

### Centriole maturation cycle superposes with centriole elongation cycle in the MCC cell cycle variant

We found that centriole elongation in MCC follows the same step-wise dynamics, correlated with cell cycle stages, as in the canonical cell cycle (Fig. 3J). A major difference however exists. After one canonical cell cycle the centriole cylinder is complete, has lost immaturity features such as cartwheel foundations, has been “modified” to gain MT nucleation capacity (Wang et al., 2011), but has not yet acquired full molecular maturity. This occurs during a second cell cycle and includes (i) the loss of daughter centriole proteins (e.g. centrobin, Cep97, Cep120) / acquisition of daughter centriole maturation proteins (e.g. C2CD3, ODF1, CEP350, CEP19, FOP) in S/G2 (Blanco-Ameijeiras et al., 2022), (ii) an enhanced capacity to nucleate MT, known as centrosome maturation at the G2/M transition (Joukov & De Nicolo, 2018), and (iii) the step-wise acquisition of distal (DA) and subdistal (SDA) appendages to dock at the membrane and anchor MT in early and late M stage respectively (Blanco-Ameijeiras et al., 2022). Called ‘daughter centriole’ at the end of the first cell cycle, the centriole is called “mother centriole” -or “basal body”- at the end of the second cell cycle (Fig. 5A) (Blanco-Ameijeiras et al., 2022; Nigg & Holland, 2018). By contrast, a single iteration of the MCC cell cycle variant allows mature basal bodies to form. We therefore sought to understand when and how centriole maturation occurs in this variant of the cell cycle.

**Figure 5:**
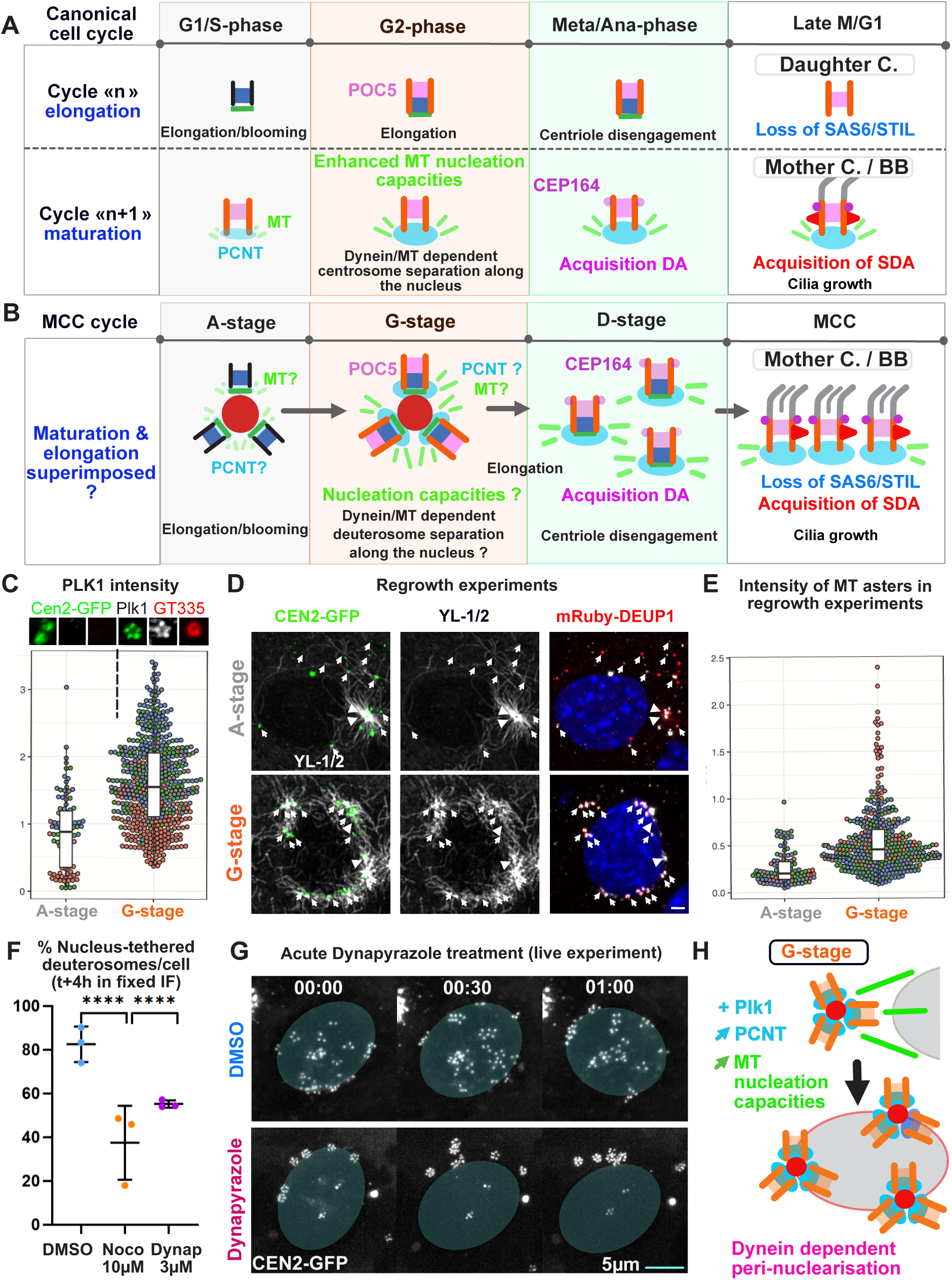
Centriole biogenesis is accelerated during a 2-in-1 MCC cell cycle variant. **(A-B)** Scheme comparing centriole biogenesis during centriole duplication in the canonical cell cycle and during centriole amplification in the MCC cell cycle variant. DA: distal appendages, SDA: sub-distal appendages, MT: microtubules. **(C)** Quantification of PLK1 intensity in A-stage and G-stage brain MCC samples using a linear regression model, with stages as the main effect and replicate as a fixed covariate, to account for batch variation. Statistical significance was assessed using Type II ANOVA. The lower and upper hinges correspond to the 25th and 75th percentiles. The upper (lower) whisker extends from the hinge to the largest (smallest) value no further than 1.5 * IQR from the hinge (where IQR is the inter-quartile range, or distance between the 25th and 75th percentiles); three independent experiments were quantified and colored differently; n= 32 cells; ****p<0,0001. Insets are from Figure 5 Supplementary 2A. **(D)** Immunostaining profile of YL1/2 in CEN2-GFP;mRuby-DEUP1 expressing brain MCCs in A- and G-stage during regrowth experiments (see methods). Arrowheads indicate centrosomal centrioles. Arrows indicate the position of A- or G-stage procentrioles. Scale bar, 1 µm. **(E)** Quantification of microtubule aster intensity in A-stage and G-stage brain MCC samples during regrowth experiments using a linear regression model, with stages as the main effect and replicate as a fixed covariate, to account for batch variation. Statistical significance was assessed using Type II ANOVA. The lower and upper hinges correspond to the 25th and 75th percentiles. The upper (lower) whisker extends from the hinge to the largest (smallest) value no further than 1.5 * IQR from the hinge (where IQR is the inter-quartile range, or distance between the 25th and 75th percentiles); three independent experiments were quantified and colored differently; n= 56 cells; ****p<0,0001. **(F)** Quantification of the percentage of tethered deuterosomes per G-stage cell in DMSO , Nocodazole (10 µM, 4h) and Dynapyrazole (3 µM, 4h). Deuterosomes above or below the nucleus are not quantified. Four independent experiments quantified; n=15 cells in DMSO, n=26 cells in Nocodazole, n=17 cells in Dynapyrazole. Error bars represent mean ± SD; ****p<0,0001; Chi-2 test with Yates’ correction. **(G)** Still images from a movie of G-stage CEN2-GFP expressing cells treated with DMSO or Dynapyrazole 3 µM at t=0. While “centriole-loaded deuterosomes” keep a perinuclear distribution in DMSO treated cells, in Dynapyrazole treated cells, the “centriole-loaded deuterosomes” start untether from the nucleus at t+30 min. Shaded grey ovals outline the nucleus identified with contrasting CEN2-GFP. scale bar, 5 µM. See Video NEW17. **(H)** Scheme synthesizing the main findings of main and supplementary Fig. 5 on G-stage procentrioles maturation and nuclear tethering.

Since (i) centriole elongation in MCC follows the dynamics of a canonical cell cycle (Fig. 3J), (ii) elongation and maturation cycles are driven by the same molecular cascade, (iii) which is also common to the MCC cell cycle variant (Al Jord et al., 2017), a parsimonious way of accelerating centriole biogenesis in the MCC variant would be to superpose the maturation “n+1” cycle, to the elongation “n” cycle. Consistent with this scenario, we found in our previous studies that the acquisition of centriole appendages is step-wise, and occurring in the second half of the MCC variant, like in the canonical cycle: distal appendages are acquired at the G-to-D transition (Al Jord 2014) and subdistal appendages (basal feet) during early MCC (Fig. 5B, (Guirao et al., 2010)). Also, our single cell RNA sequencing data set comparing MCC and canonical cell cycle variants in brain progenitors (Serizay et al., 2025), shows that the expression of proteins involved in the “n” elongation cycle (Fig. 5 Supplementary 1A), superposes with the expression of proteins involved in the “n+1” maturation cycles (Fig. 5 Supplementary 1B) during the MCC variant. Additionnaly, at the end of MCC differentiation, procentrioles transiently display both “n” (SAS6+ foundations) and “n+1” (CEP164 appendages) characteristics (Al Jord et al., 2014), which is never observed in cycling cells.

To further test whether a true superposition exists, we tested whether and when the remaining maturity feature, i.e. the enhanced MT nucleation capacities, was happening during the MCC variant. In the canonical cell cycle, this is called centrosome maturation, occurs at the G2/M transition and is dependent on PCNT recruitment and the activation of an AurA-PLK1 kinase cascade(Joukov & De Nicolo, 2018)(Blanco-Ameijeiras et al., 2022). Staining for PLK1 and PCNT in MCC progenitors, we observed that PLK1 becomes visible at procentrioles during G-stage and that G-stage procentrioles recruit significantly more PCNT than A-stage procentrioles (Fig. 5C, Fig. 5 Supplementary 2A-C). This is true even in the absence of deuterosomes (DEUP1 knock out cells, Fig. 5 Supplementary 2D). To test whether procentrioles displayed enhanced MT nucleation capacities at the A-to-G transition, we performed MT regrowth experiments (see methods) in both MEF-induced and brain MCCs. We found that GT335+ G-stage procentrioles display a microtubule regrowth efficiency significantly greater than A-stage procentrioles which show a weak ability to nucleate MT (Fig. 5D-E, Fig. 5 Supplementary 2E-F). Consistent with a process comparable to the centrosome maturation occurring at the G2/M transition, scRNAseq analysis reveals that proteins involved in centrosome maturation (PLK1, AURKA, CEP192, PCNT) are expressed during the MCC variant (Fig. 5 Supplementary 1C),

In the canonical cell cycle, centrosome maturation is concomitant with the dynein dependent attraction of centrosomes to the nucleus and their consecutive migration on the nuclear membrane for bipolar spindle formation at mitosis onset (Fig. 5A). Interestingly, in MCC, the A-to-G transition is marked by a collective migration of centriole-loaded deuterosomes from the centrosome to the nucleus where can arrange as equally distant structures forming a belt around the nucleus (e.g in Fig. 6 Supplementary 1A, (Al Jord et al., 2017, 2019)). Live imaging mRuby-DEUP1/CEN2-GFP cells at this stage confirms this switch and further reveals that procentriole-loaded deuterosomes can migrate for several hours along the nuclear membrane, suggesting a strong and dynamic attachment (Fig. 5 Supplementary 2G-H, video 16). Nucleus migration, tethering and physical distancing suggest that the same mechanisms driving newly formed centrosomes to the opposite poles of the nucleus during early prophase, are also used by G-stage centrioles loaded on deuterosomes, to organize around the nucleus. Consistently, scRNAseq analysis reveals that most of the proteins involved in centrosome migration to the nucleus (CENPF and dynein subunits DYNLRB2, DYNLL1), centrosome dysjunction (TPX2, AURKA, NEK2) are expressed during the MCC variant and with a dynamics globally comparable to the canonical cell cycle (Fig. 5 Supplementary 1C). To further test this hypothesis, we treated MCC with nocodazole (10 µM) for 4h and found that this prevented the migration of G-stage procentrioles to the nuclear membrane, which was reversed in washout experiments (Fig. 5F, Fig. 5 Supplementary 2I-J). In addition, live treating cells in which centrioles have already migrated to the nucleus with nocodazole (10 µM) induce their detachment within minutes (Fig. 5 Supplementary 2K). A block in centriole tethering during G-stage was similarly observed in fixed and live conditions under dynapyrazole treatments (Fig. 5F-G) showing that centriole migration to the nuclear membrane is driven by a dynein and MT dependent mechanism, like during centriole duplication in the canonical cell cycle (Fig. 5H).

**Figure 6:**
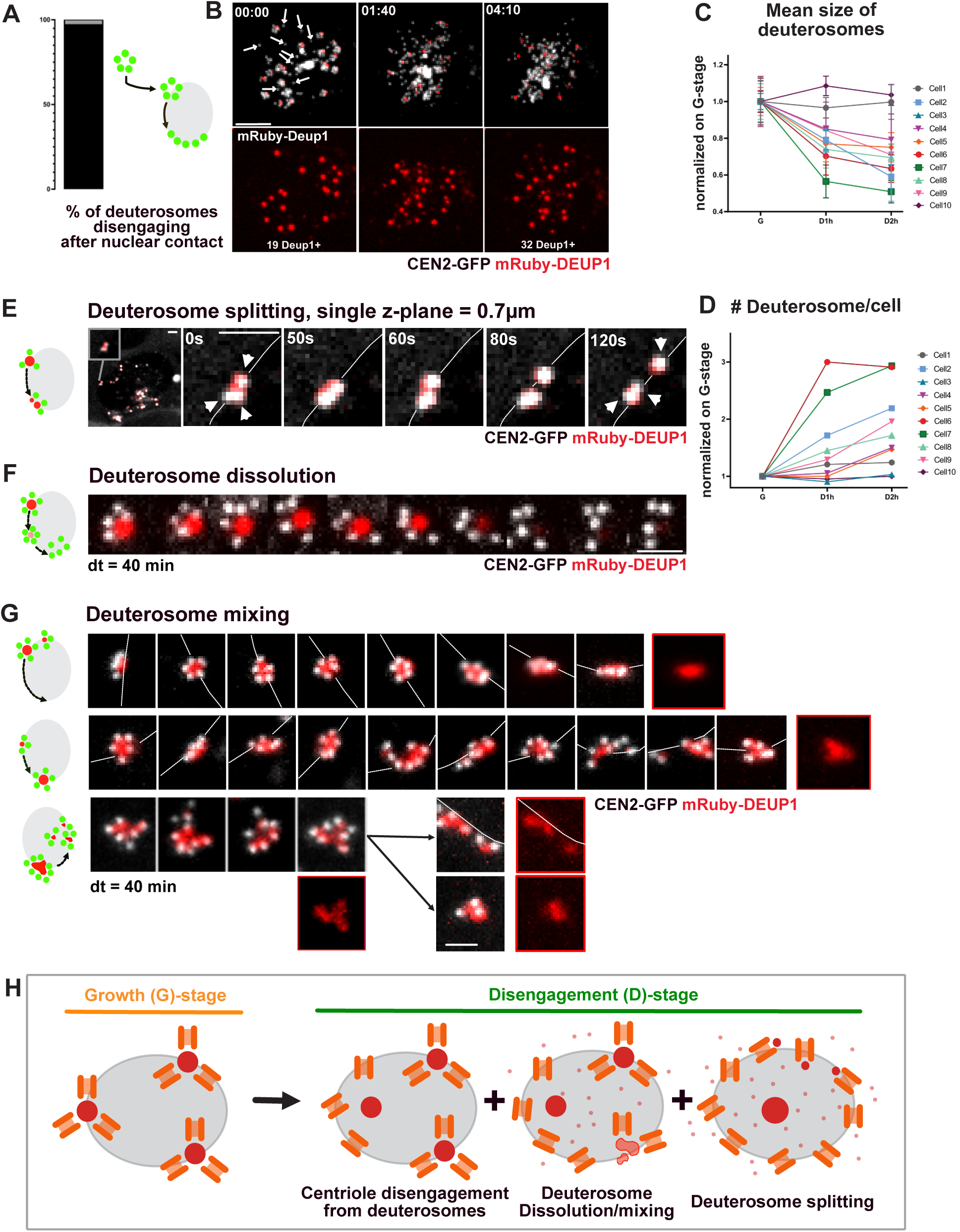
Disengagement of centrioles in MCC involves deuterosome splitting and dissolution. **(A)** Quantification of the proportion of procentrioles disengaging with a nuclear contact. Three independent experiments were quantified; n=98 deuterosomes, n=8 cells. See also Video 17. **(B)** CEN2-GFP; mRuby-DEUP1 dynamics in D-stage. Upper panels: in early D-stage (00:00), some procentrioles are already individualized, separated from DEUP1+ deuterosomes as shown by the white arrows. During D-stage, procentrioles disengaged from deuterosomes can keep a subunit of DEUP1 in their proximal portion, splitting them in smaller entities bearing fewer procentrioles not disengaged yet, as well as DEUP1 signal dissolution. Lower panels: grayscale showing the initial pool of 19 deuterosomes (00 :00) splitting to 32 DEUP1+ foci (4 :10). dt=50min; scale bar, 5 µM. See also Video 19. **(C-D)** Quantification of the size and number of deuterosomes per cell over time in D-stage relative to G-stage. n=10 cells analyzed were plotted individually to reveal the tendency to decrease in size, while increasing the number of DEUP1 foci. Each measurement in D-stage was normalized on G-stage. Three independent experiments were scored; error bars represent mean ± SD. **(E)** CEN2-GFP;mRuby-DEUP1 time lapse images of a single z-plane (0.7µm) of a D-stage cell showing a deuterosome which seemingly split into smaller deuterosomes migrating away from each others along the nuclear envelope. White dotted line outlines the nuclear membrane identified with contrasting CEN2-GFP signal. Left picture is the zoom-out picture of the D-stage cell, showing the position of the cropped still images. dt=5s, scale bar, 1 µm. See Video 20 to see the entire movie. **(F)** Time lapse images of a D-stage cell showing the deuterosome mRuby-DEUP1+ signal dissolution correlated with CEN2-GFP rosette dismantlement. dt=40min, scale bar, 1 µm. See also Video 21. **(G)** CEN2-GFP;mRuby-DEUP1 time lapse images of a D-stage cell showing migration of a centriole-loaded deuterosome along the nuclear membrane (line 1), change of deuterosome shape and mixing with another deuterosome (line 2), followed by consecutive unmixing (line 3). dt=40min, scale bar, 1 µm. See Video 22 to see the entire movie. **(H)** Scheme synthesizing the main findings of main and supplementary Fig. 6 on D-stage deuterosome dynamics and centriole disengagement.

Collectively, these data show that in MCC, the elongation of procentrioles respects the step-wise dynamics observed during the canonical “n” cycle (Fig. 3J, 5A) and is superimposed by the acquisition of maturity features, which respects a canonical step-wise dynamics comparable to a canonical “n+1” cycle (Fig. 5B). By superimposing the two processes, the MCC variant skips the daughter centriole step and drives the production of mature basal bodies in a single cycle iteration (Fig. 5A-B). In addition, this “2-in-1” cycle confers procentrioles with high MT nucleation properties by G-stage, allowing their efficient self-organization around the nuclear membrane. Given the similarities in both centriole dynamics and expression of core molecular regulators, this perinuclearisation probably occurs through the same mechanisms as centrosome bipolarisation at mitotic entry (Fig. 5H) (Agircan et al., 2014).

### Disengagement of centrioles in MCC involves deuterosome splitting and/or dissolution at the nuclear membrane

Once at the nuclear membrane, G-stage procentrioles disengage from deuterosomes or centrosomal centrioles, which constitute the so-called disengagement D-stage (Al Jord et al., 2017). Single deuterosome tracing through CEN2-GFP monitoring show that disengagement always occurs in contact with the nuclear membrane (Fig. 6A, Fig. 6 Supplementary 1A, video 17). Consistently, super-resolution imaging on CEN2-GFP+ G- and -D-stage cells immunostained with anti-nuclear pore complex proteins reveals procentrioles intermingled within nuclear pores (Fig. 6 Supplementary 1B, video 18). The global dynamics of disengagement has previously been characterized as the dismantling of empty and plain CEN2-GFP+ flower-like structures, representing CEN2-GFP+ procentrioles organized as flowers around CEN2-GFP negative deuterosomes or CEN2-GFP+ centrosomal centrioles respectively (“deuterosome” pathway and “centriolar” pathways; Fig. 6 Supplementary 1A, (Al Jord et al., 2014)). To further highlight deuterosome behavior during disengagement, we monitored in live single cells expressing mRuby-DEUP1 in addition to CEN2-GFP.

At the global cell scale, the beginning of disengagement is marked by the apparition of single DEUP1 negative procentrioles suggesting that centrioles can detach from deuterosomes, like they disengage from parental centrioles in cycling cells (Fig. 6B, 00:00, white arrows). Then, quantifications reveal that the size of deuterosomes decreases while their number increases over time, in single cells, indicating that deuterosomes split during disengagement (Fig. 6B; 01:40; Fig. 6C-D, 10 cells; Video 19). In parallel, a DEUP1 diffuse signal, absent during G-stage, appears in the cytoplasm suggesting that splitting is accompanied by DEUP1 structure dissolution (Video 19). Sometimes, at the end of disengagement, mRuby-DEUP1+ structures aggregate into larger structures until ciliation, before dissolving (Fig. 6 Supplementary 1C).

We then increased spatial resolution and followed single deuterosomes. This allowed to observe deuterosomes that seemingly split around the nuclear membrane, giving rise to deuterosomes with single centrioles (Fig. 6E, Video 20). Also, we identified that dissolution of mRuby-DEUP1+ signal from the center of CEN2-GFP+ flowers was concomitant with centriole distancing (Fig. 6F, Fig. 6 Supplementary 1D, Video 21). Finally we found that deuterosomes can loose their spherical shapes and/or intermingle with others before the mRuby-DEUP1 signal split and finally disappears (Fig. 6G, video 22). All these events are observed along the nuclear membrane.

Altogether this suggests that disengagement is mediated by a change in the physical properties of DEUP1 aggregates allowing the splitting and/or dissolution of deuterosomes (Fig. 6H). Whether this results from post-translational modifications of DEUP1 by APC/C, PLK1 and/or separase, which are known to be required for centriole disengagement in both cycling cells and MCC, requires further examination (Al Jord et al., 2017; Prosser et al., 2012; Revinski et al., 2018; Tsou et al., 2009; Tsou & Stearns, 2006). In favor of a common molecular control, centriole disengagement from centrosomal centrioles and deuterosomes occurs concomitantly in differentiating MCC (Fig. 6 Supplementary 1A), and the dynamics of disengagement (duration and perinuclear location) of doublets or triplets of centrioles when deuterosomes are absent (DEUP1 knock-out cells), seems comparable to controls (Mercey, Levine, et al., 2019).

### Disengagement of centrioles in MCC is assisted by MT and dyneins

Since centrioles disengage along the nuclear membrane which they reach through a dynein dependent process, we then sought to test if mechanical forces are also involved in centriole disengagement. In fact, if perinuclearisation mechanisms are the same in MCC progenitors and cycling cells, it means that dyneins bound to nuclar pores are running on MT nucleated by individual G-stage procentrioles. We reasoned that this would lead to procentrioles being pulled apart from each other’s, which could contribute mechanically to deuterosome splitting and centriole disengagement. In favor of such a scenario, D-stage centrioles are on the same z-section (250 nm) as nuclear pores (Fig. 6 Supplementary 1B, Video 18), they continue to migrate along the nuclear membrane after deuterosome splitting (Fig. 7A, Video 23), and nocodazole (10 µM) induces their immediate detachment from the nucleus (Fig. 7 Supplementary 1A). Also, MT regrowth experiments show that single D-stage centrioles are nucleating MT (Fig. 7B) and super-resolution confocal imaging show that disengaging centrioles are co-localizing with MT (Fig. 7C, video 24). Finally, we identified a transient stage during which disengaging procentrioles redistribute in the 3 dimensions along the nuclear membrane, which is coherent with the existence of isotropically distributed traction forces on the nuclear surface (Fig. 6 Supplementary 1B, 4:30, video 18).

**Figure 7:**
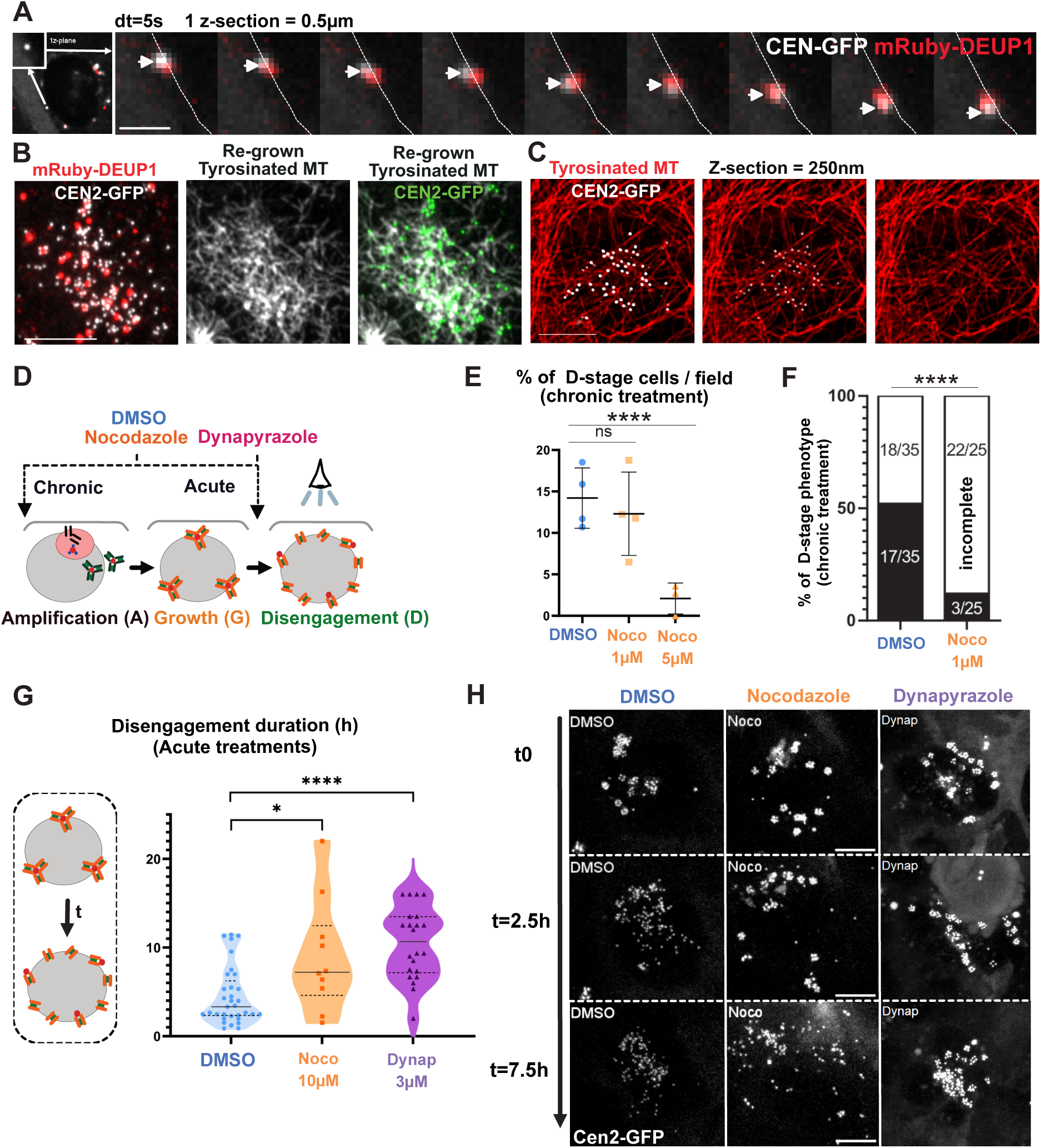
Disengagement of centrioles is assisted by MT and dyneins. **(A)** Disengaged procentriole associated to a deuterosomal subunit migrating on the nuclear envelope. White arrow head indicates the centriole, the white dot line outlines the shadow of the nuclear membrane seen in CEN2-GFP. dt=5s; Single z-section of 0.5 µm, scale bar, 2 µm. See Video 23 to see zoom out and whole movie. **(B)** Representative picture of tyrosinated microtubules (YL1/2) and DEUP1 immuno-reactivity in brain CEN2-GFP MCC during D-stage after MT depolymerization through nocodazole treatment (10 µM, 24h) and subsequent nocodazole whashout (4 h). Scale bar, 5 µm. **(C)** Tyrosinated microtubules and deuterosomes immuno-reactivity profiles in brain CEN2-GFP MCC in D-stage. These pictures show one z-section (0.25 µm) in the basal part of the cell. Tyrosinated MTs were stained with (YL 1-2). CEN2-GFP signal intensity is decreased from left to right to show that centrioles are located next to MT. Scale bar, 5 µm. See Video 24 to see all z-sections and DEUP1 staining. **(D)** Schematic representation of brain MCC differentiation depicting the timing of drug treatments. The eye shows the amplification stage monitored. **(E)** Quantification of the proportion of D-stage cells per fields among amplifying cells under chronic DMSO and Nocodazole treatments (48h, 1 µM and 5 µM). At least three independent experiments were quantified, n=2626 DMSO cells, n=1483 Noco 1 µM cells, n=909 Noco 5 µM cells; error bars represent mean ± SD. ns, not significant; ****p<0,0001; non-parametric Mann Whitney test. **(F)** Quantification of D-stage quality phenotype under chronic DMSO and Nocodazole (48h, 1 µM). Total disengagement in black, incomplete disengagement in white. At least three independent experiments were quantified, n=35 DMSO cells analyzed, n=25 Noco 1 µM cells analyzed. ****p<0,0001; Chi-2 test with Yates’ correction. **(G)** Quantification of disengagement duration in CEN2-GFP brain MCCs under DMSO and acute Nocodazole (10 µM) or Dynapyrazole (3 µM) treatments. >4 independent experiments were quantified, n=32 DMSO cells, n=10 Nocodazole cells and n=22 Dynapyrazole cells. *p<0,05; ****p < 0/0001 non-parametric Mann Whitney test. **(H)** Representative movie of CEN2-GFP dynamics in D-stage of brain MCCs under DMSO and acute Nocodazole (10 µM) or Dynapyrazole (3 µM) treatments. (t-1) represents the timepoint before drug treatments. scale bar, 5 µm. See Video 25 for whole movies.

To functionally test whether MT attachment of the centrioles to the nuclear membrane assists mechanically their disengagement, we chronically treated brain MCCs with increasing doses of nocodazole (1-5 µM, 48h) and analyzed whether procentrioles were able to individualize. We observed that at low doses, the proportion of cells in “D-stage” remained unchanged, but significantly more “D”-stage cells showed incomplete disengagement compared to controls (Fig. 7D-F). Interestingly, within the cells showing incomplete disengagement, we often saw SAS6 negative procentrioles still engaged to deuterosomes, a phenotype rarely observed in controls (Fig. 7 Supplementary 1B-C, white arrows). Indeed, SAS6 normally disappears from procentrioles when centrioles are docked, just before ciliation (Al Jord et al., 2014). This suggests that, despite an active APC/C (responsible for SAS6 degradation in cycling cells (Arquint & Nigg, 2016), and involved in centriole disengagement in MCC (Al Jord et al., 2017)), centrioles failed to disengage from deuterosomes. Then, with higher dose of nocodazole (5µm), “D”-stage cells are rarely observed (Fig. 7E). These observations suggest that microtubules are required for efficient centriole disengagement. To confirm and to further assess the role of dyneins, we performed acute nocodazole (10 µM) and dynapyrazole (3 µM) treatments on G-stage cells, and live-monitored the dynamics of disengagement. We observed, in both cases, a detachment of procentriole-loaded deuterosomes from the nuclear membrane and a long, unsynchronized and sometimes incomplete disengagement (Fig. 7 G-H, video 25).

Altogether these data suggest that the acquired ability of the newly formed centrioles to nucleate microtubules at the A-to-G transition, allows their dynein-dependent nuclear attachment, which assists mechanically their disengagement during the following D-stage.

### Disengaged centrioles finally converge apically with the former centrosome

After disengaging, centrioles detach from the nuclear membrane (Fig. 6A; 10:00) and migrate from all over the nucleus to the apical cell surface (Fig. 6 Supplementary 1A; 10:00-14:00) to dock and nucleate motile cilia (Boisvieux-Ulrich et al., 1987, 1990; Boisvieux-Ulrich et al., 1989). This is followed by an acto-myosin dependent planar basal body constriction (Hirota et al., 2010). Dynamics of this collective migration process has never been analyzed and is tedious since MCCs produce several dozens of centrioles that are therefore difficult to track. To begin, we first analyzed the global 3D displacement of all centrioles in individual cells, by segmenting the CEN2-GFP signal throughout their apical migration over time-lapse imaging. As a cellular reference, we took one of the centrosomal centrioles, trackable using their higher CEN2-GFP fluorescence, in cells where the 2 centrosomal centrioles stay close together. This global quantification showed that, at the end of disengagement, centrioles are evenly distributed within a mean radius of 8 µM with respect to the tracked centrosomal centriole (in the range of the nucleus radius). Then, by the end of apical migration and basal body constriction, centrioles are predominantly (65%) located in a radius of 3 µM around this reference point (Fig. 8A). This significant histogram shift (Chi² test, p<0.0001) suggests a collective movement of newly formed centrioles that bridges them together and with the centrosomal centrioles. Consistently, centrosomal centrioles finally dock within the newly formed basal body patch and also nucleate cilia (Liu et al., 2020).

**Figure 8:**
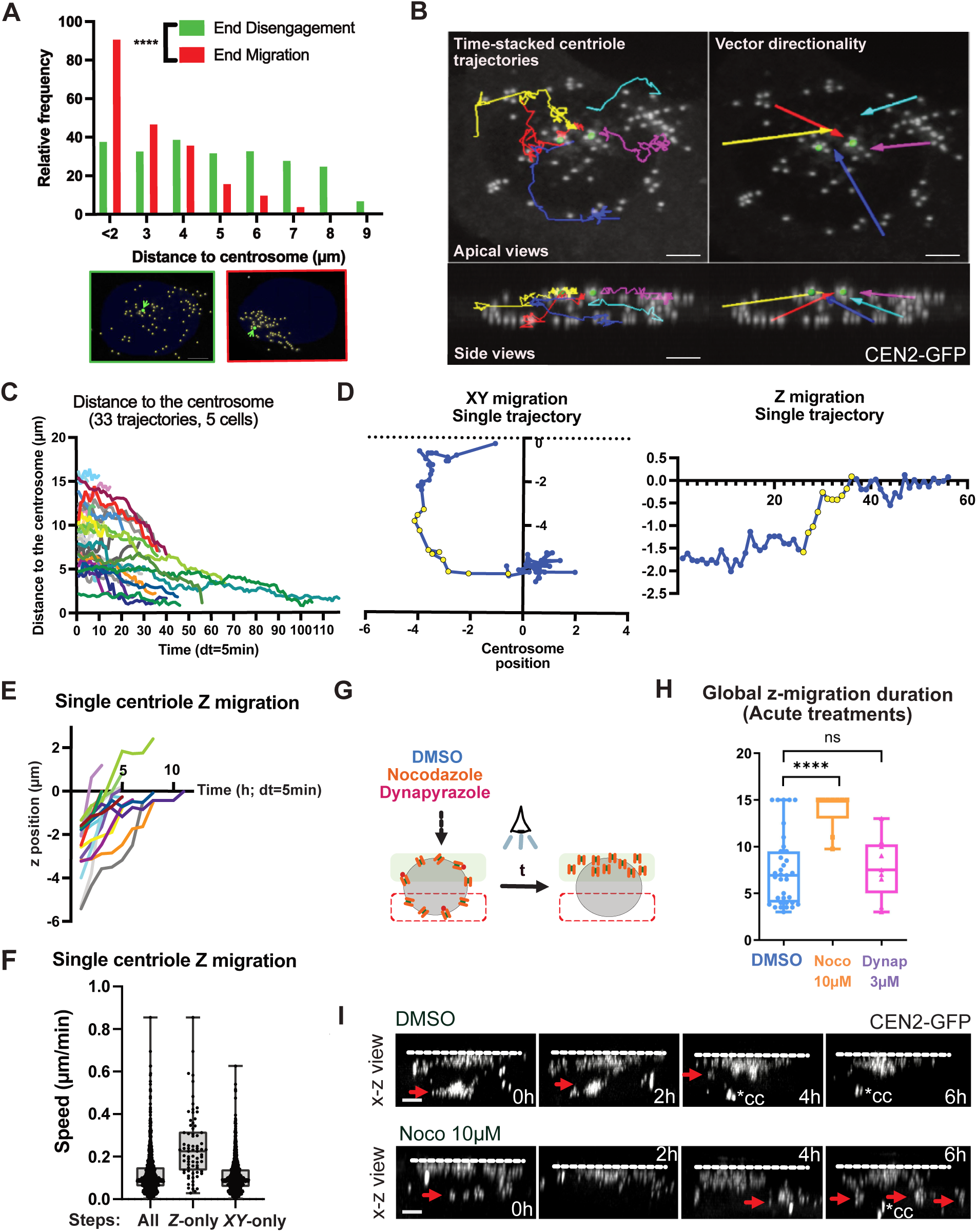
Disengaged centrioles finally converge with the former centrosome. **(A)** Quantification of the relative frequency of the centriole position regarding the centrosomal centriole (CC) at the end of centriole individualization (green) and the end of apical migration (red). Bottom: representative apical projections of segmented CEN2-GFP+ centrioles at the end of centriole individualization (green frame) and the end of apical migration (red frame). Green arrow shows the centrosome which served as the reference point; the nucleus is in dark blue. n=3 cells; scale bar, 5 µM. **(B)** Manual single centriole 3D tracking reveals convergence of centrioles towards the centrosome region. Top left panel: apical view of manually tracked migrating centrioles trajectories. Top right panel: apical view of the directionality vectors of the migrating centrioles. Bottom panel: corresponding profile views of the top panels. dt=5min; scale bar, 5 µM. See Video 26. **(C)** Evolution of individual centriole distance to centrosome over time. dt=5min. Three independent experiments were analyzed, n=33 centriole trajectories in n=5 cells. **(D)** Example of the single blue centriole trajectory from (B) towards the centrosome in the XY plane and in the Z plane. Yellow dots represent the steps of apical migration. The origin represents the position of the centrosome. See Video 26. **(E)** Evolution of the Z coordinates during the apical migration of 16 centrioles from 5 cells. Three independent experiments were analyzed. The origin represents the Z position of the centrosome. dt=5min. **(F)** Speed of migration plotted from all 953 steps of 33 migrating centrioles versus the 74 extracted steps of apical migration of the 16 centrioles basally located in the cells. Box and whiskers plots (2,5-97,5 percentile). See distribution in Fig. 8 Supplementary 1C. **(G)** Schematic representation of brain MCC differentiation depicting the timing of MT depolymerisation treatment. The eye shows the amplification stage monitored. **(H)** Quantification of apical migration duration in DMSO and acute nocodazole (10 µM) or dynapyrazole (3 µM) in brain MCCs. Eight independent experiments were scored for n=32 DMSO cells, 3 experiments for n=10 nocodazole cells and 5 experiments for n=8 dyanpyrazole cells. Error bars represent min to max ± median. ****p<0,0001; non-parametric Mann Whitney test. **(I)** Profile view of centriole apical migration after D-stage in CEN2-GFP+ DIV2 brain MCCs treated with acute DMSO and Nocodazole 10 µM. At t-1, cells display late G-stage/early D-stage procentrioles and are not treated with any drug. DMSO and Nocodazole 10 µM were added right after the first timepoint acquisition. In DMSO, 2 centriole patches are above and below (red arrow) the nucleus (t+0h). The patch below the nucleus progressively joins the apical one (t+2h) to become one apical patch (t+4h) and stabilizes (t+6h). In 10 µM Nocodazole, procentrioles start to scatter after 1h, and the group of centrioles below the nucleus travel up and down (t+2h, t+4h and t+6h) and does not succeed to become one apical group of centrioles (t+6h). Red arrowheads point to group of centrioles that we consider as basally localized. *CC corresponds to the centrosome of a neighboring cell. dt=5min; scale bar, 5 µm. See also Video 27.

To then analyze the displacement dynamics of individual centrioles, we performed live imaging with high temporal resolution (dt=5min) over the entire migration process. This allowed us to manually track a subset of migrating centrioles and catch their individual trajectories before they reach the group (Fig. 8B, Video 26). Single centrioles move at 0,12+/-0,09 µm.min^−1^. Consistent with the global 3D gathering of the whole centriole population, they are seen converging apically toward the centrosomal centriole reference point as reflected by the time-correlated decrease of their distance and by the mean vector directionalities of individual trajectories (Fig. 8B-C, Video 26). Although single centrioles display a global directional migration, analysis of the shape of the trajectories, speed of displacement and directionality with regard to the centrosome reference point, show that their migration is complex and marked by back-and-forth movements oscillating between diffusive-like and more processive steps (Fig. 8D, Fig. 8 Supplementary 1A). As steps toward the centrosome are more numerous and more rapid than steps away (Fig. 8 Supplementary 1B), centrioles finally slowly converge at 0,01+/-0,03 µm.min^−1^. Interestingly, within this complex behavior, the vertical z-motion, when centrioles below the nucleus migrate up around the nuclear membrane (Fig. 8B; light blue, dark blue and red trajectories), is clearly processive (Fig. 8D, yellow dots; Fig. 8E) and more rapid (0,25+/-0,16 µm.min^−1^, Fig. 8F, Fig. 8 Supplementary 1C), than the steps in the XY plane (0,11+/-0,08 µm.min^−1^).

To define the involvement of microtubules and dyneins in this collective migration process we compared the global dynamics of migration in control and cells treated with nocodazole (10 µM) or dynapyrazole (3 µM) from the end of D-stage. Under these treatments, the centrosomal centrioles moved too much to be traced. In the absence of a reference point, we restricted our analyzis to the quantification of the baso-apical migration. This z-migration was defined as being the duration between the take-off of the first centriole of the group below the nucleus, until the last one is reaching the apical side of the cell (Fig. 8G). While in the control, global z-migration duration lasted 7,5 +/- 3,9h, under nocodazole, complete migration was generally not reached during the course of the 15h live imaging experiments (migration duration = 14 +/- 2h) and was marked by up and down migration of groups of centrioles (Fig. 8H-I, video 27). Interestingly, under dynapyrazole treatment, even if D-stage centrioles detach from the nuclear membrane, they subsequently succeed in migrating up, with a dynamics and a duration comparable to the controls (Fig. 8H, migration duration = 7,8 +/- 3,2h). This suggests that, like for apical centrosome migration during primary ciliogenesis, the dynamics is MT dependent, but dynein-independent (Pitaval et al., 2017). Interestingly though, when waiting longer by treating the cells chronically with nocodazole for 48h (1-5 µm), centrioles seem to finally succeed in migrating up (Fig. 8 Supplementary 1D), even if they fail to gather accurately in the XY plane (Fig. 8 Supplementary 1E-F). This may explain why the first report on quail oviduct MCCs failed to detect any effect of MT depolymerization on final centriole apical positionning (Boisvieux-Ulrich et al., 1989).

Altogether, these data show that centrioles migrate collectively, following a complex and slow motion, from all over the nuclear membrane, to gather with the former centrosome. The migration is characterized by fluctuations between forward and backward directionalities and between processive and diffusive-like movements, but switches to a more processive and rapid motion when directed toward the apical pole. This migration is altered under microtubule depolymerization but seems independent from dynein motors. Wether the apical migration is dependent on actin and Cep164-dependent MT stabilization pushing the centrioles apically, like proposed for primary ciliogenesis, requires further examination (Pitaval et al., 2017).

## Discussion

Mature centrioles can be produced in less than an hour in amoebae differentiating into flagellates (Fritz-Laylin et al., 2016). However, in the large majority of vertebrate cells, centriole biogenesis is curbed within a double iteration of the cell cycle, allowing to meet the need of a single primary cilium at the end of cell division. Here, we show that the recently described MCC cell cycle variant is an accelerated version of the cell cycle which superposes two centriole biogenesis cycles (elongation and maturation), to obtain multiple mature centrioles within a single cycle iteration. The precocious maturation of amplified procentrioles is even determinant for their spatial self-organization, disengagement and apical migration.

### A pericentrosomal nest

Using live imaging on brain MCC, we highlight that concentration of core deuterosome and centriole proteins around the centrosome precedes the biogenesis onset of deuterosomes and procentrioles. We named this transitory compartment a “nest” since deuterosomes and procentrioles emerge specifically in this region and grow while moving away from it. We highlight that this nest is composed, at least, of DEUP1, PCM1, PCNT and CENTRIN2 and is dependent on MT and dyneins. Its dissolution through MT depolymerization or dynein inhibition is correlated with an alteration -or even a block- in the formation of deuterosomes and procentrioles, suggesting that concentration of core elements of deuterosome and centriole organelles permit their efficient massive production.

This concentration of proteins can result from the retrograde transport of the proteins themselves, as suggested by the movements of DEUP1 or CEN2-GFP foci. Non exclusively, it could result from retrogradely transported mRNA and local translation at centriolar satellites. In fact, the core component of centriolar satellites PCM1 (i) is highly enriched in the pericentrosomal nest, (ii) diffuse when MT are depolymerized, and (iii) has been recently shown to be sites of local translation of centrosomal and ciliary proteins in vertebrate cells (Pachinger et al., 2025). Local translation of centriolar and ciliary proteins, notably at the centrosome, has been shown to be involved in both centriole (over)duplication and primary ciliation (Fang & Lerit, 2022; Martinez et al., 2025; Pachinger et al., 2025; Villa et al., 2025). In brain and airway MCC, PCM1 depletion alters deuterosome formation and centriole production (Hall et al., 2023; Zhao et al., 2021). Such concentration of centriolar proteins by a retrograde MT transport would explain why, (i) in DEUP1 KO MCC where deuterosomes are absent, centrioles are emerging on, and around, the centrosomal centrioles (Mercey et al., 2019), and (ii) when both centrosome and deuterosomes are absent, procentrioles emerge in a cloud of PCM, at the convergence of the microtubule network (Mercey et al., 2019) . Our observations are also consistent with theoretical modeling, simulations and *ex cellulo* experiments suggesting that the place where a critical threshold level of PLK4 activation is surpassed determines procentrioles’ apparition spot (Nabais et al., 2021; Yamamoto & Kitagawa, 2019).

Whether the existence of such a pericentrosomal nest exists in respiratory and reproductive cells is unclear. Live imaging is difficult to perform because cells are in a stratified epithelium. In addition, the number of centrioles produced is much greater than in brain cells, and their production seems to be rapid. Consequently, the early dynamics of amplification is difficult to describe. However some studies in fixed cells at early stages observed a pericentrosomal onset of procentrioles and deuterosomes emergence in chick and mouse airway MCCs (Al Jord et al., 2014; Kalnins et al., 1972; Lu et al., 2025; Mori et al., 2017). Also, our observations in non-tissue specific micro-patterned MEF-MCCs suggest that, at least the onset of amplification, occurs in the pericentrosomal region, in a MT-dependent manner. One can hypothesize that, in airways MCC producing more centrioles, the concentration of centriolar proteins may reach the threshold for centriole biogenesis in the pericentrosomal region first, and then rapidly in the rest of the cytoplasm. Also, pericentrosomal localization of pericentriolar satellites is not a conserved feature (Pachinger et al., 2025) and may differ in airway MCC.

### Dynamics in the pericentrosomal nest and earliest steps of deuterosome formation

The pericentrosomal nest hosts the formation of early deuterosomal structures whose size and shape are diverse. Correlative light and electron microscopy reveals that early deuterosomes can appear as horse-shoe or spherical shaped structures, not larger or 4-times bigger than a procentriole. In all cases, they bear very few or no centriole, suggesting that the size of young deuterosomes does not correlate with the number of centrioles they associate with. Consistently, size and number of deuterosomes are not changed when PLK4 is depleted and centrioles absent (LoMastro et al., 2022). Further live imaging suggests that small structures become spherical and grow over A-stage progression, and large structures resize into smaller entities. We still don’t know which mechanisms (fusion, accretion, splitting, etc.) drive (i) the homogenization of deuterosome size/shape and (ii) the scaling of deuterosome size with centriole number, both observed when arriving at G-stage.

Our data suggest that the pericentrosomal nest is a particular micro-environment characterized by the convergence of MT, the accumulation of centriolar satellites, a concentration of centriole and deuterosome proteins (and maybe mRNA), and the emergence of multiple new organelles. Whether this leads to regional changes in the rheology of the cytoplasm which could facilitate/stabilize the formation of deuterosomes and centrioles is challenging but would be worth studying (Delarue et al., 2018; Fernandes-Mariano et al., 2025; Kalbfuss & Gönczy, 2023).

Interestingly, our live imaging identifies that mRuby-DEUP1+ and CEN2-GFP+ foci (or deuterosomes) are moving with stochastic back and forth fluctuations to the centrosome in a MT-and dynein dependent manner. Random change of cargoes direction alternating between runs and very long pauses exists in numerous cellular contexts. They are driven by complex mechanisms ranging from the binding of cargoes to dyneins and kinesins which can compete or activate each-others to the detachment of motor-cargoes complexes from MT due to their interaction with the surroundings (Hancock, 2014; Shen et al., 2025). Alternative to a direct transport of protein complexes by molecular motors, minus-end directed streaming flows were shown to be produced by the motion of dyneins on MT, and could indirectly drag, or contribute to drag, protein complexes toward the centrosome (Shinar et al., 2011). Consistent with a passive viscous drag of pericentrosomal material, the CEN2-GFP+ aggregates, frequently present in cells over-expressing CEN2-GFP, can also be attracted to the centrosome and subjected to back and forth movements (see Video 4).

### Mother/daughter asymmetries at the centrosome during centriole amplification

Deuterosomes are frequently seen connected to the daughter centriole during A-stage (Al Jord et al., 2014; Khoury Damaa et al., 2025). Live imaging mRuby-DEUP1 now suggests that this connection can last for hours and be marked by back and forth fluctuations (Fig. 2 Supplementary 1C, E, Video 2, 6, 7). In addition to DEUP1, PLK4 and SAS6 also accumulate to the daughter centriole both in WT and DEUP1 KO cells (Al Jord et al., 2014; Mercey, Levine, et al., 2019, this study). In DEUP1-KO cells, this is accompanied by a larger number of centrioles growing on the daughter versus mother centriole (Mercey, Levine, et al., 2019). This asymmetry has never been explained and was not altered when primary cilium growth was inhibited (Al Jord et al., 2014). Given the back and forth fluctuations we observe to one centrosomal centriole, one could consider the existence of asymmetric hydrodynamic forces at the mother versus daughter centriole.

Studying flow fields at the convergence of MT is a technical challenge. Outside cell biological contexts and for engineering purposes, flow fields existing around macroscopic or microscopic (50µm) cylinders and subjected to a directional flow are studied. It reveals the existence of horse-shoe vortical / mixing flow structures upstream of the cylinder, which can lead to horse-shoe shaped material sedimentation at the bottom of the cylinder (Cuéllar et al., 2009; Dargahi, 1989; Ozawa et al., 2023). One could make a parallel with the horseshoe deuterosomes we frequently observe in the pericentrosomal region or connected to the proximal part of the daughter centriole. Since the daughter centriole does not nucleate MT at the beginning of A-stage, it represents a microscopic cylinder subjected to a directional flow toward the mother centriole. Live imaging mRuby-DEUP1 now suggests that the connection of deuterosomes to the daughter centriole can last for hours, explaining the frequency of previous observations, and be marked by back and forth fluctuations (Fig. 2 Supplementary 1C, E, Video 2, 6, 7). Although the viscosity of the cytoplasm is much higher than the viscosity of water solutions, it is tempting to hypothesize that some vortical forces would contribute to the “sedimentation” of DEUP1 at the bottom of the daughter centriole (Al Jord et al., 2014; Khoury Damaa et al., 2025). Such asymmetric hydrodynamic contexts could explain why deuterosomes are never seen connected to the mother centriole, and seem mostly forming on the side of the daughter centriole opposite to the mother centriole. Such forces could also lead to the sedimentation of other proteins such as PLK4 and SAS6 (Al Jord et al., 2014; Mercey, Levine, et al., 2019). This mother/daughter asymmetry which does not seem to have a function in MCC progenitors (Mercey, Al Jord, et al., 2019), may be worth studying as it could have implications on both the control of centriole number and cell fate decision (Tozer et al., 2017).

### Physical property/ies of DEUP1-composed structures

Using live imaging, we show that DEUP1 exists first as a dissolved state organized in a pericentrosomal cloud during A-stage, then, form deuterosomes which eventually split and dissolve in D-stage. They can reform spherical dense aggregates before definitively disappear in the terminally differentiated MCC. Since mature deuterosomes are spherical membrane-less organelles, it would have been appealing to explain their formation through liquid-liquid phase separation. However, evidences accumulate in favor of a solid-like state: (i) DEUP1 first accumulates as a dissolved -yet concentrated- state in a PCM-like manner (ii) we and other did FRAP experiments on mature centriole-loaded deuterosomes composed of exogenous or endogenously tagged DEUP1 and did not observe recovery (Fig. 8 Supplementary 2; (Yamamoto et al., 2021)), (iii) we never observed fusion of mature G-stage deuterosomes, that frequently bump into each other, in the tens of different movies we analyzed (Video 30), (iv) consistently, the number of deuterosomes seems stable during G-stage as previously quantified in single cell live experiments (Mercey, Al Jord, et al., 2019). These evidences are consistent with the hypothesis of Sorokin in 1968 proposing that deuterosome results in solid crystallization of a super-saturated solution of deuterosome components (Sorokin, 1968). However, before deuterosomes disappear during D-stage, they frequently merge or split in/into amorphous structures (Fig. 6G), suggesting a change in the physical properties of the organelle between G- and D-stage, in favor a liquid-like state. Such change could facilitate deuterosome dismantlement and could result from post-translational modifications during the APC/C dependent G-to-D transition (Al Jord et al., 2017). Given that disengagement is dynein and MT dependent, the change in physical properties may also involve a dynein dependent transport of molecular actors such as separase.

The debate is intense for identifying the physical properties of the centrosome, which is massively studied organelle (Raff, 2019). We will probably have to wait better tools to assess this question for the different stages of deuterosome structures, keeping in mind that a cellular structure may probably not behave like any of the solid or liquid ideal state.

### A 2-in-1 centriole biogenesis cycle

Using CEN2-GFP live imaging and immunostainings, we previously described that procentrioles forms sequentially during A-stage, and accumulate in a latency immature state until they synchronously acquire POC5 and get MT wall polyglutamylated (GT335+) during G-stage. CLEM further showed that this A-to-G transition was also marked by an increase in procentriole diameter, accompanied by the completion and lengthening of MT walls (Al Jord et al., 2014). Such widening has been also recently described, and called blooming, in cycling cells (Laporte et al., 2024). Now, the use of UExM and the identification of a spatio-temporal progression of centriole biogenesis, allowed us to further decompose the events occurring during A-stage. In the earliest stages, where only a small pericentrosomal nest is present, PLK4 and SAS6 accumulate at the parent centrioles and inside the nest. Later on, nascent PLK4+/SAS6+ procentrioles deprived of a microtubule wall grow on (or associate with) small DEUP1+ structures in the vicinity of the centrosome (0-3um). Such “naked” cartwheel assembly was also recently revealed as the earliest step of centriole biogenesis during centriole duplication in cycling cells (Laporte et al., 2024). Then, MT walls begins to appear and lengthen as procentriole migrate away from the nest, to reach a plateau at around 150 nm. This lengthening of MT walls is accompanied by a widening of the procentriole diameter that grows from 120 nm to reach a plateau at around 220 nm. The range and correlation of procentriole length and width increases are consistent with what was recently described in cycling cells during S-stage (Laporte et al., 2024). Then, during radial migration and elongation, MT walls progressively acetylate. As previously documented, the A-to-G transition is then marked by lengthening of procentrioles and further widening which falls in the range recently documented for G2 stage procentrioles and is consistent with the documented acquisition of POC5 – a G2 protein also acquired during G-stage – when procentrioles exceed 160 nm in length (Laporte et al., 2024). Finally, procentrioles, which had reached their final width during G-stage, continue to elongate during D-stage and reach a length of typical M-stage procentrioles (Al Jord et al., 2014; Laporte et al., 2024). In early MCC stage, SAS6 is degraded, as during late M-stage (Al Jord et al., 2014). Altogether, this shows that in the MCC cell cycle variant, the formation of centriole barrels - e.g. core molecular architecture, MT wall acquisition, blooming and elongation- follows the same step-wise dynamics, correlated with cell cycle stages, as in the canonical cell cycle (Fig. 9).

**Figure 9.**
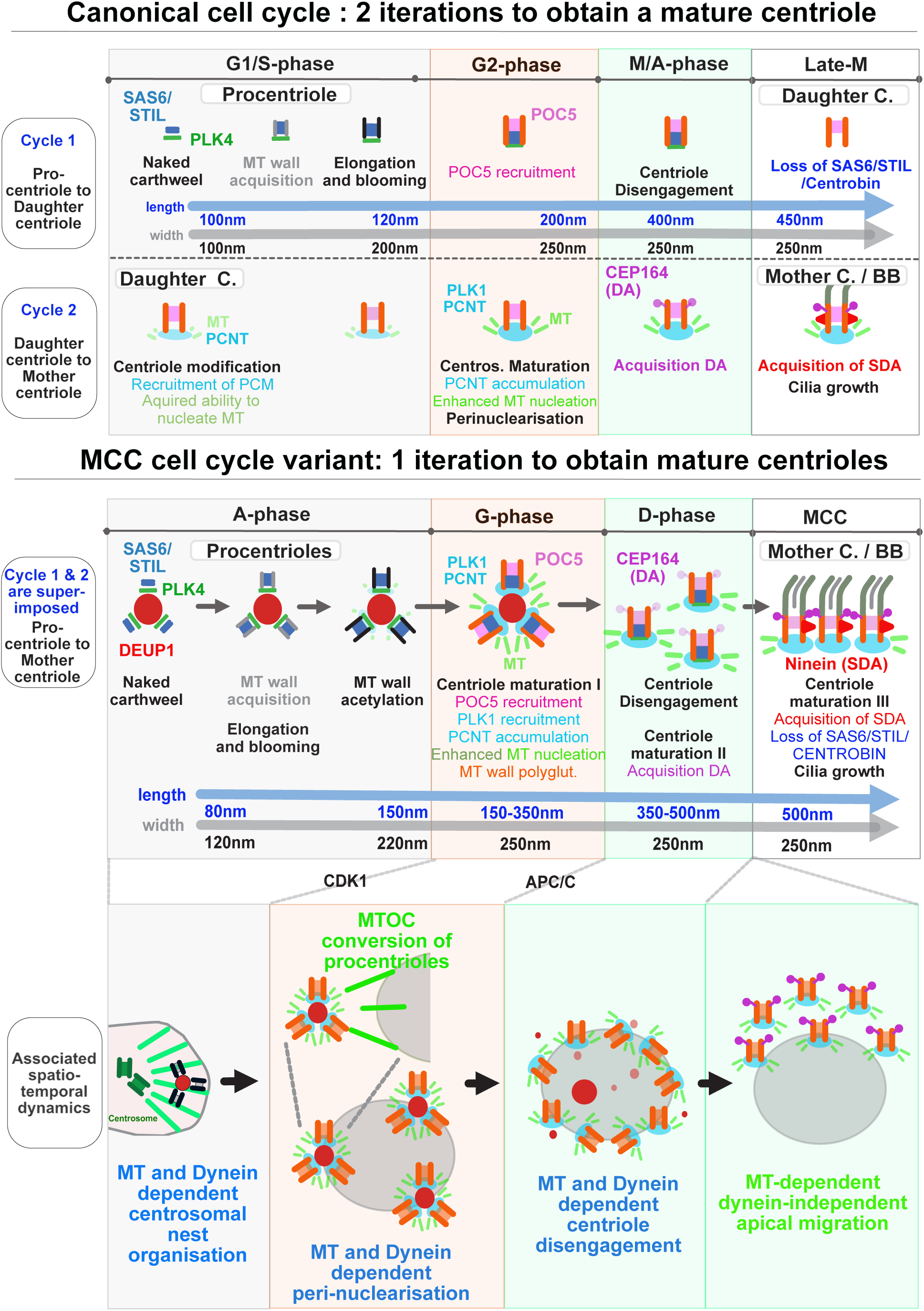
The MCC cell cycle variant incorporates a 2-in-1 centriole biogenesis cycle. Parallel between centriole biogenesis during the canonical and the MCC cell cycle variants showing that centriole elongation and maturation cycles are conserved but superimposed during the MCC cell cycle. The “daughter centriole” step is skipped and prematurely mature centrioles participate in their own dynamics of sub-cellular organization disengagement and migration. DA: distal appendages, SDA: sub-distal appendages, MTOC: microtubule organizing center.

Importantly, using immuno-stainings, single cell RNA sequencing and MT regrowth experiments, we found that the acquisition of maturity features, such as enhanced MT nucleation capacities and centriole appendage recruitment, also follows a canonical step-wise dynamics correlated with cell cycle stages. However instead of occurring over two successive cycle iterations, as in the canonical cell cycle, maturation occurs on elongating procentrioles within a single iteration of the MCC cell cycle variant (Fig. 9). As a consequence, the “daughter centriole stage” is skipped for the amplified procentrioles. Their maturation is superposed with their own elongation, and is nearly concomitant with the maturation of the MCC daughter centriole (Fig. 9 Supplementary 1). Since centriole elongation and maturation in consecutive cell cycles are controlled by some common key players (CDK1, PLK1, APC/C), one can hypothesize that a molecular break is just lifted in MCC to accelerate centriole biogenesis by allowing elongation and maturation to occur concomitantly on the same centriolar structures. Such maturation of nascent procentrioles over a single iteration of the canonical cell cycle has been experimentally obtained by enhanced PLK1 activity (Kong et al., 2014). This suggests that the decomposition of centriole biogenesis in two cycle iterations is not mandatory, and may have been adopted in dividing cells to ensure the growth of a solitary primary cilium.

### Precocious maturation of procentrioles allows their spatial self-organization and disengagement

The superposition of centriole elongation and maturation allows procentrioles to become strong MT nucleators from G-stage. This is accompanied by a dynein-dependent spatial switch in procentriole organization, where the focalized emergence of immature procentriole-loaded deuterosomes, characteristic of A-stage, is followed by their migration and perinuclear distribution around the nuclear membrane during G-stage. This migration and physical distancing of centrosomes and deuterosomes reminds the dynein-dependent nuclear migration of new centrosomes at mitosis onset and their consecutive separation along the nuclear membrane (Agircan et al., 2014). Since A-to-G and G2-M transitions are regulated by PLK1 and CDK1, one can suppose that the same mechanisms are at work in both cycling cells and brain MCC. Consistently, core players involved in centrosome maturation (PLK1, AURKA, CEP192, PCNT), dynein-dependent centrosome-nuclear migration (BICD2, CenpF) and centrosome dysjunction (TPX2, AurA, Nek2A-6-7-9, Mst2) are expressed with comparable dynamics in the canonical and MCC cell cycle variants (Agircan et al., 2014) (Fig. 5 Supplementary 1C).

As during centriole duplication, this perinuclearisation is followed by centriole disengagement, and is known to be PLK1 and APC/C dependent in both canonical and MCC cell cycle variants (Al Jord et al., 2017; Kim et al., 2022; Ruiz García et al., 2019). However, by contrast to the cell cycle where the nuclear disassembly precedes centriole disengagement, in MCC progenitor, the disengagement is tightly correlated with nuclear association. First, the migration of procentriole loaded deuterosomes systematically precedes centriole disengagement. Second, disengagement proceeds through centriole separation from deuterosomes and deuterosome splitting along the nuclear membrane. Third, disengagement is followed by the migration of individualized procentrioles along the nuclear membrane which frequently results in their isotropic distribution before nuclear detachment. Finally, both microtubule depolymerisation and dynein inhibition detach disengaging centrioles from the nucleus and hinders centriole disengagement. Interestingly, a strong effect of diazepam on centriole disengagement from deuterosomes in quail MCC progenitors was shown long ago, and diazepam was later shown to alter MT nucleation (Boisvieux-Ulrich et al., 1987; Spurck & Pickett-Heaps, 1994). Altogether this strongly suggests that the same mechanical forces that pulled procentrioles to the nuclear membrane, also assist their disengagement. One could therefore hypothesize that, once the molecular switch of the mitotic oscillator is ‘on’, the splitting and/or separation of deuterosomes and centrioles are assisted by dynein-dependent microtubule mechanical forces that pull centrioles away from each other’s.

### A complex baso-apical migration dynamics

Disengagement around the nuclear membrane is followed by the migration of centrioles to the apical cortex to dock and nucleate cilia. Characterizing this process has never been done and is very challenging since individual centrioles move in close proximity with tens of other centrioles. Following individual trajectories requires high resolution spatio-temporal live imaging while avoiding excessive light exposure which disturbs centriole migration (Boudjema et al., 2024). Our trackings reveal a global apical convergence of new centrioles around the former centrosome. Using high temporal resolution microscopy, we further identify that individual dynamics is complex and can be split at least between the baso-apical migration of centrioles and their motion once they have reach the apical side of the nucleus. The baso-apical migration is processive and more rapid than the apical motion, marked by alternance between back and forth movements and between diffusive-like and processive steps. Such intermittent movement is probably multifactorial. It may result from (i) the resistance of a crowded environment on a massive population of large organelles that congregate (Vale, 1985), (ii) the binding and release of centrioles by molecular motors (Ananthanarayanan et al., 2013), (iii) binding to molecular motors going in opposite directions (Hancock, 2014) (iv) a combined action of microtubules with the acto-myosin network which is known to be concentrated apically in terminally differentiated MCC and involved in the gathering of basal bodies in brain MCC (Al Jord et al., 2014; Hirota et al., 2010; Kunimoto et al., 2012; Lemullois et al., 1988), and (v) an asynchronized docking of basal bodies nucleating cilia, leading to their association with the nuclear membrane.

The rate of pure baso-apical migration we quantify is in the range of the microtubule-dependent centriole migration described in multiciliated mouse olfactory neurons (Ching et al., 2022). It seems also comparable to the more extensively characterized dynamics of centrosome migration for primary ciliogenesis in serum-starved RPE1 cells (Pitaval et al., 2017). In these cells, even if the morphology is standardized by micro-patterning, apical migration duration is highly variable and stands in the same range (0,03-0,3 µm/min) as what we describe here (Fig. 8F). Interestingly, in this case also, the migration is dependent on MT but independent from dyneins, probably explaining why centriole migration for ciliogenesis is 2 order of magnitude slower than centrosome repositioning at the T-lymphocyte synapse, where it relies on dynein pulling forces (Yi et al., 2013). It now remains to study whether multiple centriole apical migration in MCC relies on the same mechanisms as centrosome migration for primary ciliogenesis, e.g. pushing forces exerted by the stabilization of MT polymerized from the newly formed centrioles, combined with actin contractions. Several old and more recent studies on different organisms (ctenophore *Beroë*, *Xenopus* and quail) suggest the involvement of actin in final centriole migration in MCC (Boisvieux-Ulrich et al., 1990; Ioannou et al., 2013; Tamm & Tamm, 1988).

## Supporting information

Supplemental material with figures

**See supplementary materials for methods, 20 supplementary figures and their captions and 28 movie captions. Video files are available on:** https://hub.bio.ens.psl.eu/index.php/s/NnQS5PEGJXDTTok

## Acknowledgements

We thank all the members of the Spassky lab who contributed in the elaboration of this research work, Chris Kintner for sharing the adenoviral vectors for the expression of MCIDAS-E2F4-Dp1 and Xavier Morin for sharing the MEFs cells. We are thankful to the IBENS administrative team and imaging platform for their support, and to the IBENS Animal Facility for animal care. We thank Camille Noûs, a collective of researchers, who encouraged us to maintain open, honest and slow science (https://www.cogitamus.fr/indexen.html). The team was funded by the Agence Nationale de la Recherche Investissements d’Avenir (ANR-10-LABX-54 MEMO LIFE and ANR-11-IDEX-0001-02 PSL Research University). The team is supported by Inserm, the CNRS, the École Normale Supérieure (ENS), the ANR (ANR-19-CE13-0027, ANR-20-CE45-0019, ANR-21-CE16-0016, and ANR-22-CE16-0011), PSL (Q-Life program), the Fondation pour la Recherche Médicale (FRM EQU202103012767), and the European Research Council (ERC Consolidator grant 647466). A-R.B received a fellowship from La ligue and the Labex MEMOLIFE. A. AJ is supported by the Spanish Ministry of Science and Innovation through the Agencia Estatal de Investigación grant PID2024-157329NA-I00, the Centro de Excelencia Severo Ochoa (CEX2020-001049-S, MCIN/AEI /10.13039/501100011033), and by the Generalitat de Catalunya CERCA programme. This work was also funded by the Damon Runyon Cancer Research Foundation (C. E. J is a Merck Fellow DRG-2478-22); a Hartwell Foundation Fellowship to C. E. J; R01GM133897, R01GM114119, R01CA266199 to A.. J. H.

## Author contributions

A-R. B, R. B. designed, performed and analyzed the majority of the experiments, with C. E. J performing all the U-ExM and FRAP experiments, O. M. providing the first data on the role of microtubules in centriole amplification, A. A. J. having performed some of the movies analyzed in this study and M. F. providing general cell culture support. G. L. M., C. E. J. and A. J. H. designed, produced and validated the PLK4 antibody and the mRuby-DEUP1 mice. All the experiments of MEFs on micropatterns were performed in the lab of M.T with the support of A. S. A-R. B., R. B. C. E. J, N. D., N. S., A. J. H. and A.M. analyzed the data. C.N is a collective of researchers, who encouraged A.M. to maintain open, honest and slow science. A. M. conceived and supervised the study. A-R. B., R. B, C. E. J., A. M. co-wrote the manuscript.

