## Supplemental material with figures for "The cell cycle variant in multiciliated cells incorporates 2 centriole biogenesis cycles"

### Supplementary materials

- Methods and associated references
- 20 Supplementary figure legends
- 20 supplementary figures: https://hub.bio.ens.psl.eu/index.php/s/DeX2QwoAgGHKtTy
- 28 Movie legends
- Video files: https://hub.bio.ens.psl.eu/index.php/s/NnQS5PEGJXDTTok

### Methods

#### Animals

All animal studies were performed in accordance with the guidelines of the European Community and French Ministry of Agriculture and were approved by the Direction départementale de la protection des populations de Paris (Approval number APAFIS#9343-201702211706561 v7). Animal experiments were approved by the Johns Hopkins University Institute Animal Care and Use Committee (MO21M300). Both male and female pups were used for experiments. The CEN2-GFP (CB6-Tg(CAG-EGFP/CETN2) 3-4Jgg/J, The Jackson Laboratory) mice used in this study have already been described (Al Jord et al., 2014).

**Generation of mRuby-DEUP1 mice:** mRuby-DEUP1 knock-in mice were generated using CRISPR-Cas9 genome editing. A 908 base pair double-stranded DNA donor template containing mRuby3-mAID with homology arms to DEUP1 was created by PCR amplification from a vector template. The DNA donor template, along with crRNA, tracrRNA, and Cas9 ribonucleoprotein complexes were injected into B6SJLF1/J mouse zygotes and transplanted into pseudopregnant females. Microinjection and transplantation were performed by the Johns Hopkins Transgenic Core. F0 progeny were genotyped and confirmed knock-ins were outcrossed to C57BL/6J lines. These F1 offspring were again confirmed by genotyping and outcrossed at least twice before using for experiments. sgRNAs were designed using (http://crispor.tefor.net). The crRNA guide was synthesized by IDT to the following sequence: GATGTAGACATTTCTTGGCA. The DEUP1 5' homology arm sequence: CAAGAAGTACCAAGCCAGATGTAGACATTTCTTG. The DEUP1 3' homology arm sequence: GAGAACCAAGCCCATACCACAGCAGGGTAAGTGCTTCAGAA.CEN2-GFP/mRuby3-DEUP1 were obtained by crossing CEN2GFP with mRuby3-DEUP1 mice.

#### Primary ependymal cell cultures

Newborn mice (P0–P2) were killed by decapitation. The brains were dissected in Hank's solution (10% HBSS, 5% HEPES, 5% sodium bicarbonate, 1% penicillin/streptomycin (P/S) in pure water) and the extracted ventricular walls were cut manually into pieces, followed by enzymatic digestion (DMEM glutamax, 33% papain (Worthington 3126), 17% DNase at 10 mg ml−1, 42% cysteine at 12 mg ml−1) for 45 min at 37 °C in a humidified 5% $CO_2$

incubator. Digestion was stopped by addition of a solution of trypsin inhibitors (Leibovitz Medium L15, 10% ovomucoid at 1 mg ml−1, 2% DNase at 10 mg ml−1). The cells were then washed in L15 and resuspended in DMEM glutamax supplemented with 10% fetal bovine serum (FBS) and 1% P/S in a 1X Poly-l-lysine (PLL)-coated flask. Ependymal progenitors proliferated for 5 days until confluence followed by shaking (250 rpm) overnight. Cells were replated at a density of $1,8.10^5$ cells/20µL (corresponding to days in vitro (DIV) −1) in DMEM glutamax, 10% FBS, 1% P/S on 1X PLL-coated coverslides for immunocytochemistry experiments, Lab-Tek chambered coverglasses (Dütscher) for time-lapse experiments. The day after, the medium was then replaced by serum-free DMEM glutamax 1% P/S, to trigger ependymal differentiation in vitro (DIV 0). The protocol is detailed in (Delgehyr et al., 2015).

### MEF-MCC cell culture and viral infections

#### Cell culture

Primary MEFs were grown in DMEM supplemented with 10% FBS and 0.1 mM non-essential amino acid (#11140-050; invitrogen) in a 37 °C humidified incubator with 5% CO2. Only MEFs maintained in culture for less than 4 passages were used in all experiments reported here.

#### Adenovirus infection process (Multicilin/E2f4VP16)

Plasmids for adenoviruses (Kim et al., 2018) were a gift of C. Kintner. Adenoviral vectors expressing MCIDAS/E2f4/VP16 were generated by the JHU plaform and VVTG platform at Necker Institute. Primary MEFs were plated on 12 mm coverslips or proper size of cell dishes with 10% FBS-DMEM for overnight then serum-starved with 0.5% FBS-DMEM for 1 d before adenovirus infection. Adenovirus infection was performed by adding the adenovirus crude lysate to serum-starved MEF cells (500 MOI) for four hours and shaking every hour for optimal infection. Washing was performed twice with pre-warmed 0.5% FBS-DMEM, and infected cells were grown under serum-starved condition with 0.5% FBS-DMEM. The cells were fixed and subjected to analysis according to days post infection (DPI).

#### Micropatterning MEF-MCC

The protocol used is described in (Gazzola et al., 2022). Two days before seeding, MEF cells were thawed and plated on plastic dishes. On seeding day, cells were plated on crossbow shape (3.500 µm²) micropatterned coverslips in 6-well plate as -2 DPI. Then, the cells were grown under the same protocol as described after adenovirus infection.

### Drug treatments

**Concentrations/efficiency:**

A range of nocodazole and dynapyrazole treatment conditions was tested. Efficiency of MT depolymerization after nocodazole treatments were evaluated by immunostaining alpha tubulin (DM1A antibody, Fig. 4 Supplementary 1A). We mainly used protocoles where MT are perturbed but not entirely depolymerized, allowing centrioles to be produced. One has to note that MT in MCC are very stable and difficult to depolymerize. Efficiency of dynein inhibition after dynapyrazole treatments was tested by assessing Golgi apparatus fragmentation/dispersion (Fig. 4 Supplementary 1B). Because of the low stability of the drug, we tested dynapyrazole efficiency for each single experiment (live or chronic).

**Protocols:**

Chronic treatments were done to test the overall efficiency of centriole amplification when MT and dyneins are perturbed, while acute treatments were done to test the role of MT and dyneins at each stage of amplification (A-amplification, G-growth, D-disengagement, M-migration). More specifically, chronic treatments (i.e. 24h-48h for nocodazole and 16h for dynapyrazole which is more damaging to the cells) were followed by methanol or PFA fixation and immunostainings or electron microscopy to assess the consequences of MT depolymerization and dynein inhibition on deuterosome size, deuterosome shape, deuterosome number, centriole number, and the analysis of centriole amplification stages. Acute treatments of 4h were used to assess the consequences of MT depolymerization and dynein inhibition on the migration of centrioles to the nuclear membrane. Live treatments were done to assess the consequences of MT depolymerization and dynein inhibition on the dynamics of the different stages of centriole amplification/migration, deuterosome formation and DEUP1 foci oscillations. Except for oscillations were the “dt” used is too short, the drugs were added after the first imaging time point.

**Regrowth experiments**

Regrowth experiments were performed by adding Nocodazole 10 µM during 24h between DIV2 and DIV3 in a 37 °C humidified incubator with 5% $CO_2$, then washed out with DMEM

0% FBS, 37°C. Microtubules were allowed to regrow for 0 to 5 min after which the cells were fixed with Methanol for immunostainings.

**Washout experiments**

Washout experiments were performed by adding Nocodazole 10 µM during 24h between DIV2 and DIV3 in a 37 °C humidified incubator with 5% $CO_2$, then washed out with DMEM 0% FBS, 37°C for 4h after which the cells were fixed with Methanol for immunostainings.

| | **Acute treatments** | **Chronic treatments** |
|---|---|---|
| Nocodazole (Merck ; M1404-2MG) | Live, 4h & 24h<br>1 µM and 10 µM | 48h<br>1 µM and 5 µM |
| Dynapyrazole-A (Sigma ; SML2127) | 4h (3 µM), Live dynamics of amplification (10h), 3 µM<br>Live imaging of oscillations (0.25-2h)<br>7.5 µM | 16h<br>3 µM |

### Immunostainings

Cell cultures were fixed between DIV2 and DIV5 for brain MCC, and DPI0 and 2 for MEF-MCCs, in methanol at −20 °C for 10 min. Samples were pre-blocked in 1× PBS with 0.2% Triton X-100 and 10% FBS before incubation with primary and secondary antibodies. Cells were counterstained with Hoechst (10 µg ml−1, Sigma) and mounted in Fluoromount (Southern Biotech). The following antibodies were used: rabbit anti-DEUP1 (1:5000, homemade, raised against the peptide TKLKQSRHI); mouse IgG1 anti-GT335 (1:500, Adipogen) ; mouse anti-DM1A (1:500, Sigma) ; mouse IgG2b anti-SAS6 (1:750, Santa

Cruz) ; rabbit anti-Pericentrin (1:2000, Covance) ; rat anti-YL1/2 (1:500, Abcam); mouse anti-MAB414 (1:500; Eurogentech); mouse IgG2b anti-PLK1 (1:1000, Abcam) and species-specific Alexa Fluor secondary antibodies (1:400, Invitrogen).

### PLK4 Antibody Generation

Mouse PLK4 spanning amino acids 475-925 was Gibson cloned into a pET-6xHis expression vector with an N-terminal tag using the following primers:
Forward primer: ACCATCATCACAGTAGCCATATGCAGGACACGTTACGAAACG and

Reverse primer:
GTGGTGGTGCTCGAGTTACTACTGAAAATTAGGAGTTGGATTAGAAAACA.

PLK4 was expressed in BL21 Rosetta 2 competent *E. coli* cells. *E. coli* cultures were grown at 37°C to an OD of 0.5-0.6 and then induced with 1mM IPTG for 4 hours at 37°C. Cells were lysed using sonication and then PLK4 protein purification was carried out under denaturing conditions in 8M urea using Ni-NTA beads. PLK4 was cleaved from the 6xHis tag and 2mg of protein was sent to Pocono Rabbit Farm and Laboratory, Inc for immunization in rabbits. The rabbit immune serum was affinity purified. The PLK4 antibody was validated with immunofluorescence in mouse tracheal epithelial cells cultures using a conditional loss-of-function PLK4 allele. See (LoMastro et al., 2022) for PLK4 allele.

### Microscopy

***Epifluorescence microscopy**:* Fixed cells were examined with an upright epifluorescence microscope (Zeiss Axio Observer.Z1) equipped with an Apochromat ×63 (NA 1.4) oil-immersion objective and a Zeiss Apotome with an H/D grid. Images were acquired using Zen with 240-nm or 500-nm z-steps. When better resolution was needed, confocal image stacks were collected with a 63 x/1.4 Oil objective on an inverted LSM 880 Airyscan Zeiss microscope with 440, 515, 560 and 633 laser lines with 160-nm z-steps.

***Videomicroscopy**:* Cultured cells between DIV1 and DIV6 were filmed in vitro using an inverted spinning disk Nikon Ti PFS microscope equipped with oil-immersion ×63 (NA 1.32) and ×100 (NA 1.4) objectives, an Evolve EMCCD Camera (Photometrics), dpss laser (491 nm, 25% intensity, 70-100ms exposition), appropriate filter sets for DAPI/FITC/TRITC, a motorized scanning deck and an incubation chamber (37 °C; 5% CO2; 80% humidity). Images were acquired with Metamorph Nx with 5 s, 15 s, 5 min, 20 min and 40 min time intervals. Image stacks were recorded with a z-step of 0.5 µm. Detailed protocol for live imaging is described in (Boudjema et al., 2024). Drugs were added just after the first acquisition for experiments on the dynamics of amplification (quantification of cloud and deuterosome area during A-stage, nucleus tethering of deuterosome during G-stage, disengagement of centrioles during D-stage, migration of centrioles), or 0.25-2h before acquisition for analysis of oscillations of DEUP1+ structures.

***Ultrastructure Expansion microscopy (U-ExM):*** U-ExM was performed as previously described (Gambarotto et al., 2021) with slight modification. Coverslips with brain MCCs were incubated in PBS with 1.4% formaldehyde and 2% acrylamide overnight at 37 °C. Gel polymerization was then performed in a humid chamber on ice by placing coverslips face down on top of a 50 µl drop MS solution composed of 23% (w/v) sodium acrylate, 10% (w/v) acrylamide, 0.1% (w/v) N,N′-methylenbisacrylamide (BIS), 0.5% APS, and 0.5% TEMED in PBS. Samples were allowed to harden for 20 min on ice and then placed at 37 °C for 1 hour to complete polymerization. Coverslips were turned face up and a 4 mm biopsy puncher (Integra; 33–34 P/25) was used to create one punch per coverslip. Each punch was transferred into a 1.5 ml Eppendorf tube containing 1 ml of denaturation buffer containing 200 mM SDS, 200 mM NaCl, and 50 mM Tris in water, pH 9, and incubated at 95 °C for 1h. Gel punches were then moved to beakers filled with water at room temperature for the first expansion. Water was exchanged twice after 30 min, and then gels were incubated overnight in water. Gels were then washed twice in PBS for 10 min before blocking in PBS with 2% BSA for 30 min-1 hour at 37 °C with gentle agitation. Gels were moved to primary antibody solution in PBS with 2% BSA overnight at 37 °C with gentle agitation. Gels were washed in PBS with 0.1% Tween 3x for 10 min each and then incubated with secondary antibody solution in PBS with 2% BSA for 2.5 hours at 37 °C with gentle agitation. Gels were again washed in PBS with 0.1% Tween 3x for 10 min each. Gels were moved to water for the second round of expansion. Water was exchanged twice after 30 min, and then gels were incubated overnight in water. Gels expansion was consistently around 4.0-4.1x.

To image gels, punches were placed on uncoated 35 mm glass bottom dish and excess water was blotted away. DAPI staining was used to identify which side of the punch the cells were located. Then, punches were moved to a fresh 35 mm glass bottom dish coated with poly-L lysine and imaged on either a Leica SP8 confocal microscope with a 40x plan-apochromat oil immersion objective with 1.30 NA, or Zeiss Axio Observer 7 inverted microscope with Slidebook 2023 software (3i—Intelligent, Imaging Innovations, Inc.), CSU-W1 X1 (Yokogawa) T1 Super-Resolution SoRa Spinning Disk, and Prime 95B CMOS camera (Teledyne Photometrics) with a 40x plan-apochromat oil immersion objective with 1.30 NA. Images were deconvolved with Leica's lightening process software or Microvolution software built into Slidebook. Images were processed in FIJI (Schindelin et al., 2012). A minimum of three biological replicates were performed for each experiment. All images presented in figures are max projections.

The following primary antibodies were used for U-ExM. Mouse anti-Acetyl-α-Tubulin (Lys40) (6-11B-1) (1 :1000, Cell Signaling #12152), mouse anti-SAS-6 (1 :250, Santa Cruz #81431), rabbit anti-PLK4 (1 :500, homemade), rabbit anti-DEUP1 (1 :1000, homemade), guinea pig anti-β-tubulin/ scFv-S11B (1 :1000, Geneva Antibody Facility #AA344), rabbit anti-Centrin (1 :1000, homemade), mouse anti-PCNT (1:250, BD Transduction Laboratories #611814).

***FRAP (fluorescence recovery after photobleaching):*** MCC progenitors from brains homozygous for EGFP-Centrin1 and mRuby-DEUP1 were plated on 35 mm glass-bottomed dishes. The following day, cells were washed with PBS 1X and the medium replaced with serum free DMEM/Glutamax with 1% penicillin/streptomycin. Cells were allowed to differentiate for 4-8 days prior to imaging. Cells were imaged on a Leica Sp8 inverted microscope using a 63×/1.4 NA objective. A 5 μm region of interest (ROI) was drawn and photobleached at approximately 80% fluorescence intensity before bleaching with 5 pulses of 552 nm laser light. A Z-series of 9 planes (0.5 μm steps) was acquired immediately before and after bleaching for 10 minutes at 1- minute intervals in the 488nm and 552nm channels. Fluorescence intensities of mRuby-DEUP1 were measured using ImageJ. The first post-bleach intensity was subtracted from the pre-bleach and post-bleach values, then normalized to give the percentage of the pre-bleach intensity.

### Quantification and statistics

***Density mapping in MEF-MCC:*** For centrosome stage and early amplification stage in control cells, only cells with apically positioned centrosomes were analyzed. All crossbow MEF-MCCs micrographs were cropped following the same ROI. For centrosome and procentrioles mapping, each point (centrosomal centriole or procentrioles) was pointed with the “multi-point” option on Image J and added to ROI Manager file. For each condition, all points were projected on a single image containing the average intensity of each pixel and intensities were color coded with the “fire” look-up table. For PCNT fluorescence intensity mapping, each micrographs were max-projected on Image J, grouped together using the “concatenate” tool, then stacked on a single image containing the maximal intensity of PCNT and intensities were color coded with the “fire” look-up table.

***Quantifications of oscillations:*** Oscillation behavior was scored on A-stage cells live imaged with a dt=5-15s after 0.25-2h treatments of DMSO, Nocodazole 10 µM or dynapyrazole 7.5 µm. Cells were scored positively if at least one DEUP1+ foci or deuterosome is seen oscillating from and to the centrosome in 1-4 min movies.

***Clouds and deuterosome areas quantifications:*** Surface covered by the CEN2-GFP; mRuby3-DEUP1+ cloud and by CEN2-GFP; mRuby3-DEUP1+ deuterosomes during A-stage were manually measured on image J at each timepoint for 10h. On the x axis, “0” represents the onset of the movies quantified. DMSO, Nocodazole 10 µM or dynapyrazole 3 µM are added just after the first acquisition. dt=40 min; >3 independent experiments were analyzed.

***Deuterosome size quantifications:*** Deuterosome diameter was quantified on ImageJ using the manual tool “Straight line. Each image was set to the same fluorescence intensities.

***Fluorescence intensities quantifications:*** PCNT, PLK1 and YL1/2 (tyrosinated MT) fluorescence intensities on procentrioles in ependymal MCCs were normalized on PCNT and PLK1 and YL1/2 fluorescence intensities of the centrosome of a non-differentiating neighboring cell. For A-stage, measurements were done on ROI (0,57 $\mu m^2$) covering a group of procentriole on adeuterosome, as procentrioles are not identifiable. For G-stage, measurements were done on ROI (0,57 $\mu m^2$) covering each procentriole separately. For regrowth experiments, YL1/2 fluorescence intensities were measured on CEN2-GFP ependymal cells or SAS6-stained MEF-MCCs marked with GT335 to discriminate A- versus G-stage procentrioles. Measurements were done on ROI covering an area of 0,57 $\mu m^2$ around each procentriole-loaded deuterosome in both A- and G-stages. YL1/2 fluorescence intensities were normalized on YL1/2 fluorescence intensity of the centrosome of a non-differentiating neighboring cell.

***Migrating centriole trajectory analysis:*** To analyze the global 3-dimensional displacement of all centrioles in individual cells throughout their apical migration, we used Imaris software to automatically segment all centrioles based on fluorescence quality and particle size threshold filters (0.25 µm diameter), and extracted the *X, Y, Z* coordinates of all centrioles at the end of the disengagement stage and at the end of migration and constriction of the basal body patch. Using ImarisTrack software, we manually tracked centrosomal centrioles to extract their coordinates *Xc, Yc, Zc* at these two stages. As the two centrosomal centrioles remained close to each other throughout the process, we chose one of the centrosomal centrioles indiscriminately to calculate centriole-centrosome distances.

To track individual centriole movements, we decreased the Δt from 40 to 5 minutes. Due to the large number of centrioles and the limited distance between them, Imaris' automatic tracking algorithms produced erroneous tracings. We therefore used the manual option of the Imaris software to track several centrioles as well as the coordinates of one centrosomal centriole. Coordinates of the centrosomal centriole reference point are substracted from the coordinate of migrating centrioles for normalization. The « Directionality » is defined here as the evolution of the distance between the tracked centriole and the centrosomal centriole reference point between each time frame. A positive value means that the centriole is approaching this reference point, and a negative value means that the centriole is moving away from it. The velocity represents the distance covered per minute in relation to the centrosome reference point. The absolute speed is calculated as the distance covered per minute by the centriole since the last time point.

**Distance to the centrosome :**

$$\sqrt{(X-Xc)^2+(Y-Yc)^2+(Z-Zc)^2}$$

**« Directionality » :**

$$(\sqrt{(X-Xc)^2+(Y-Yc)^2+(Z-Zc)^2})t - (\sqrt{(X-Xc)^2+(Y-Yc)^2+(Z-Zc)^2})t+1$$

**« Velocity » :**

(« Directionality ») / 5

**« Speed » :**

$$\sqrt{(Xt+1-Xt)^2+(Yt+1-Yt)^2+(Zt+1-Zt)^2}$$

***Statistical analysis.*** All graphs and statistical analyses were obtained using GraphPad Prism software. Data were obtained from at least three independent experiments unless differently stated, and the results presented as the mean ± s.d unless differently stated. Non-parametric two-tailed Mann–Whitney U-tests were used to compare groups of data unless differently stated. Chi-square tests were performed on R to compare proportions and distributions. For p-values: ns, not significant ; P>0.05 ; * : P≤0.05 ; ** : P≤0.01 ; *** P≤0.001 ; **** P≤0.0001.

#### Materials availability statement

Newly generated materials are available from the corresponding author upon reasonable request and completion of a standard material transfer agreement.

#### Data availability

Previously published scRNAseq data sets were used (Serizay et al., 2025; DOI: 10.1016/j.celrep.2025.116523). Briefly, raw sequencing data have been deposited at the NCBI Gene Expression Omnibus under the accession ID GEO: GSE201773 (GEO; https://www.ncbi.nlm.nih.gov/geo/) and are publicly available as of the date of publication. Accession numbers are listed in the key resources table. Processed scRNA-seq data are also available for interactive investigation using the CELLxGENE platform (collection # 33f48a52-31d8-4cc8-bd00-1e89c659a87f) at https://cellxgene.cziscience.com/collections/33f48a52-31d8-4cc8-bd00-1e89c659a87f as of the date of publication. All original code has been deposited at Zenodo (record #14105247) and is publicly available at https://doi.org/10.5281/zenodo.14105247 as of the date of publication. Any additional information required to reanalyze the data reported in this paper is available from the corresponding author upon request.

### Supplementary Figures Legends

#### Figure 1 supplementary 1

**(A)** Immunostaining profiles of YL1/2, SAS6 and GT335 in MEF-MCCs grown on crossbow micropatterns treated with DMSO and Nocodazole (1 µM and 3 µM). Stainings mark respectively tyrosinated MT, procentrioles and mother and daughter centrosomal centrioles. Scale bar, 10 µm. Insets: magnification of centrosomal centrioles and early procentrioles to compare their spacing under each treatment. Scale bar, 5 µm.

**(B)** Quantification of early procentriole (PC) and centrosomal centriole (CC) distance under DMSO and Nocodazole 3 µM on MEF-MCC crossbow micropatterns. Two independent experiments were scored, n= 7 DMSO cells, n= 7 Noco cells. Error bars represent mean ± SD. **** $P<0.0001$, non-parametric Mann Whitney test.

#### Figure 2 supplementary 1

**(A)** CEN2-GFP; mRuby-DEUP1 dynamics of the new mouse line during centriole amplification. The previously described stages of centriole amplification can be identified with the CEN2-GFP signal (yellow). The procentrioles are seen organizing around the endogenous mRuby3-DEUP1+ (cyan) deuterosomes in A-, G- and D-stages. Scale bar, 5 µm.

**(B)** CEN2-GFP and mRuby-DEUP1 dynamics in the primordial cloud. CEN2-GFP+ and mRuby-DEUP1+ foci are not always colocalized. Yellow dotted circles enclose centrosomal centrioles. Note that DEUP1+ foci can interact persistently with one parental centriole. A single z-plane of 0,5 µm is showed. dt=1h; CEN2-GFP in white, DEUP1 in red. Scale bar, 1 µm. See Video 2.

**(C)** CEN2-GFP and mRuby-DEUP1 dynamics showing a cell beginning A-stage with big deuterosomes that dissolve into smaller ones as A-stage progresses. CEN2-GFP in white, mRuby-DEUP1 in red. Scale bar, 1 µm. Red and white arrowheads show procentriole-loaded deuterosomes. mc= centrosomal mother centriole; dc= centrosomal daughter centriole.

**(D)** Deuterosome oscillating behavior during A-stage. White arrows show deuterosomes back and forth movements towards the centrosome ; blue arrows show a deuterosome moving to the cytoplasm and away from other deuterosomes. CEN2-GFP in cyan, DEUP1 in yellow. A single z-plane of 0,5 µm is filmed to allow high temporal resolution. dt=5s; scale bar, 5 µm. See Video 5.

**(E)** mRuby-DEUP1+ foci (white and red arrowheads) intermittent interaction with one centrosomal centriole (white circles) in CEN2-GFP+ cells. A single z-plane of 0,5 µm is filmed to allow high temporal resolution. dt=5s; scale bar, 1 µm. See Video 7.

**Figure 2 supplementary 2**

Correlative light and electron microscopy of a mRuby-DEUP1+ primordial cloud composed of fibrogranular aggregates (same cell as Fig. 2C). Light microscopy images integrate the fluorescence signal of a z-stack of 240 nm from the apical or more basal portion of the centrosome. Serial electron microscopy z-sections are 70 nm. dc= centrosomal daughter centriole, mc= centrosomal mother centriole.

**Figure 2 supplementary 3**

Correlative light and electron microscopy of a mRuby-DEUP1+ primordial cloud composed of fibrogranular aggregates, procentriole-free and procentriole-loaded deuterosomes. Deuterosomes are either spherical or horse-shoe shaped. Deuterosome 1 is free of procentriole and connected to the daughter centriole. Light microscopy images integrate the fluorescence signal of a z-stack of 500 nm around the centrosome. Serial electron microscopy z-sections are 70 nm. dc= centrosomal daughter centriole, mc= centrosomal mother centriole.

**Figure 2 supplementary 4**

Correlative light and electron microscopy of a mRuby-DEUP1+ primordial cloud composed of fibrogranular aggregates, procentriole-free deuterosomes, procentriole-loaded deuterosomes, and procentriole apparently not connected to a visible deuterosome. Deuterosomes are either spherical, amorphous or horse-shoe shaped. Deuterosome 1 is loaded with one procentriole and connected to the daughter centriole. Serial ultra-thin sections are aligned and shown as a z-stack in the Video 3. Light microscopy images integrate the fluorescence signal of a z-stack of 750 nm around the centrosome. Serial electron microscopy z-sections are 70 nm. dc= centrosomal daughter centriole, mc= centrosomal mother centriole.

**Figure 2 supplementary 5**

Correlative light and electron microscopy of a mRuby-DEUP1+ primordial cloud composed of fibrogranular aggregates, and procentriole-loaded deuterosomes. Deuterosomes are either spherical, amorphous or horse-shoe shaped. Light microscopy images integrate the fluorescence signal of a z-stack of 1.25 µm around the centrosome. Serial electron microscopy z-sections are 70 nm. mc= centrosomal mother centriole.

**Figure 2 supplementary 6**

Correlative light and electron microscopy of a mRuby-DEUP1+ primordial cloud composed of fibrogranular aggregates, and large deuterosomes loaded with only one or two procentrioles. Deuterosomes are spherical. Light microscopy images integrate the fluorescence signal of a z-stack of 750 nm around the centrosome. Serial electron microscopy z-sections are 70 nm. dc= centrosomal daughter centriole, mc= centrosomal mother centriole.

**Figure 3 supplementary 1**

**(A-E)** Low contrast (1x brightness) and High (10x brightness) of U-ExM images, brain MCCs were immunostained with antibodies to β-TUBULIN, CENTRIN, DEUP1, SAS6 and PLK4. In the right panels the brightness is increased 10-fold to allow for visualization of the cloud of DEUP1.

**(F)** Quantitation showing the procentriole distance from parents is shorter in procentrioles without Acetylated-TUBULIN compared to procentrioles with Acetylated-TUBULIN. PLK4 is used as a marker for procentrioles associated with deuterosomes. Graphs show mean ±SD from n=3 cells across 3 different experiments. ****$p<0,0001$ using unpaired t-test.

**(G)** Representative U-ExM images of brain MCCs during early A-stage. Cells were immunostained with antibodies to β-TUBULIN, PCNT and CENTRIN.

**(H)** Quantitation showing procentriole length measured by β-tubulin increases with distance from parents during A-stage. Graphs show mean ±SD from n=10 cells across 3 different experiments.

**(I)** Quantitation showing procentriole width measured by β-tubulin increases with distance from parents during A-stage. Graphs show mean ±SD from n=10 cells across 3 different experiments.

**(J)** Representative U-ExM images of brain MCCs during late A-stage. Boxes denote zoomed in regions on right, with 1-4 labels marking increasing distance from the parent centrioles. Cells were immunostained with antibodies to Acetylated-TUBULIN, β-TUBULIN and CENTRIN. Yellow arrows in (J) indicate procentrioles with non acetylated MT and white arrows point at procentrioles with acetylated MT. mc: mother centriole, dc: daughter centriole.

**Figure 3 supplementary 2**

**(A)** Quantitation comparing the frequency distribution of procentrioles according to distance from parents in early-A stage, late A-stage, and G-stage. Graphs show mean +/-SD from n=10 cells for A-stage, n=5 cells for G-stage, and n=7 cells for D-stage across three different experiments.

**(B)** Quantitation showing an increase in procentriole length measured by β-tubulin in A-stage compared to G-stage and D-stage. Graphs show mean ±SD from n=10 cells for A-stage, n=5 cells for G-stage, and n=7 cells for D-stage across three different experiments.

**(C)** Quantitation showing an increase in procentriole width measured by β-tubulin in A-stage compared to G-stage and D-stage. Graphs show mean ±SD from 10 cells for A-stage, n=5 cells for G-stage, and n=7 cells for D-stage across three different experiments.

**(D)** Quantitation showing procentriole length measured by β-tubulin is similar at all distances from parents during G-stage. Graphs show mean ±SD from n=5 cells across three different experiments.

**(E)** Quantitation showing procentriole width measured by β-tubulin is similar at all distances from parents during G-stage. Graphs show mean ±SD from n=5 cells across three different experiments.

**Figure 4 supplementary 1**

**(A)** Representative immunostainting profile of microtubules in DMSO, Nocodazole acute (4h, 10 µM) and chronic (24h and 48h, 1 µM and 5 µM). DM1A stains for total MT. Square in the right bottom of panels are magnifications of the MT signal of one cell of the field. Scale bar, 10 µm.

**(B)** Representative immunostaining profile of Sas6 in Cen2-GFP brain MCC progenitors treated with DMSO or chronic Nocodazole (from the onset of amplification DIV0 to DIV2 (48h); 1µM and 5µM). Each column represents one stage of centriole amplification. Note that a defect in centrin recruitment can be observed in Nocodazole 5µM. Disengagement under 5µM is very rarely observed and often lack SAS6 as mentionned in the Results. Scale bar, 5µm.

**(C)** Representative immunostaining of the Golgi apparatus in CEN2-GFP cells after treatment with DMSO or Dynapyrazole. Cells were treated for 2h (as for oscillation monitoring) or 16h (as for other live imaging or "chronic treatments") with DMSO or Dynapyrazole (7.5 µM or 3 µM respectively). Golgi disorganization was used as a read-out of dynein inhibition in each experiments. Scale bar, 10 µm.

**Figure 4 supplementary 2**

**(A)** Quantification of the area occupied by the deuterosomes over time in A-stage cells under DMSO and acute Nocodazole 10 µM or Dynapyrazole 3 µM treatments. Black arrow indicates that DMSO and drugs were added right after the first acquisition. Three independent experiments were analyzed, n=12 DMSO cells; n= 15 Noco 10 µM cells; n= 15 Dynap 3 µM cells. dt=40min; ****$p<0,0001$; non-parametric Mann Whitney test; error bars represent mean ± SD. See Video 12, 13 and 14.

**(B)** Quantification of the amplification efficiency in DMSO, Nocodazole (5 µM, 48h) or Dynapyrazole (3 µM, 16h) chronic treatments. Each dot represents the % of

DEUP1+ cells per field. >4 independent experiments were quantified; n= 5076 DMSO cells, n=1492 Nocodazole cells, n=1886 Dynapyrazole cells analyzed; error bars represent mean ± SD. ****p<0.0001; Chi-2 test with Yates' correction.

**(C)** Quantification of the deuterosome number per cell in A and G-stage cells (pooled) under DMSO, Nocodazole (5 µM, 48h) or Dynapyrazole (3 µM, 16h) chronic treatments. Three independent experiments were scored, n=80 cells in DMSO, n=63 cells in Nocodazole and n= 19 cells in Dynapyrazole. Error bars represent min to max ± median. *** P<0.001, **** P<0.0001, non-parametric Mann Whitney test.

**(D)** Quantification of deuterosome size in A and G-stage cells (pooled) under DMSO, Nocodazole (5 µM) and Dynapyrazole (3 µM) chronic treatments (48h). Three independent experiments were scored, $n_{deut}$=448 in DMSO, $n_{deut}$=836 in Nocodazole 5 µM, $n_{deut}$=573 in Dynapyrazole 3 µM. Error bars represent mean ± SD. **** P<0.0001; non-parametric Mann Whitney test.

**(E) Left panel**: **r**epresentative immunostainings of PCM1 in CEN2-GFP expressing differentiating MCC during the A/G stage. Note that PCM1 accumulates as a large cloud around the centrosome (arrowheads) in DMSO cells. This cloud is more diffuse in Dynapyrazole (3 µM, 16h) treated cells. **Right panel:** quantification of PCM1 mean grey value along a 10 µm line centered on the centrosome in DMSO and Dynapyrazole (3 µM, 16h) treated cells. N=3 independent experiments on >30 A/G stage cells. Error bars represent mean ± SD.

**(F)** Representative fluorescence images of mRuby-DEUP1;CEN2-GFP expressing G-stage cells counterstained with Hoescht after DMSO or Dynapyrazole (3 µM, 16h) treatment. Note the small size of deuterosomes and the absence of nuclear contact in Dynapyrazole treated cells.

**Figure 4 supplementary 3**

Correlative light and electron microscopy of a mRuby-DEUP1;CEN2-GFP expressing cell treated with Nocodazole 10 µM (24h) showing scattered mRuby-DEUP1+;CEN2-GFP+ structures in the cytoplasm. These structures either correspond to small deuterosomes loaded with a single procentriole, or to group of fibrogranular aggregates. Note a small deuterosome connected to the centrosomal daughter centriole. Serial electron microscopy z-sections are 50 nm. dc= centrosomal daughter centriole, mc= centrosomal mother centriole.

**Figure 4 supplementary 4**

mRuby-DEUP1;CEN2-GFP+ "inverted flower" profile frequently observed under chronic Nocodazole (10 µM, 24h) along with the underlying deuterosome and procentriole arrangement revealed by correlative light and electron microscopy (CLEM). Light microscopy images integrate the fluorescence signal of a z-stack of 1 µm. Serial electron microscopy z-sections are 70 nm. Scale bar 1 µm.

**Figure 5 supplementary 1**

Expression of set of genes involved in centriole biogenesis (A), daughter centriole maturation (B) and centrosome maturation / migration along the nuclear membrane (C) along the pseudotime of the canonical cell cycle (upper graphs) and MCC cell cycle variant (lower graphs). Black curves represent gene set mean expression in each variant. From the single cell RNA sequencing data set published in Serizay et al., 2025.

**Figure 5 supplementary 2**

**(A)** Representative immuno-reactivity profile of PLK1 andGT335 in A- and G-stages in CEN2-GFP+ brain MCC progenitors. Square surrounds procentrioles shown in the magnified view. Scale bar, 5 µm.

(**-B)** Representative immuno-reactivity profile of PCNT in A- and G-stages in CEN2-GFP+ brain MCC progenitors. Dotted line square surrounds procentrioles shown in the magnified view. Scale bar, 5 µm.

**(C)** Quantification of PCNT intensity on procentrioles in A- and G-stage brain MCCs. Fluorescence intensity was normalized on PCNT intensity of the centrosome of one non-differentiating neighboring cell. Three independent experiments were scored; n=

19 cells; error bars represent mean ± SD. ****p<0,0001; non-parametric Mann Whitney test.

**(D)** Intensity profile of PCNT in A- and G-stage DEUP1 KO brain cells. Intensities are normalized on PCNT intensity of one centrosome of a non-differentiating neighboring cell. Three independent experiments were quantified; n=77 cells analyzed. ****p<0,0001; non-parametric Mann Whitney test.

**(E)** Immunostaining profile of YL1/2 in SAS6 stained MEF-MCCs in A- and G-stage after regrowth experiments (see methods). Arrowheads indicate centrosomal centrioles. Arrows indicate the position of A- or G-stage procentrioles. Scale bar, 10 μm.

**(F)** Quantification of microtubule aster intensity in A-stage and G-stage MEF-MCCs. Fluorescence intensity was normalized on the centrosome of one non-differentiating neighboring cell. Three independent experiments were quantified, n= 46 cells. Error bars represent mean ± SD; ****p<0,0001; non-parametric Mann Whitney test.

**(G)** Quantification of the duration of the interaction between procentriole-loaded deuterosome and the nuclear envelope before D-stage in CEN2-GFP brain MCCs. Four independent experiments quantified, n=75 flower, n=8 cells analyzed. Error bars represent min to max ± median.

**(H)** Magnified view of a CEN2-GFP; mRuby-DEUP1 flower dynamics in G-stage. White arrowheads indicate the procentriole-loaded deuterosomes migrating on the nuclear envelope; dotted white line outlines the nucleus identified by contrasting CEN2-GFP signal. dt=5s, scale bar, 1 μm. See Video also Video 10.

**(I)** Immuno-reactivity of DEUP1 on CEN2-GFP brain MCCs during G-stage, treated acutely with DMSO or Nocodazole (10 μM, 4h). The nucleus is counter-stained with Hoechst. Scale bar, 5 μm.

**(J)** Quantification of the % of tethered procentriole-loaded deuterosomes per G-stage cell in Nocodazole (10 μM) and washout condition (Nocodazole 10 μM followed by a 4h washout with Nocodazole-free medium). Three independent experiments were quantified. N=26 cells in Nocodazole and n=42 cells after washout. Error bars represent mean ±SD; ****p<0,0001; non-parametric Mann Whitney test.

**(K)** CEN2-GFP+ procentriole-loaded deuterosome interaction with the nucleus in G-stage cells treated with DMSO or acute Nocodazole 10 µM. At t-1, cells display perinuclearized procentriole-loaded deuterosomes and are not treated with any drug. DMSO and Nocodazole were added right after the first timepoint acquisition. In DMSO, "centriole-loaded deuterosomes" keep their perinuclear distribution (t+45 min). In 10 µM Nocodazole, the "centriole-loaded deuterosomes" start untethering from the nucleus (+15 min), the centrosome also slides towards the edge of the nucleus while most of the centriole-loaded deuterosomes are already untethered (+30 min), until most of the "centriole-loaded deuterosomes" are detached from the nuclear membrane (+45 min). Blue lined circle outlines the shadow of the nucleus identified with contrasting CEN2-GFP. dt=5 min; scale bar, 5 µM. See also Video NEW16.

**Figure 6 supplementary 1**

**(A)** CEN2-GFP+ dynamics from G-stage to migration stage. Top line: time-lapse images showing perinuclear distribution of centrioles at G-stage (00:00); perinuclear disengagement of centrioles from deuterosomes (empty flowers) and parental centrioles (plain flowers, arrows) in a CEN2-GFP+ cell (4:30); release of centrioles from the nuclear membrane (6:30) for apical migration (from 10:00). Bottom line: profile view of the CEN2-GFP+ signal dynamics. Dotted white line outlines the nucleus identified by contrasting CEN2-GFP signal. dt=30min, scale bar, 5 µm. See See also Video 17.

**(B)** Single z-slice of 0.25 µm of CEN2-GFP expressing cells prior (1) or immediately after (2) disengagement show that procentrioles in G- and D- stages are at the same z-level than nuclear pore complex marked with MAB414. Scale bars, 5 µm. See also video 18.

**(C)** Still images of a time-lapse on CEN2-GFP;mRuby-DEUP1 expressing cells showing that DEUP1 can form big aggregates under the basal body patch at the end of amplification, that finally dissolve. Scale bar, 1 µm.

**(D)** Time lapse images of a D-stage cell showing the deuterosome mRuby-DEUP1+ signal dissolution correlated with CEN2-GFP rosette dismantlement. dt=40 min, scale bar, 1 µm. See Video 21 for the entire movie.

**Figure 7 supplementary 1**

**(A)** CEN2-GFP cell during D-stage after acute Nocodazole 10 µM treatment. Nocodazole is added after the first time point. Individualizing centrioles are seen detaching from the nuclear membrane within minutes (00:20). dt=20 min; scale bar, 5 µm.

**(B)** Immuno-reactivity profiles of SAS6 and DEUP1 in CEN2-GFP+ brain MCCs treated with DMSO and Nocodazole (48h, 1 µM) showing $SAS6^{-}/DEUP1^{+}$ structures. White arrow heads show $SAS6^{-}/DEUP1^{+}$ centrioles. Scale bar, 5 µm.

**(C)** Quantification of the proportion of D-stage brain MCCs treated with DMSO or chronic Nocodazole (48h, 1 µM) displaying unusual $SAS6^{-}/DEUP1^{+}$ centrioles. Three independent experiments were quantified, n=40 DMSO cells, n=19 Nocodazole cells. ****p<0,0001; Chi-2 test with Yates' correction.

**Figure 8 supplementary 1**

**(A)** Plots representing the z-profile of migration of the blue trajectory from Fig. 8B (same plot as Fig. 8D, right), the associated directionality relative to the centrosome and the speed of migration.

**(B)** Distribution of the velocity of each 953 centriole migration steps of 33 migrating centrioles relative to the centrosome from n=5 cells. Black bars represent the distribution of velocities of steps away from the centrosome and red bars represent the distribution of velocities of steps toward the centrosome.

**(C)** Distribution of the speed of migration plotted from all 953 steps of 33 migrating centrioles versus the 74 extracted steps of apical migration of the 16 centrioles basally located in the cells. See also Fig. 8F.

**(D)** Quantification of Z distribution of centrioles at the end of migration (SAS6 negative) in DMSO, chronic Nocodazole 1 µM and 5 µM treated brain MCCs. Three independent experiments were scored, n=22 DMSO cells analyzed, n=15 Noco 1 µM cells and n=18 Noco 5 µM cells. Error bars represent min to max ± median. ns, non-significant; non-parametric Mann Whitney test.

**(E)** Representative XY distribution of CEN2-GFP+ centrioles at the end of migration (SAS6 negative) in brain MCCs treated with DMSO, Nocodazole 1 µM and 5 µM (48h). Scale bar, 5 µM.

**(F)** Quantification of MBB patch area (in $\mu m^2$) in DMSO and chronic Nocodazole treatment (48h, 1 and 5 µM) in brain MCCs. Three independent experiments were scored, n=22 DMSO cells, n=15 Noco 1 µM cells and n=18 Noco 5 µM cells. Error bars represent min to max ± median. ****p<0,0001; non-parametric Mann Whitney test.

**Figure 9 supplementary 1**

Acceleration of centriole biogenesis during the MCC cell cycle variant leads to a quasi-simultaneous maturation of the centrosomal daughter centriole and newly born procentrioles.

**Figure 9 supplementary 2**

**(A)** mRuby-DEUP1 signal aspect during centriole amplification before and after FRAP. Rows correspond to a centriole amplification stage. Left column: CEN2-GFP; mRuby-DEUP1 profiles before mRuby-DEUP1 bleaching ("pre-bleach"). Middle column: CEN2-GFP; mRuby-DEUP1 profiles right after mRuby-DEUP1 bleaching ("t0 post-bleach"). Right column: CEN2-GFP; mRuby-DEUP1 profiles 10 minutes after mRuby-DEUP1 bleaching ("t+10 min post-bleach"). Scale bar, 2 µm.

**(B)** Quantification of the mRuby-DEUP1 signal intensity overtime ( min) in all the stages of centriole amplification after FRAP.

**(C)** Quantification of the mRuby-DEUP1 signal intensity overtime ( min) in the bleached ROI compared to an unbleached ROI.

### Movie legends

**Video 1** A-stage dynamics in CEN2-GFP (white); mRuby-DEUP1+ (red) cells. A mRuby-DEUP1+ cloud forms around the centrosomal centrioles. Arrowheads point to centrosomal centrioles in the first and last time frames. As the A-stage progresses in time, CEN2-GFP+; mRuby-DEUP1+ procentriole loaded deuterosomes exit the cloud. CEN2-GFP is in white, mRuby-DEUP1 is in red. Left panel is a merge, middle panel is CEN2-GFP with enhanced contrast, right panel is mRuby-DEUP1. Scale bar, 5 µm. dt=50min; displayed at 2 frame/s.

**Video 2** CEN2-GFP and mRuby-DEUP1 dynamics during the cloud stage. CEN2-GFP+ and mRuby-DEUP1+ foci are not always colocalized and some DEUP1+ foci can interact persistently with one parental centriole. A single z-slice of 0.5 µm is showed. dt=1h; CEN2-GFP in white, DEUP1 in red. Arrowhead point to centrosomal centrioles. Scale bar, 1 µm. Displayed at 2 frame/s.

**Video 3** Serial ultra-thin sections (50nm) of the cell represented in Fig. 2 supplementary 4. White arrowheads points to centrosomal mother centriole. Green arrowhead points to centrosomal daughter centriole. Scale bar, 0,5µm.

**Video 4** mRuby-DEUP1+/CEN2-GFP negative foci oscillatory dynamics during early A-stage. mRuby-DEUP1+/CEN2-GFP negative foci can oscillate back and forth between the cytoplasm and the centrosomal centrioles (arrowheads). Some foci can stay still for a long period of time before moving again. A single z-slice of 0.5 µm is filmed. Black cross indicates an artefactual CEN2-GFP aggregate; CEN2-GFP in white, mRuby-DEUP1 in red; dt=5s; scale bar, 1µm. Displayed at 3 frame/s.

**Video 5** mRuby-DEUP1 negative/CEN2-GFP+ foci oscillatory dynamics during early A-stage. mRuby-DEUP1 negative/CEN2-GFP+ foci can oscillate back and forth between the cytoplasm and the centrosomal centrioles (arrowheads; mother centriole in white and daughter centriole in green). Some foci can stay still for a long period of time before moving again. A single z-slice of 0.5 µm is filmed. CEN2-GFP in white, mRuby-DEUP1 in red; dt=5s; scale bar, 1 µm. Displayed at 3 frame/s.

**Video 6** Deuterosome behavior during A-stage. Procentriole loaded deuterosomes remain in the centrosome vicinity (arrowheads) and describe back and forth movements. A single z-slice of 0.5 µm is filmed. CEN2-GFP in white, mRuby-DEUP1 in red. Note that a deuterosome stays connected to one centrosomal centriole. dt=5s; scale bar, 1 µm. Displayed at 3 frame/s.

**Video 7** mRuby-DEUP1+ foci (arrows) intermittent oscillatory interactions with one of the two centrosomal centriole (arrowheads) in CEN2-GFP+ cells. CEN2-GFP in white, mRuby-DEUP1 in red. A single z-slice of 0.5 µm is filmed. dt=5s. Scale bar, 5 µm. Displayed at 3 frame/s.

**Video 8** Block of the mRuby-DEUP1+/CEN2-GFP+ foci oscillatory dynamics during early A-stage after acute nocodazole treatment (1h, 10µM). A single z-slice of 0.5 µm is filmed. CEN2-GFP in white, mRuby-DEUP1 in red. Arrowheads point to centrosomal centrioles. dt=5s; scale bar, 1 µm. Displayed at 3 frame/s.

**Video 9** Block of the deuterosome oscillatory dynamics during A-stage after acute nocodazole treatment (1h, 10µM). A single z-slice of 0.5 µm is filmed. CEN2-GFP in white, mRuby-DEUP1 in red. Arrowheads point to centrosomal centrioles. dt=5s; scale bar, 1 µm. Displayed at 3 frame/s.

**Video 10** Block of the mRuby-DEUP1+/CEN2-GFP+ foci oscillatory dynamics during early A-stage after acute dynapyrazole treatment (1h, 7.5µM). A single z-slice of 0.5 µm is filmed. CEN2-GFP in white, mRuby-DEUP1 in red. X indicates CEN2-GFP aggregate. Arrowhead points to one centrosomal centriole (the other centrosomal centriole is out of focus). dt=5s; scale bar, 1 µm. Displayed at 3 frame/s.

**Video 11** mRuby-DEUP1+/CEN2-GFP+ deuterosome oscillatory dynamics during A-stage before (left) and after (right) acute dynapyrazole treatment (1h, 7.5µM). A single z-slice of 0.5 µm is filmed. CEN2-GFP in white, mRuby-DEUP1 in red. Arrowheads point to typical oscillatory dynamics. Empty arrowheads indicate centrosomal centrioles. X indicates CEN2-GFP aggregate. dt=5s; scale bar, 1 µm. Displayed at 1 frame/s.

**Video 12** CEN2-GFP (left); mRuby-DEUP1 (right) profiles in control A-stage cell. t=0h00 corresponds to the first acquisition when cells were still untreated. DMSO is added just after the first acquisition. At the onset of the movie, a mRuby-DEUP1+;CEN2-GFP+ cloud (outlined in white in this example and defined using high contrast) is already formed around the centrosomal centrioles (arrowheads); procentrioles already start associating to DEUP1+ focis inside the cloud. As the cell progresses in A-stage, the early deuterosomes loaded with procentrioles accumulate in the cloud and start migrating outside the cloud. Quantitative measurements (surface of the pericentrosomal cloud and surface of the area covered with deuterosomes) were done on multiple cells and plotted in Fig. 4E and Fig. 4 Supplementary 2A. CEN2-GFP is on the left and mRuby-DEUP1 is on the right. dt=30min; Scale bar, 5µm. Displayed at 1 frame/s.

**Video 13** CEN2-GFP (left); mRuby-DEUP1 (right) profile in A-stage cell under acute Nocodazole (10µM) treatment. t=0h00 corresponds to the first acquisition when cells were still untreated. Nocodazole is added just after the first acquisition. At the onset of the movie, mRuby-DEUP1+/CEN2-GFP+ cloud (outlined in white in this example and defined using high contrast) is already formed around the centrosomal centrioles (arrowheads); procentrioles already associated to DEUP1+ focis are inside the cloud and in the cytoplasm. After Nocodazole addition, the mRuby-DEUP1+/CEN2-GFP+ cloud cloud is shrinking in size for a few hours before disappearing. The already formed procentriole loaded deuterosomes disperse in 1h after Nocodazole addition and new ones are seen appearing at random spots in the cytoplasm, covering almost all the cell's surface (t+10h30, end of the movie). Quantitative measurements (surface of the pericentrosomal cloud and surface of the area covered with deuterosomes) were done on multiple cells and plotted in Fig. 4E and Fig. 4 Supplementary 2A. CEN2-GFP is on the left and mRuby-DEUP1 is on the right. dt=30min; Scale bar, 5µm. Displayed at 1 frame/s.

**Video 14** CEN2-GFP (left); mRuby-DEUP1 (right) profile in A-stage cell under acute Dynapyrazole (3µM) treatment. Dynapyrazole is added just after the first acquisition. At the onset of the movie, a mRuby-DEUP1+/CEN2-GFP+ cloud (outlined in white in this example) is already formed around the centrosomal centrioles (arrowheads); procentrioles already associated to DEUP1+ focis are inside the cloud. After Dynapyrazole addition, the CEN2-GFP+; mRuby-DEUP1+ cloud is shrinking in size for a few hours before disappearing. Quantitative measurements (surface of the pericentrosomal cloud and surface of the area covered with deuterosomes) were done on multiple cells and plotted in Fig. 4E and Fig. 4 Supplementary 2A. CEN2-GFP is on the left and mRuby-DEUP1 is on the right. dt=40min; Scale bar, 5µm.

**Video 15** CEN2-GFP dynamics in A-stage cells treated with DMSO (left) and Dynapyrazole (right, 3μM). DMSO or Dynapyrazole were added after the first time point. White arrowheads point to procentrioles. Red arrowheads point to centrosomal centrioles. dt=40 min, scale bar= 1μm.

**Video 16** CEN2-GFP dynamics during G-stage where centrioles -organized as flower-like structures around non-fluorescent deuterosomes- are seen migrating along the nucleus, identified by contrasting CEN2-GFP. dt=5min. Scale bar, 5 μm

**Video 17** CEN2-GFP dynamics in G and D-stage. Apical view of a group of procentriole loaded deuterosome (empty « flowers ») in G-stage (t0), covering the surface of the nucleus and distributing around the nuclear envelope before disengaging. After disengagement of the centrioles, they are seen reorganizing around the nucleus (t16). The nucleus is visible thanks to the absence of CEN2-GFP signal. dt=10min. Scale bar, 5 μm.

**Video 18** Super-resolution imaging of a CEN2-GFP (white) D-stage cell immunostained with MAB414 (nuclear pores, red) and counterstained with Hoescht. Scale bar=1μm.

**Video 19** CEN2-GFP; mRuby-DEUP1 dynamics during D-stage. In early D-stage (t0), most centrioles are organized in rosettes around deuterosomes. Some are already isolated and appear at singlets on deuterosomes. By t4, centrioles are no longer organized in rosettes; they appear individualized when looking at the CEN2-GFP channel. Some deuterosomes remain but are less spherical and connected with a single centriole. During the rest of the movie, mRuby-DEUP1+ signal oscillates between a diffuse and more aggregated state, mostly connected with centrioles. The mRuby-DEUP1+ diffuse signal sometimes forms clouds (arrowheads at t6, t9, t17, t21). This is correlated with the reaggregation of centrioles. CEN2-GFP is in white and mRuby-DEUP1 is in red. dt=10min; Movie duration=4h10; scale bar, 5 μm. Displayed at 1 frame/s.

**Video 20** CEN2-GFP; mRuby-DEUP1 movie of a single z-plane of 0.7μm of a D-stage cell showing deuterosome which seemingly split into smaller deuterosomes migrating away from each others along the nuclear envelope. The inset shows a magnified view. CEN2-GFP is in white and mRuby-DEUP1 is in red. Associated with Fig. 6E. dt=5s; scale bar, 1μm. Displayed at 2 frame/s.

**Video 21** CEN2-GFP; mRuby-DEUP1 movie of a single z-plane of 0.7μm of a D-stage cell showing deuterosome dissolution correlated with CEN2-GFP rosette dismantlement. The inset shows a magnified view. CEN2-GFP is in white and mRuby-DEUP1 is in red. Associated with Fig. 6 Supplementary 1D. dt=40min; scale bar, 5µm. Displayed at 2 frame/s.

**Video 22** CEN2-GFP; mRuby-DEUP1 movie a D-stage cell showing deuterosome migration along the nuclear membrane, change in deuterosome shape and mixing/unmixing with another deuterosome. The inset shows a magnified view. CEN2-GFP is in white and mRuby-DEUP1 is in red. Associated with Fig. 6G. dt=40min; scale bar, 5µm. Displayed at 2 frame/s.

**Video 23** Disengaged centriole associated to a deuterosomal subunit migrating and moving back and forth on the nuclear envelope. The inset shows a magnified view of this disengaged centriole. Note the perinuclear position and movement of other centrioles, and the distortion of mRuby-DEUP1 signal of disengaging deuterosome(s) on the right upper part of the nucleus. The shape of the nucleus is seen thanks to the absence of CEN2-GFP signal. CEN2-GFP is in white and mRuby-DEUP1 is in red. dt=5s; scale bar, 5 µm; single z-section of 0.5µm. Displayed at 2 frame/s.

**Video 24** Tyrosinated microtubules (YL1/2, white) and DEUP1 (red) immuno-reactivity profiles in brain CEN2-GFP (green) MCC progenitors during D-stage. This movie shows 0.25µm z-sections of the whole cell. Note that all centrioles are next to a MT fiber. Scale bar, 5 µm.

**Video 25** CEN2-GFP dynamics in D-stage of brain MCC progenitors under acute DMSO (left), nocodazole (10µM, middle) or dynapyrazole (3µM, right) treatments. DMSO and drugs are added just after the first time point. Compared to DMSO, the centrioles organized as flowers around deuterosomes (not visible here) of the cells treated with nocodazole (10µM) or dynapyrazole (3µM) take more time, or even fail to dismantle. dt=30 min. Scale bar, 5 µm. Displayed at 3 frame/s.

**Video 26** Movie showing the final migration of individual centrioles to the apical membrane. **Upper pannel:** apical view of manually tracked migrating centrioles trajectories. The circle indicates the centrosomal centriole taken as the origin. **Bottom panel:** corresponding profile view of the top panel. At t0, centrioles have just disengaged and are isotropically organized around the nucleus. Centrioles progressively migrate up. Each color represents an individual centriole trajectory track. dt=5min. Scale bar, 5 µm.

**Video 27** Profile view of centriole apical migration after D-stage in CEN2-GFP+ brain MCC peogenitors treated with DMSO and acute Nocodazole 10µM. At t0, cells display late G-stage/early D-stage procentrioles and are not treated with any drug. DMSO and Nocodazole 10µM were added right after the first timepoint acquisition. In DMSO, 2 centriole patches are visible above and below the nucleus (t+0h). The patch below the nucleus progressively joins the apical one to become one apical patch and stabilizes. In 10µM Nocodazole, centrioles migrate up and down and fail to form a stable apical patch. dt=5min; scale bar, 1 µm.

**Video 28** CEN2-GFP; mRuby-DEUP1 movie of a single z-plane (0.7µm) of a G-stage cell showing deuterosome bumping into each other and staying close together without fusing. dt=5s; scale bar, 1µm. Displayed at 2 frame/s.

#### Figure 1 supplementary 1

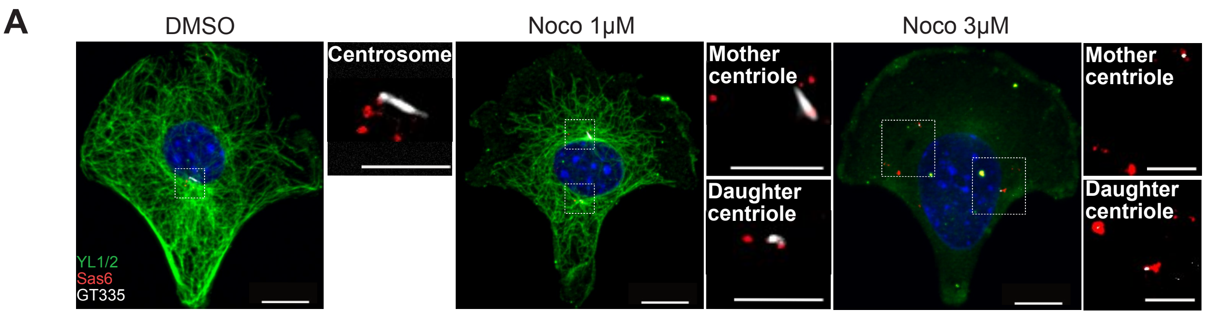

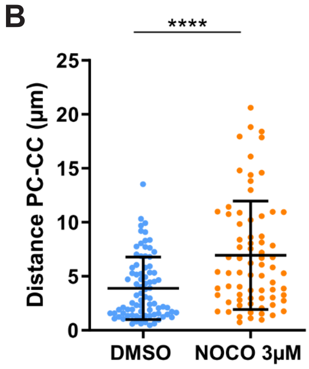

#### Figure 2 supplementary 1

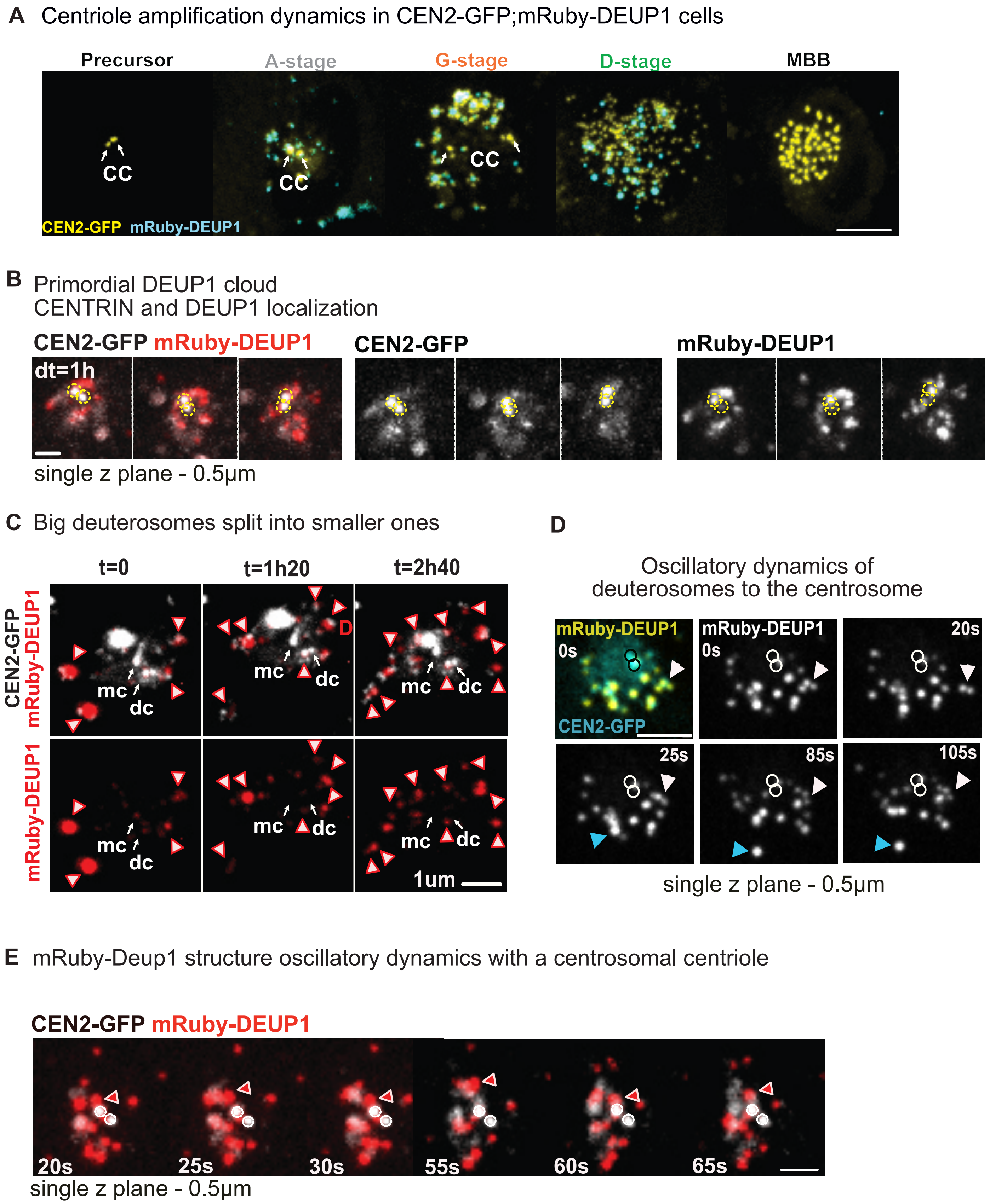

Figure 2 supplementary 2

#### mRuby-DEUP1 primordial cloud - Correlative light and electron microscopy (CLEM)

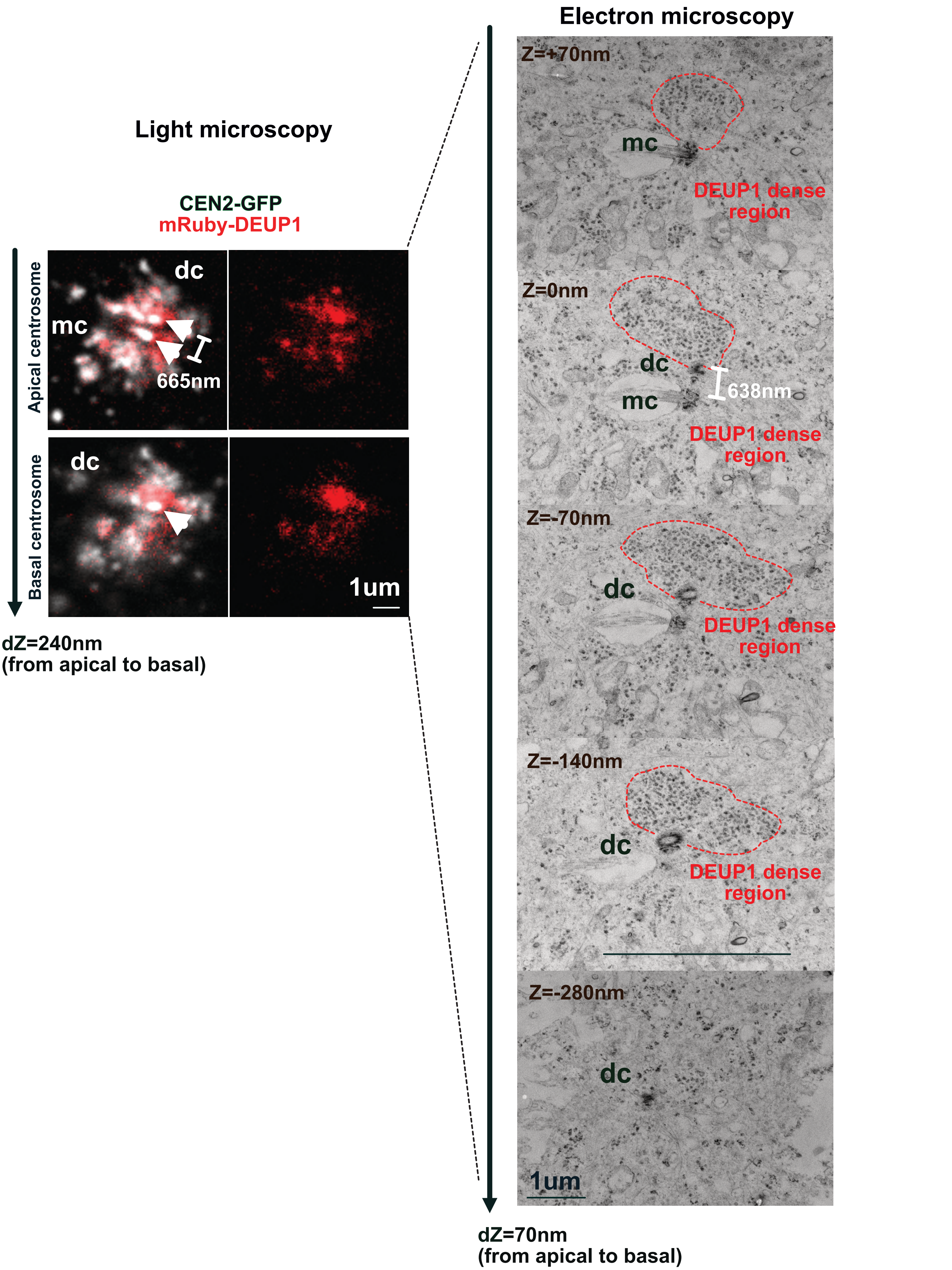

Figure 2 supplementary 3

#### mRuby-DEUP1 primordial cloud - Correlative light and electron microscopy (CLEM)

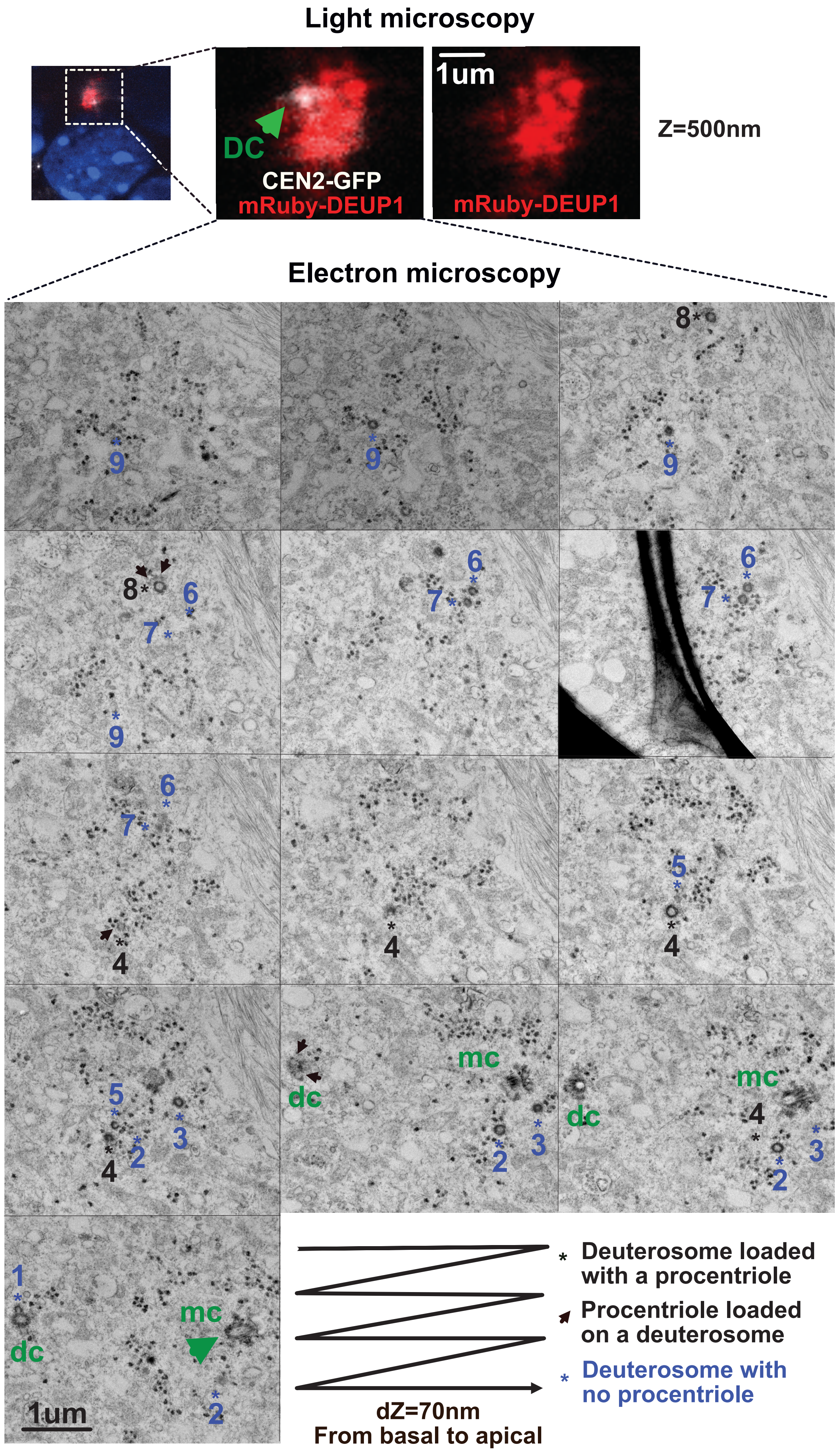

Figure 2 supplementary 4

#### mRuby-Deup1 primordial cloud - Correlative light and electron microscopy (CLEM)

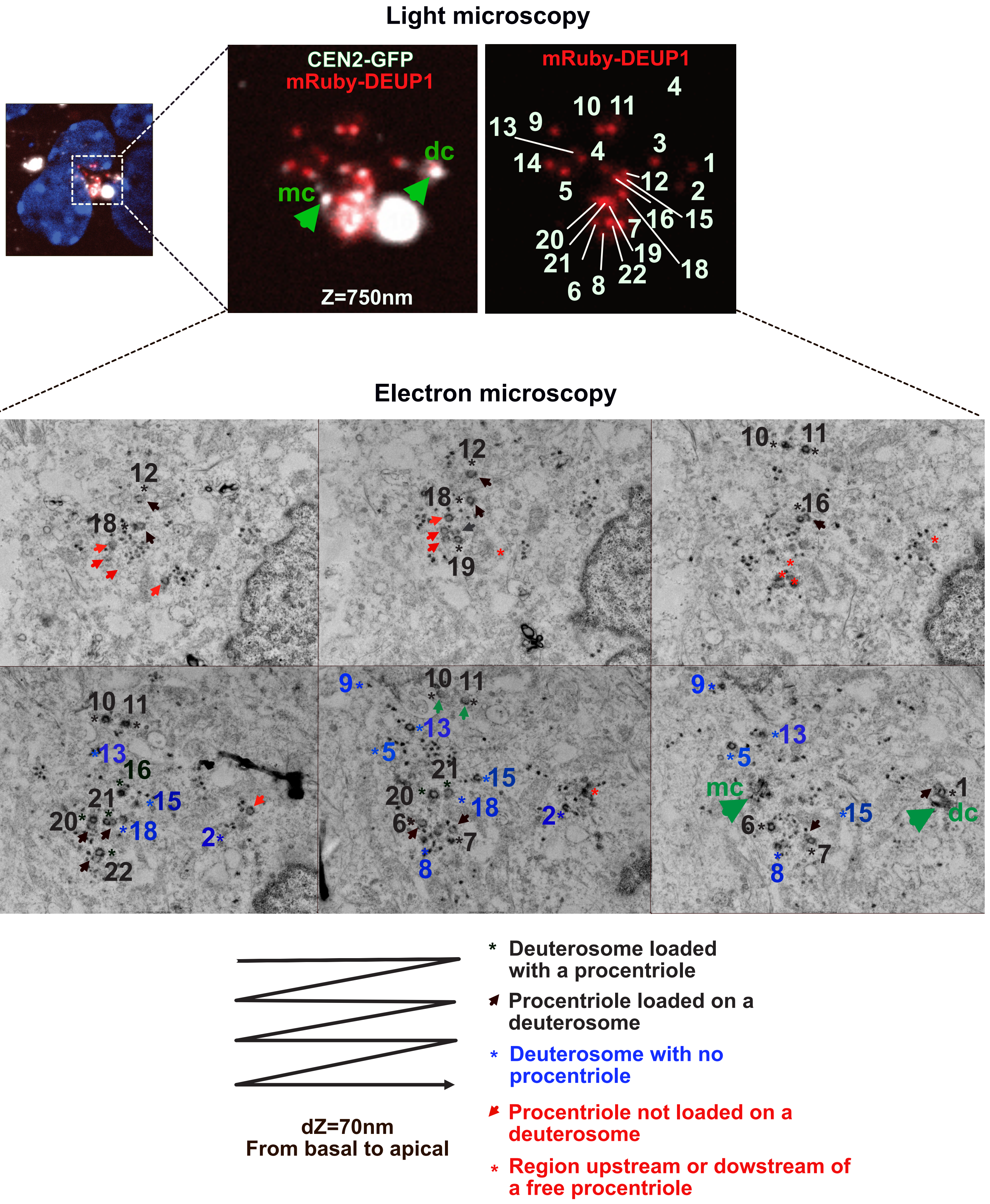

Figure 2 supplementary 5

#### Early pericentrosomal cloud - Correlative light and electron microscopy (CLEM)

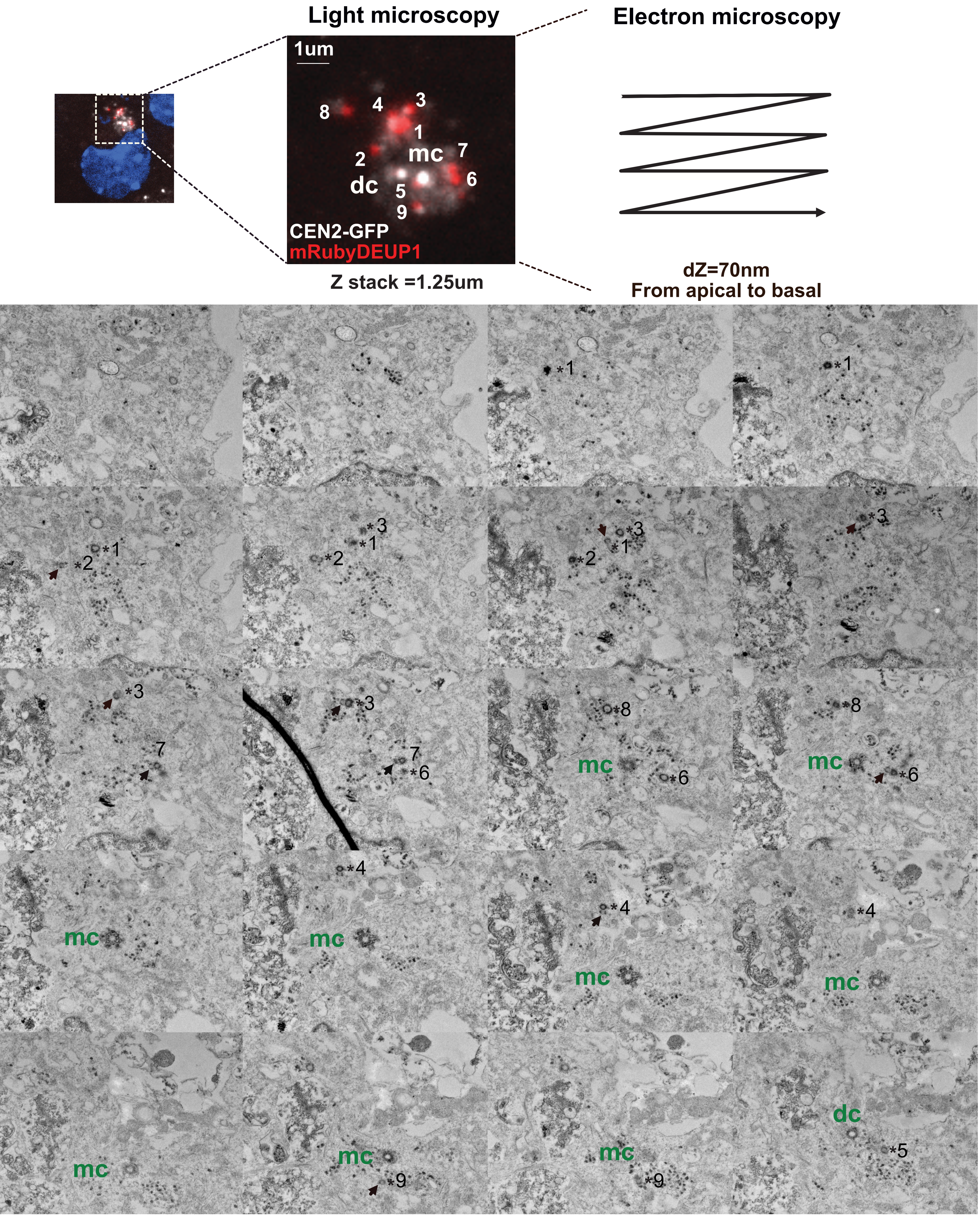

* Deuterosome loaded with a procentriole

Procentriole loaded on a deuterosome

Figure 2 supplementary 6

#### mRuby-Deup1 primordial cloud - Correlative light and electron microscopy (CLEM)

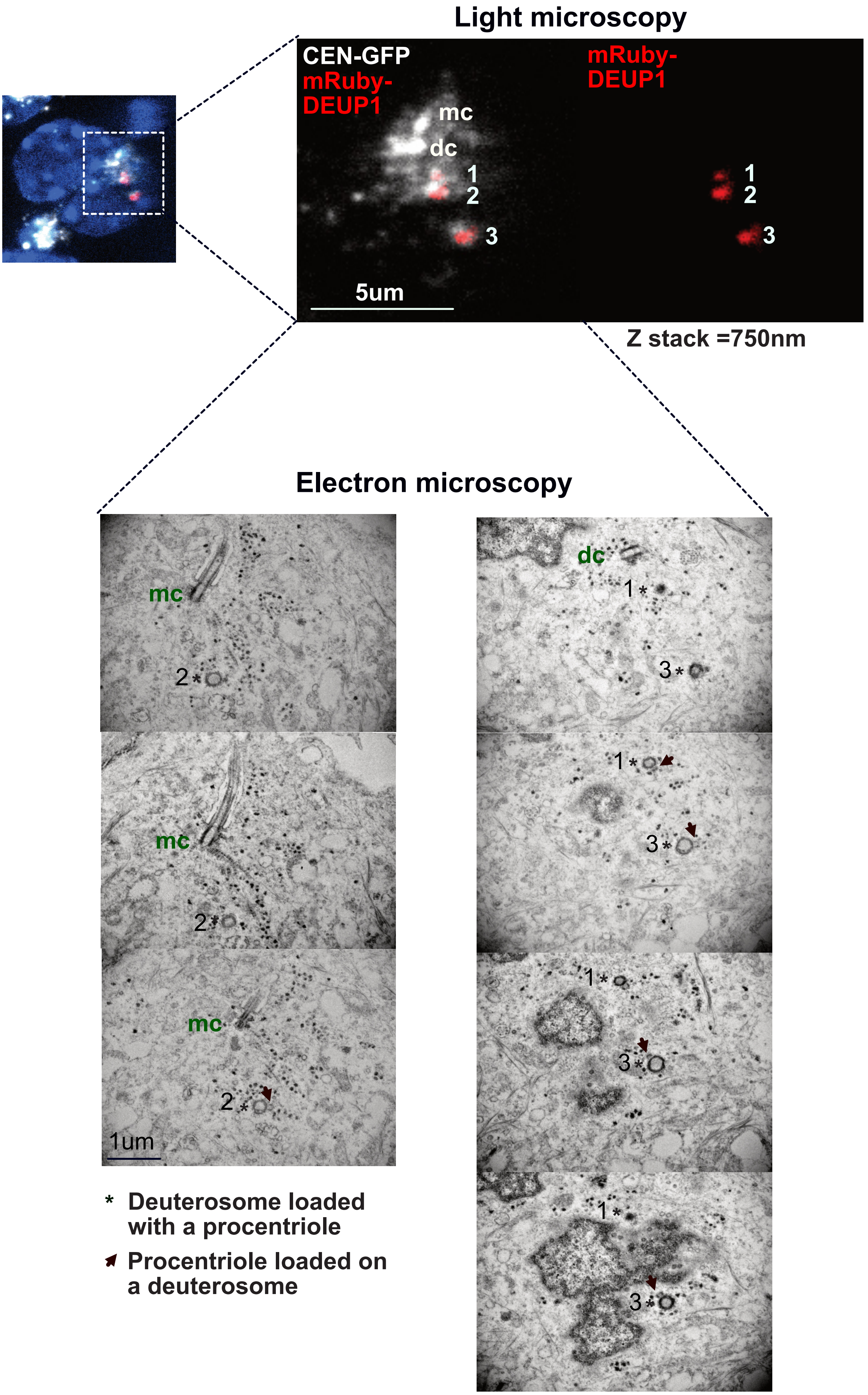

* Deuterosome loaded with a procentriole

➶ Procentriole loaded on a deuterosome

dZ=70nm
From apical to basal

**Figure 3 supplementary 1**

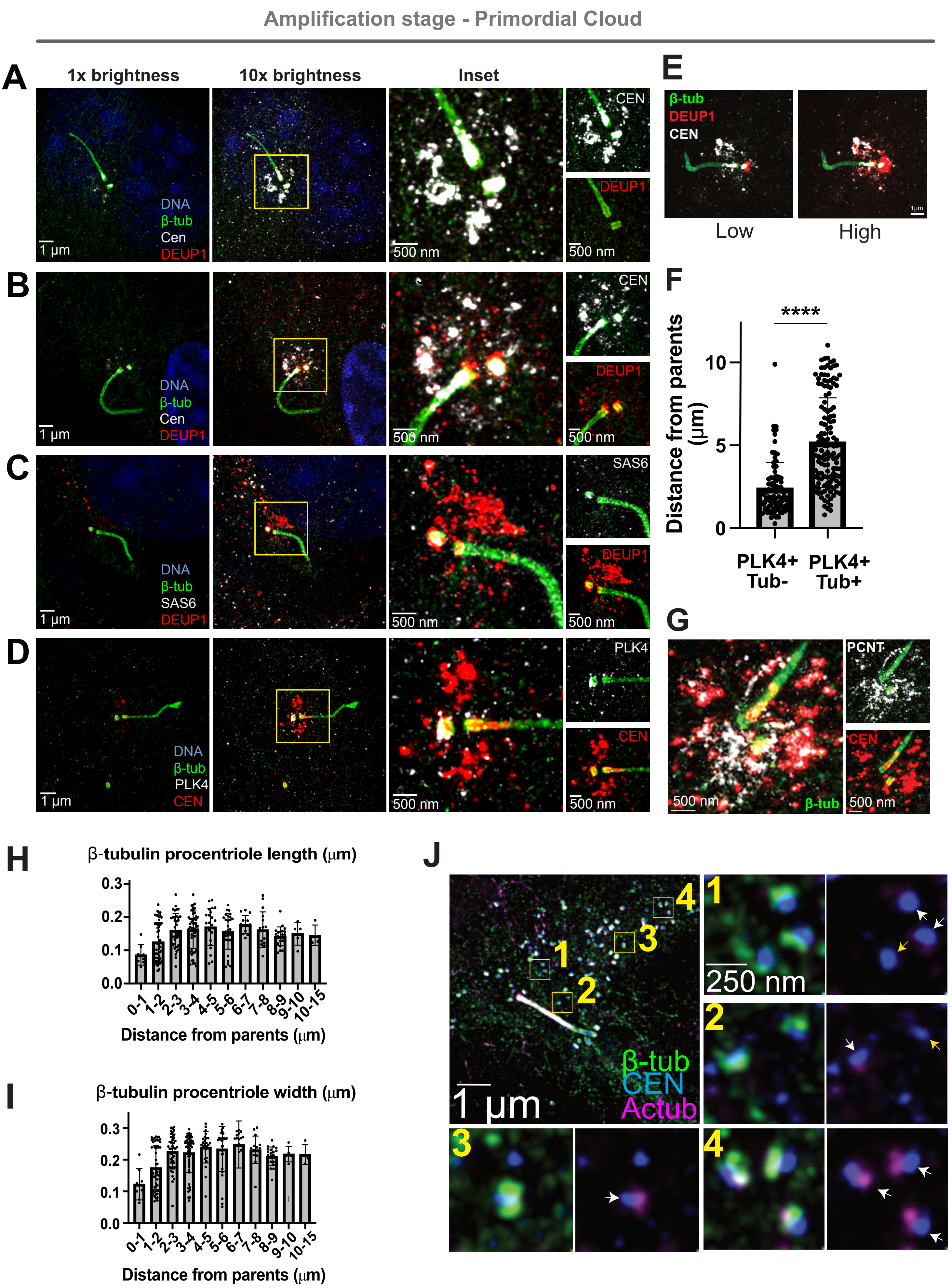

### Figure 3 Supplementary 2

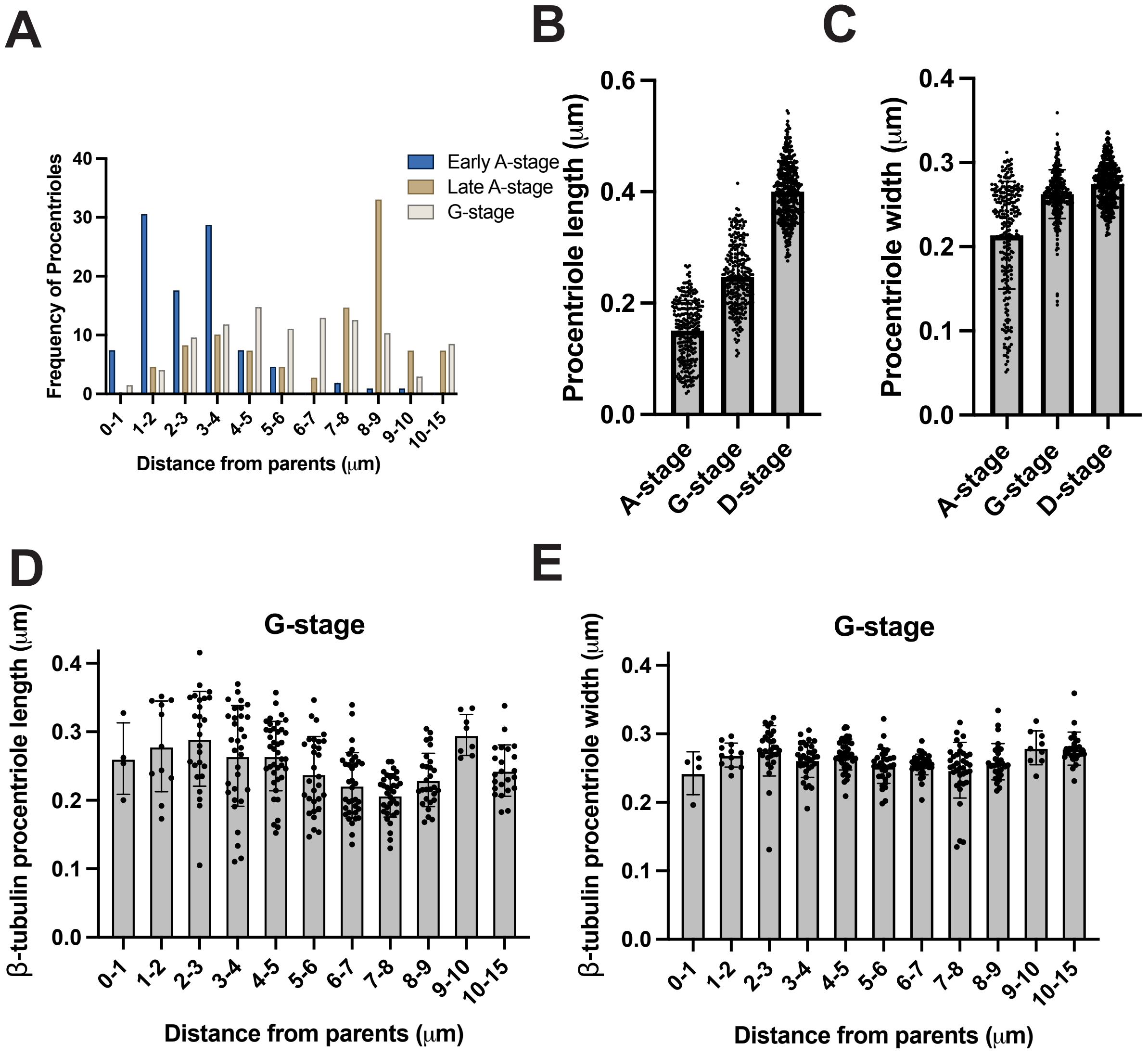

Figure 4 supplementary 1

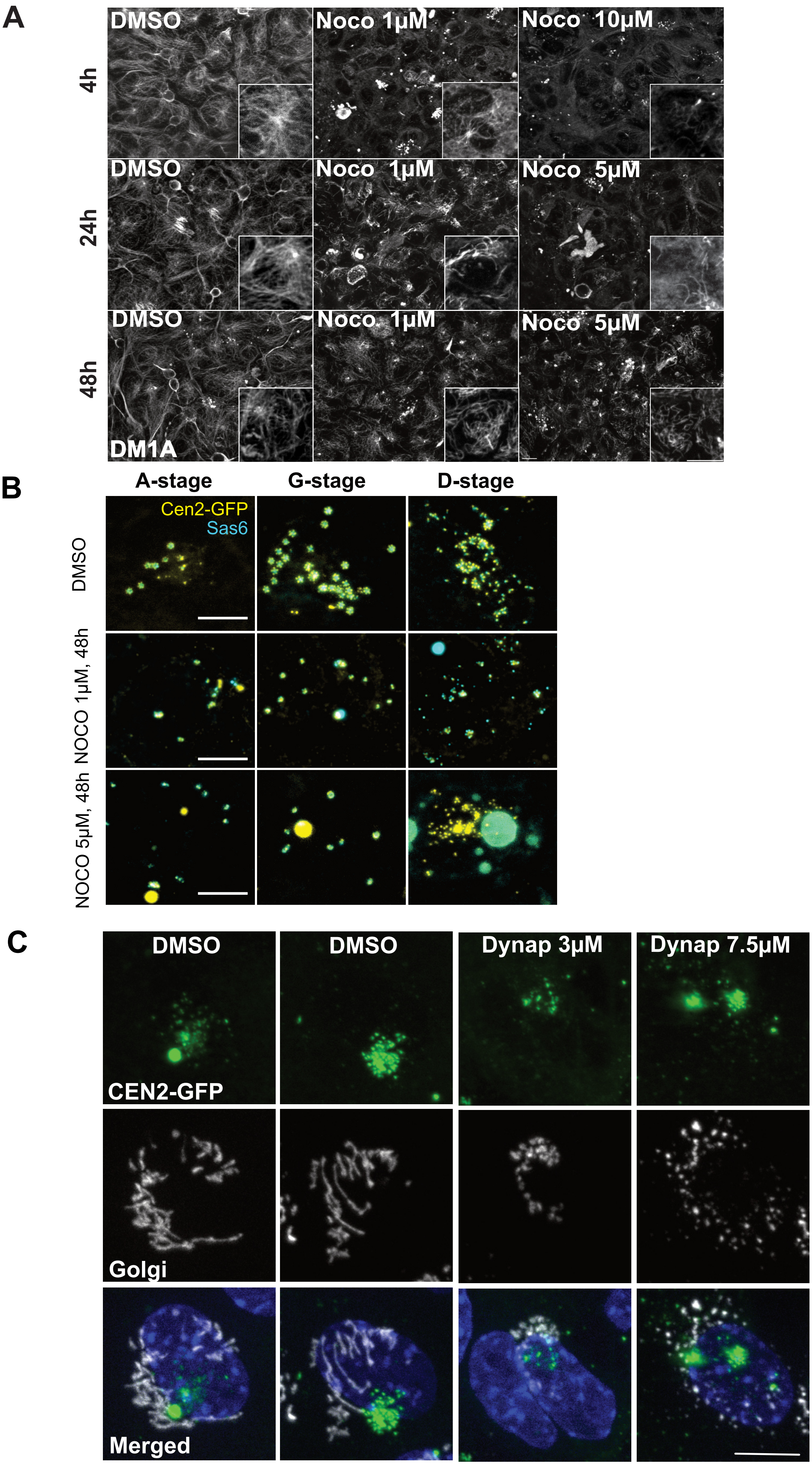

Figure 4 supplementary 2

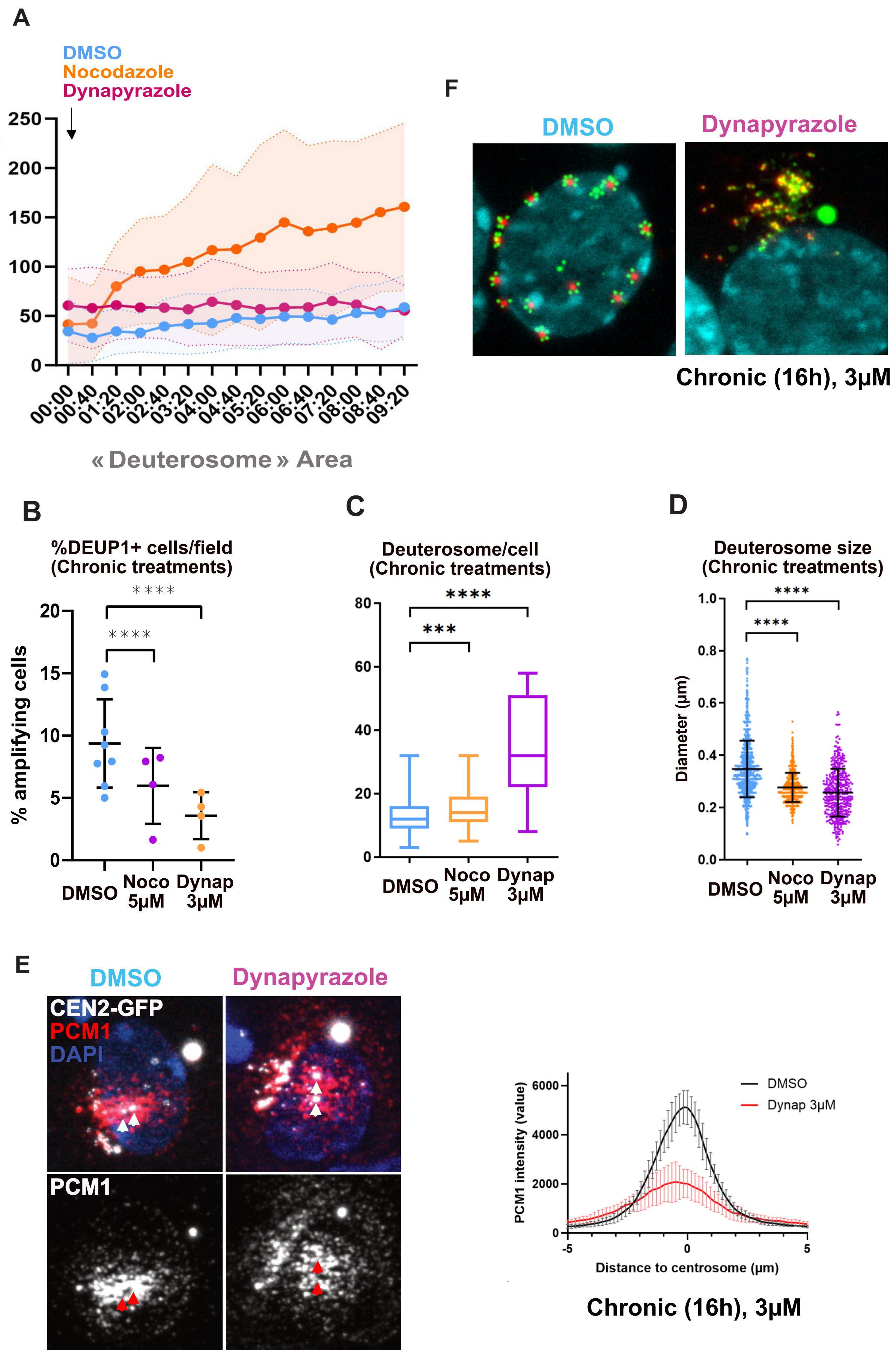

Figure 4 supplementary 3

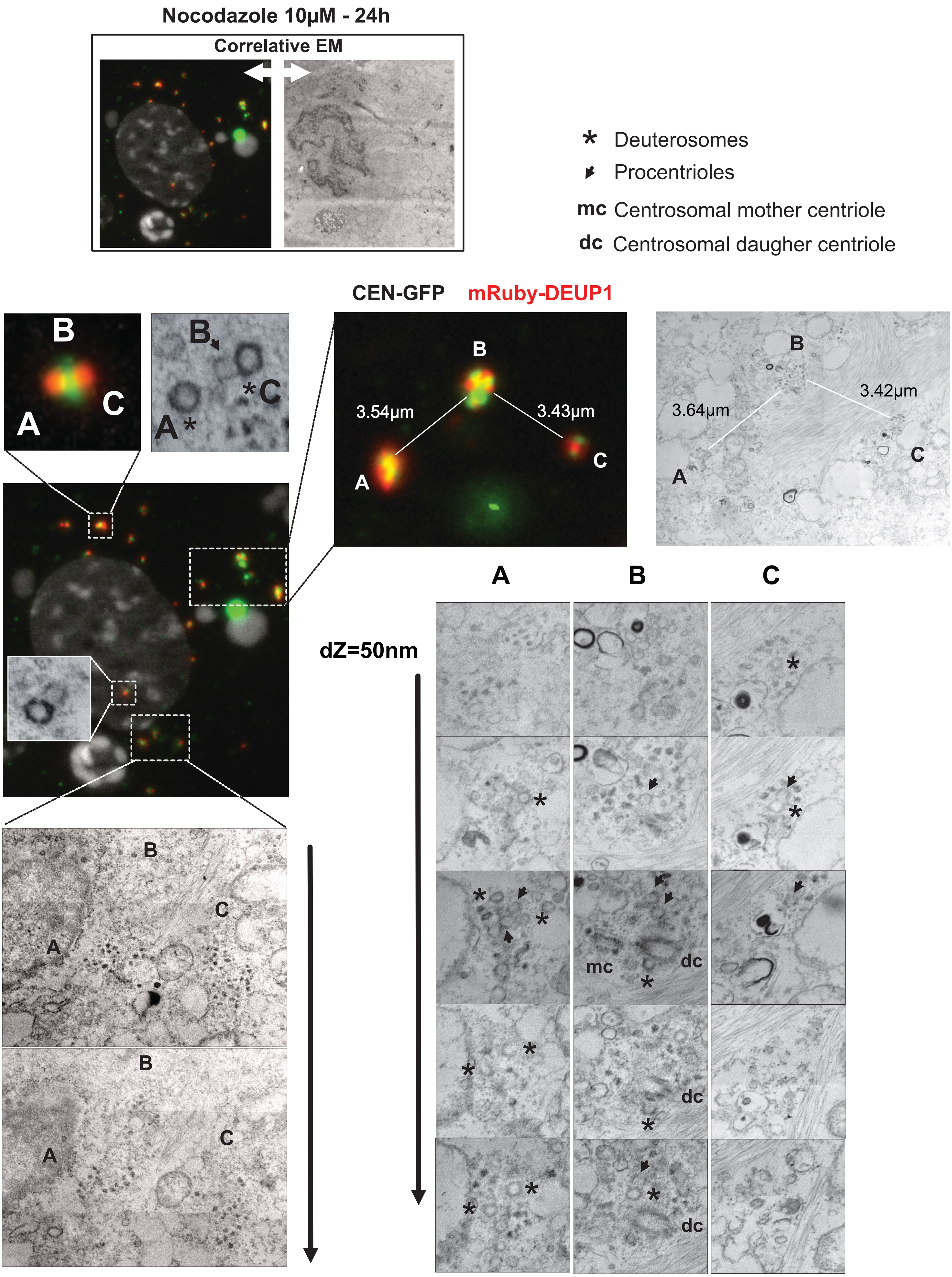

Figure 4 supplementary 4

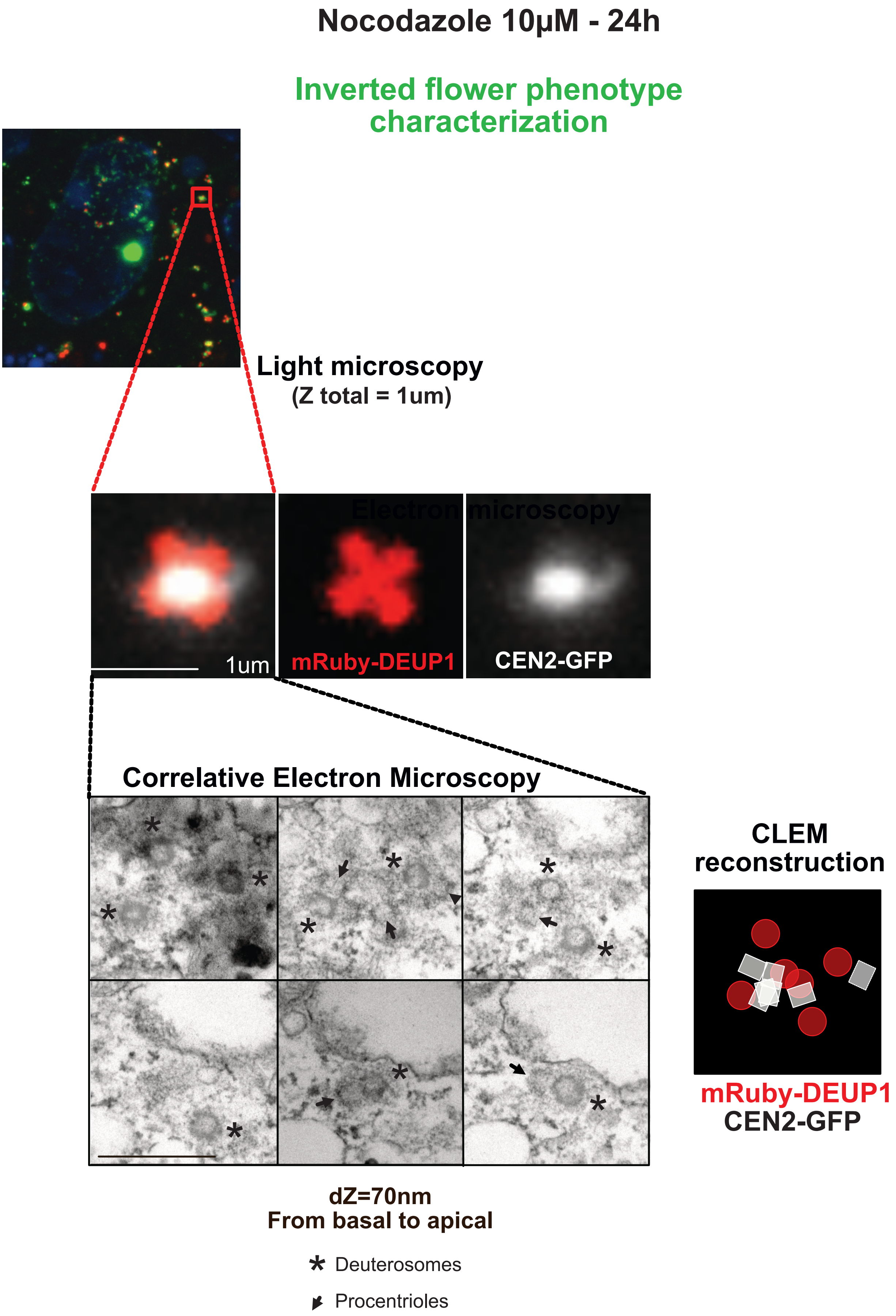

#### Figure 5 supplementary 1

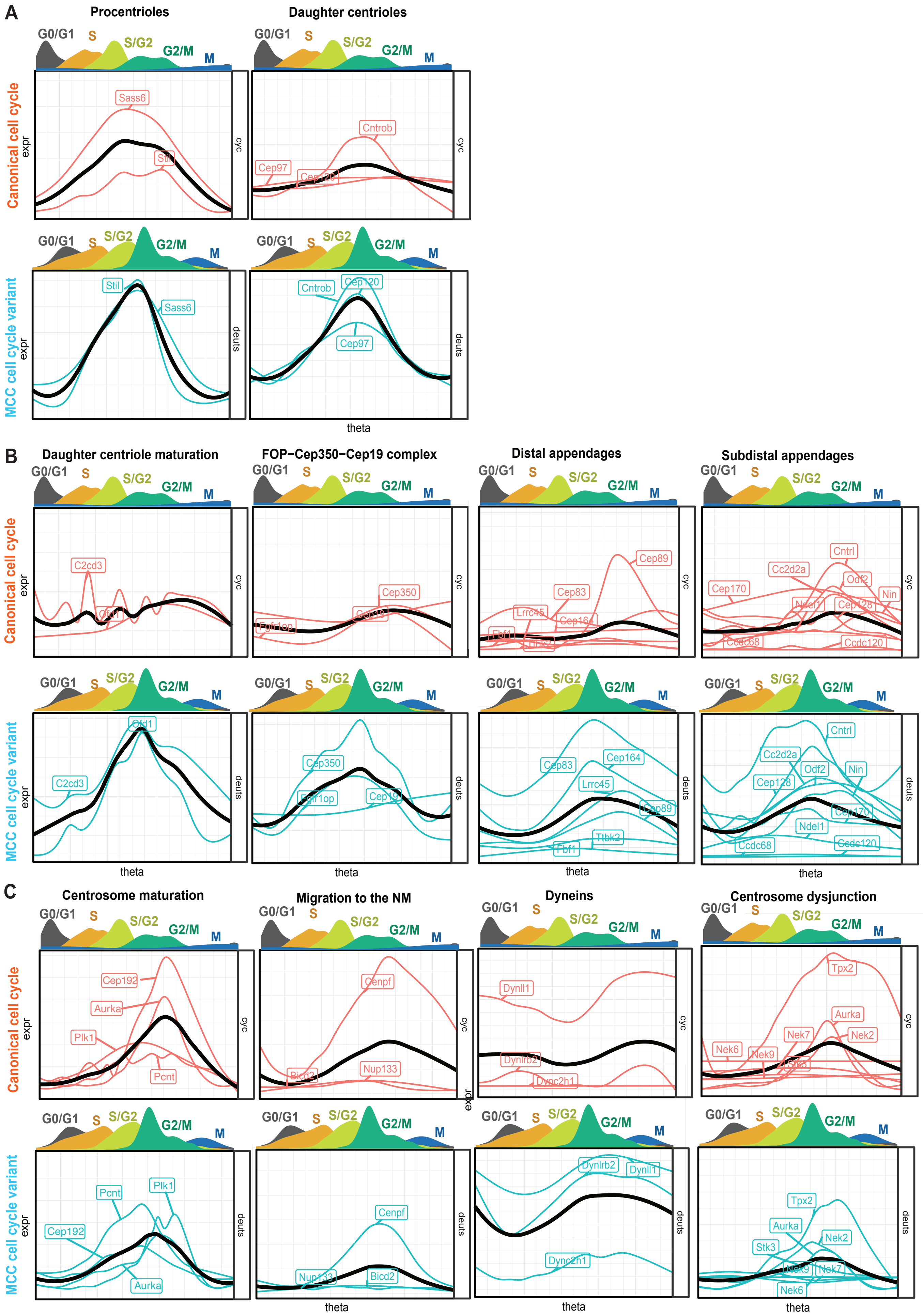

### Figure 5 supplementary 2

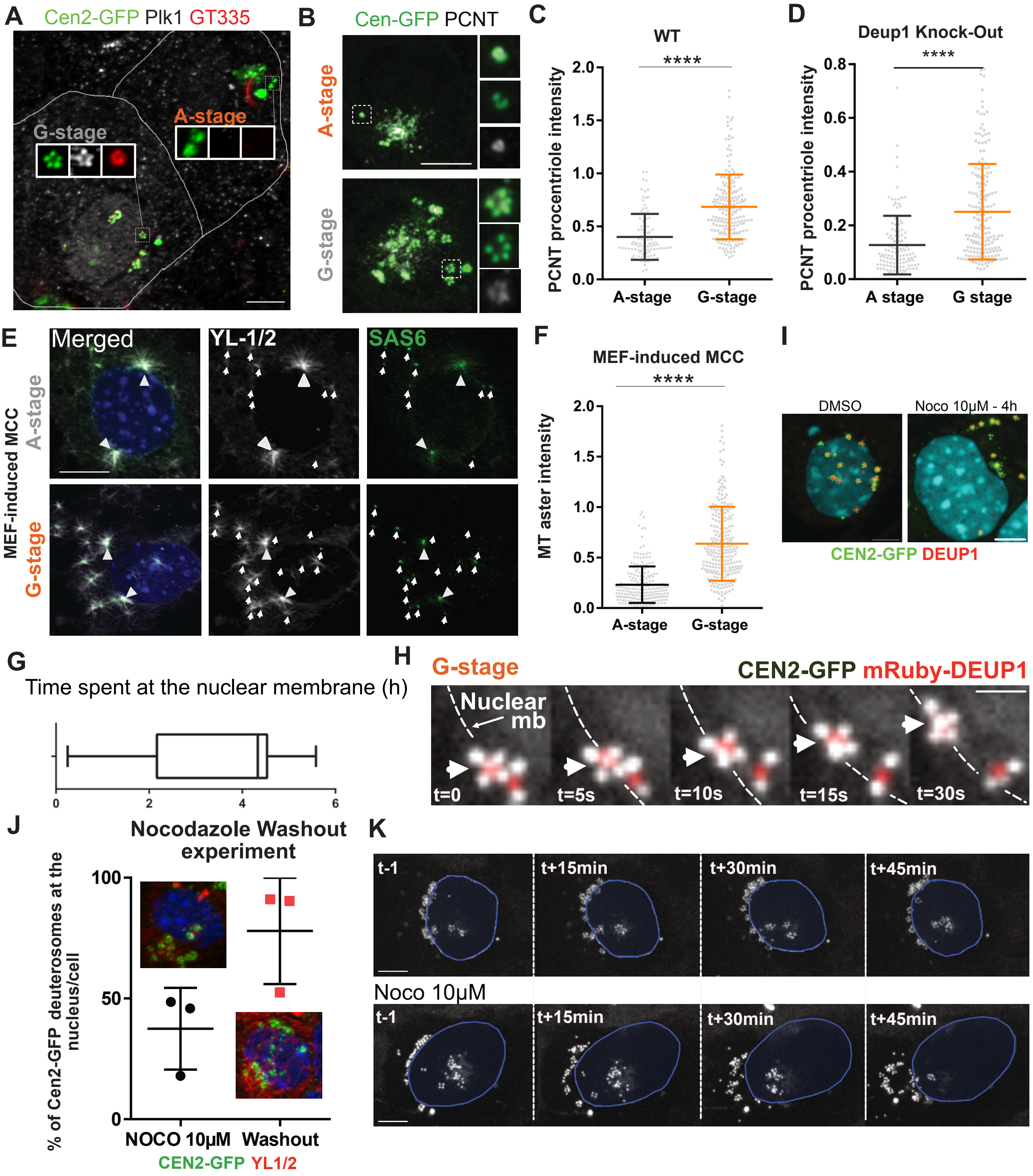

#### Figure 6 supplementary 1

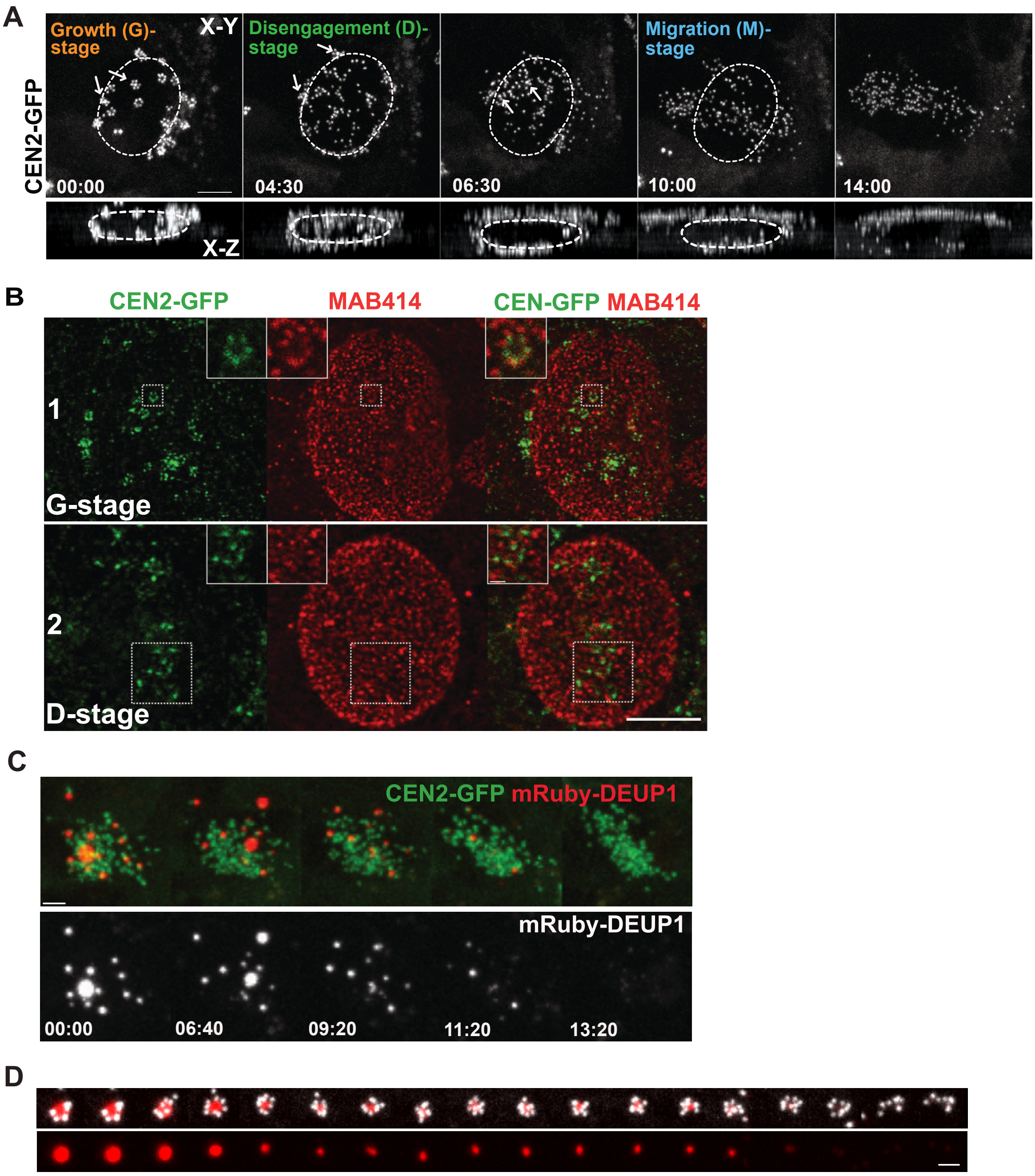

CEN-GFP mRuby-DEUP1 - dt = 40 min

#### Figure 7 supplementary 1

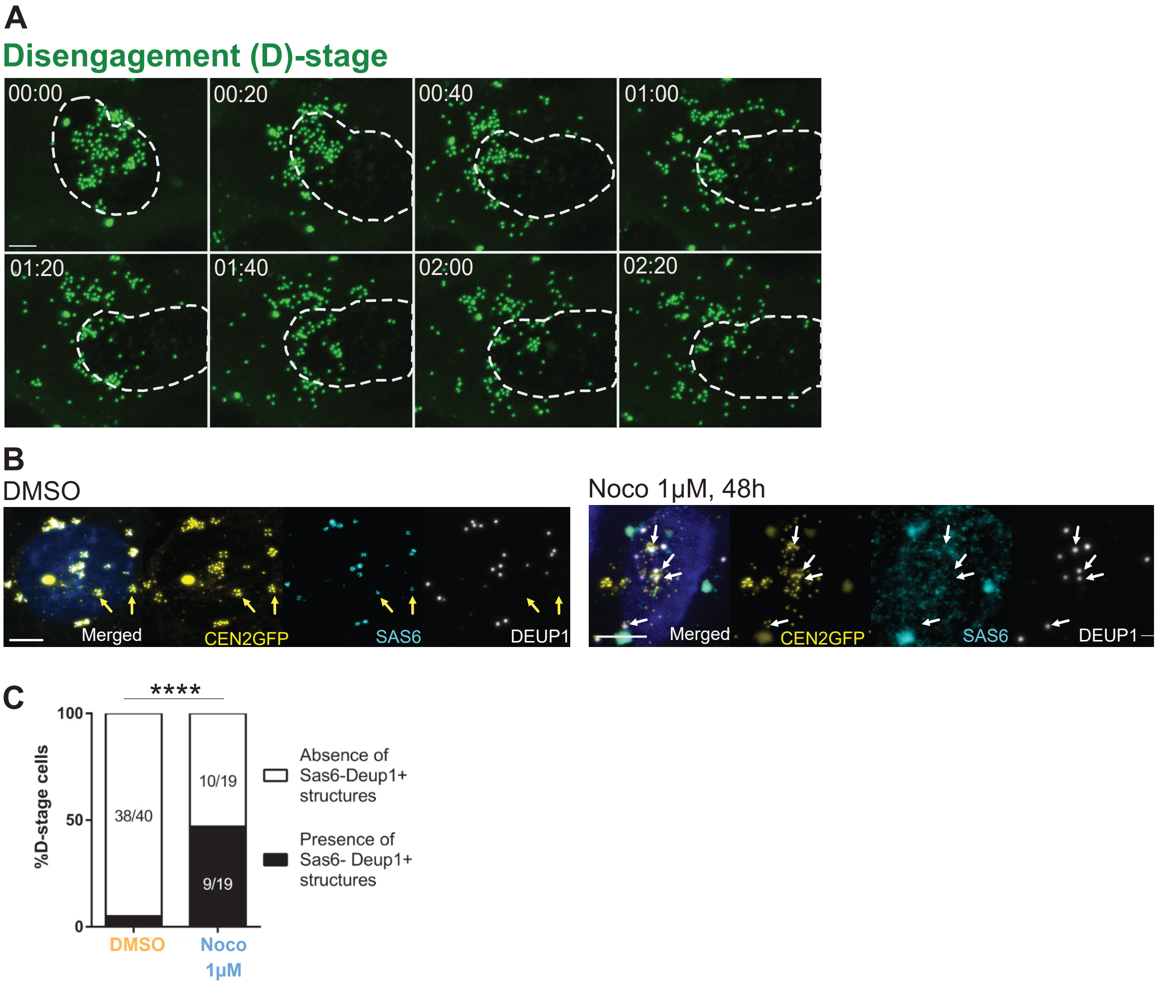

### Figure 8 supplementary 1

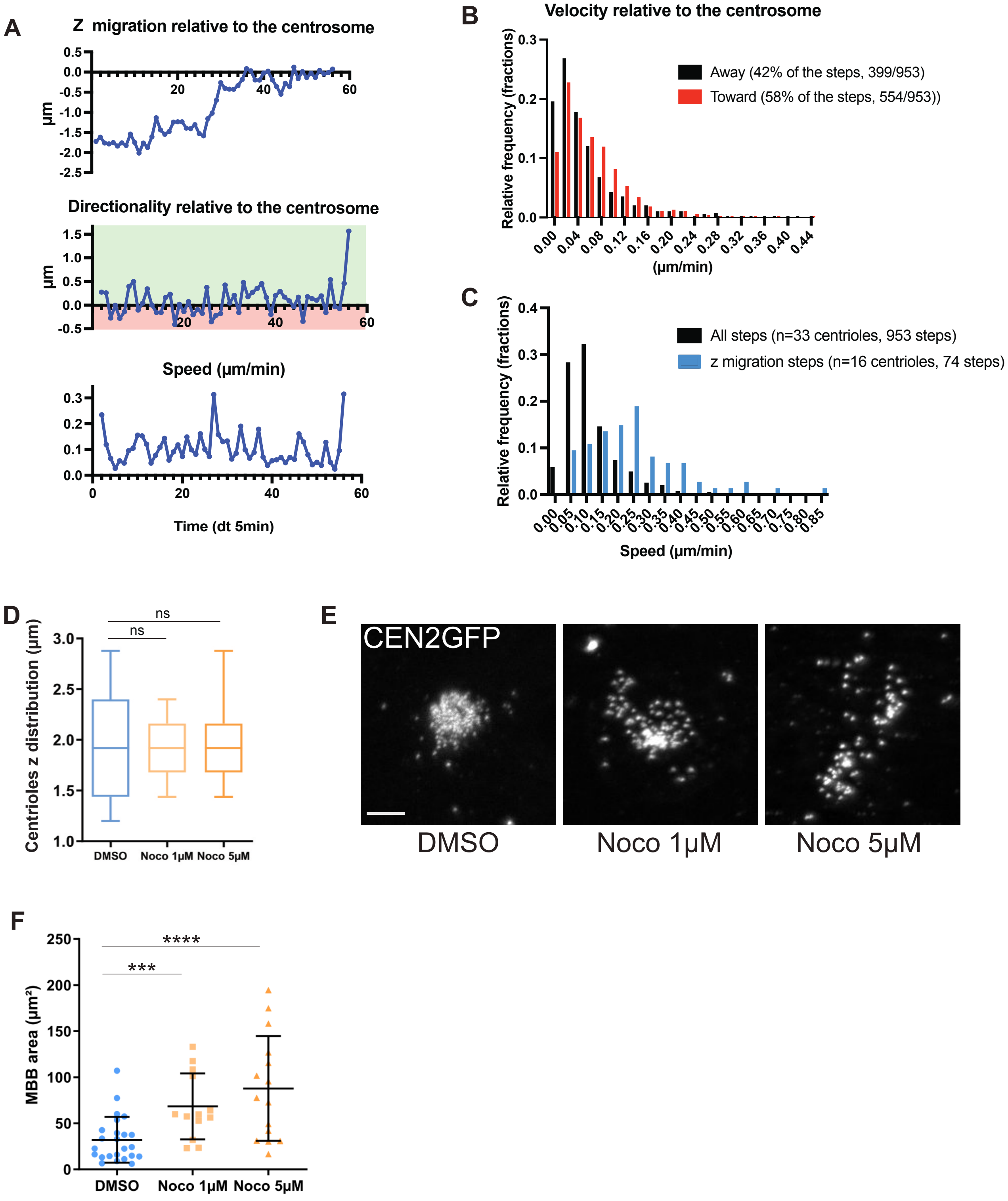

### Figure 8 supplementary 2

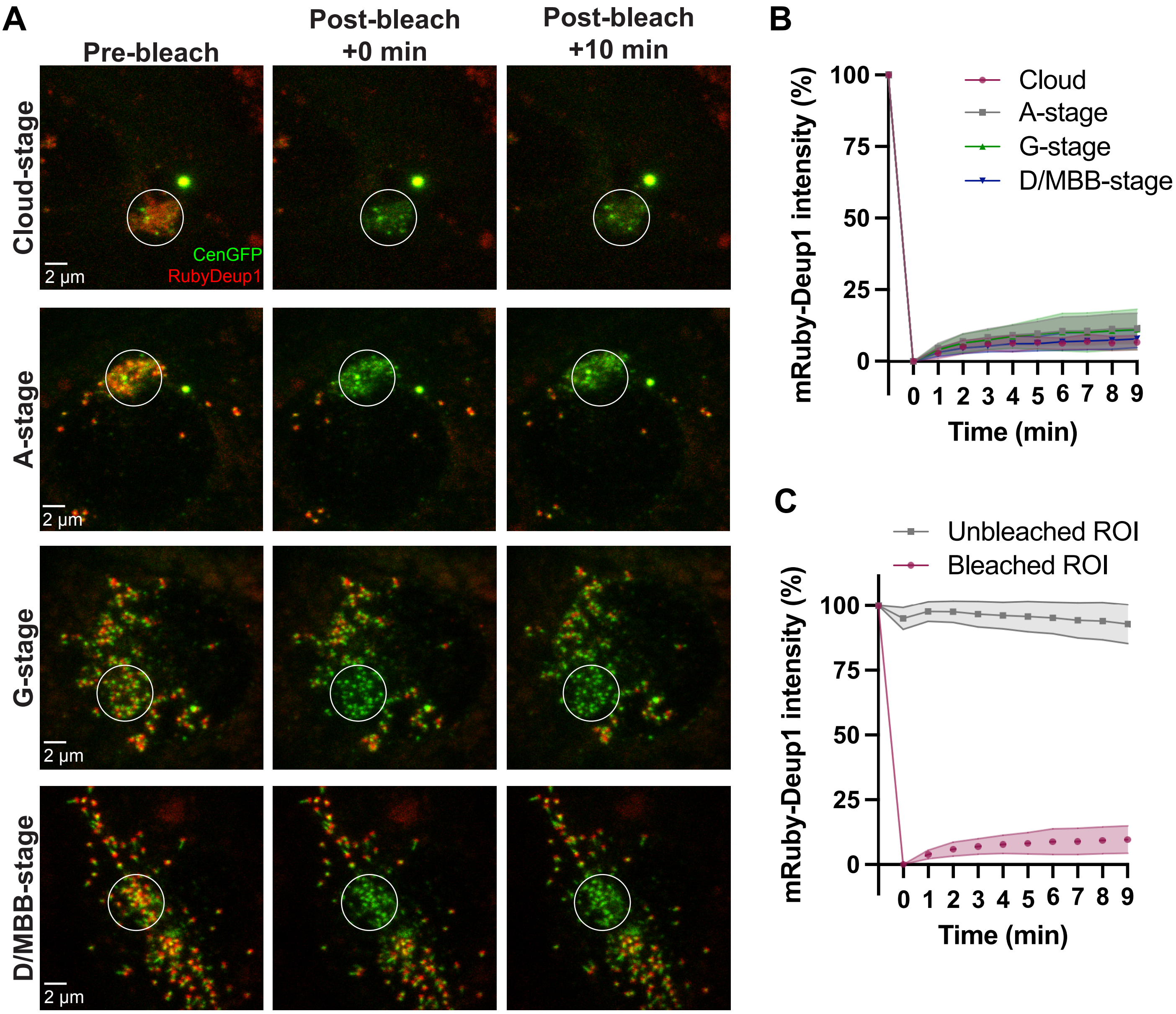

#### Figure 9 Supplementary 1

##### Maturation of procentrioles as regards to centrosomal centrioles during the MCC variant

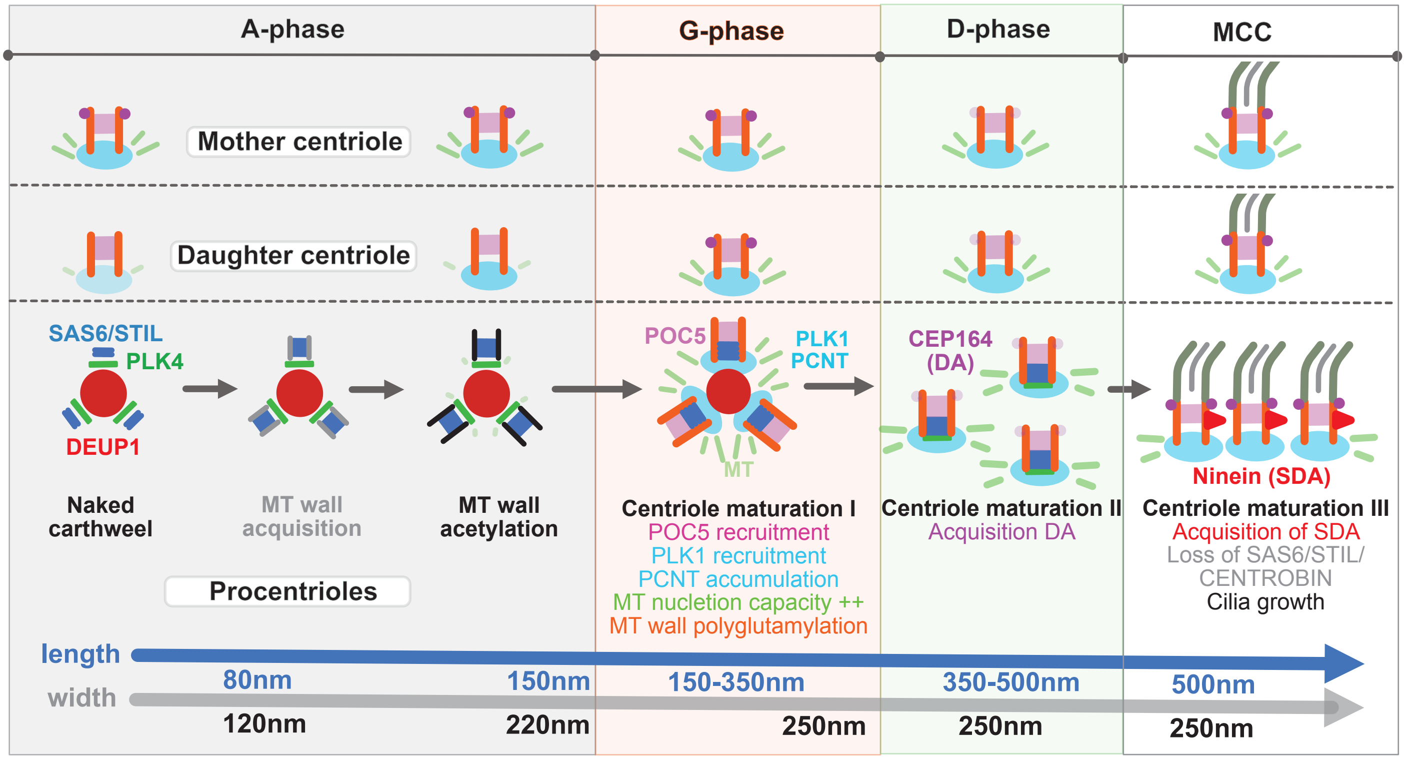
